# Heterogeneity-Preserving Discriminative Feature Selection for Disease-Specific Subtype Discovery

**DOI:** 10.1101/2023.05.14.540686

**Authors:** Abdur Rahman M. A. Basher, Caleb Hallinan, Kwonmoo Lee

## Abstract

The identification of disease-specific subtypes can provide valuable insights into disease progression and potential individualized therapies, important aspects of precision medicine given the complex nature of disease heterogeneity. The advent of high-throughput technologies has enabled the generation and analysis of various molecular data types, such as single-cell RNA-seq, proteomic, and imaging datasets, on a large scale. While these datasets offer opportunities for subtype discovery, they also pose challenges in finding subtype signatures due to their high dimensionality. Feature selection, a key step in the machine learning pipeline, involves selecting signatures that reduce feature size for more efficient downstream computational analysis. Although many existing methods focus on selecting features that differentiate known diseases or cell states, they often struggle to identify features that both preserve heterogeneity and reveal subtypes. To address this, we utilized deep metric learning-based feature embedding to explore the statistical properties of features crucial for preserving heterogeneity. Our analysis indicated that features with a notable difference in interquartile range (IQR) between classes hold important subtype information. Guided by this insight, we developed a statistical method called PHet (Preserving Heterogeneity), which employs iterative subsampling and differential analysis of IQR combined with Fisher’s method to identify a small set of features that preserve heterogeneity and enhance subtype clustering quality. Validation on public single-cell RNA-seq and microarray datasets demonstrated PHet’s ability to maintain sample heterogeneity while distinguishing known disease/cell states, with a tendency to outperform previous differential expression and outlier-based methods. Furthermore, an analysis of a single-cell RNA-seq dataset from mouse tracheal epithelial cells identified two distinct basal cell subtypes differentiating towards a luminal secretory phenotype using PHet-based features, demonstrating promising results in a real-data application. These results highlight PHet’s potential to enhance our understanding of disease mechanisms and cell differentiation, contributing significantly to the field of personalized medicine.

## 1 Introduction

Uncovering disease-specific subtypes within recognized cell types is crucial for understanding disease heterogeneity and responses to therapeutic treatments, as these subtypes provide detailed insights into the complexities of disease mechanisms ^1^. This area of study enhances our understanding of the various disease traits exhibited by different cells and patients. Systematic exploration of these subtypes across a spectrum of healthy and diseased conditions can contribute to advances in personalized and effective treatment approaches, which is particularly valuable given the diverse responses of cells and patients to treatments.

The discovery of disease-specific cell subtypes in various diseases began to emerge due to the capabilities of single-cell RNA-seq analysis ^2^. For instance, a previous study demonstrated that transcriptionally distinct sub-populations of major brain cell types are linked to the pathology of Alzheimer’s disease, involving myelination, inflammation, and neuron survival ^3^. Additionally, new pathological subtypes of epithelial cells and fibroblasts were recognized to be highly enriched in pulmonary fibrosis ^4^. Beyond cell subtypes, the impact extends to patient subtypes, where chromosomal translocations can give rise to different tumor types ^5,6^, thereby influencing the selection of optimal and personalized treatments ^7–9^. This underscores the significance of uncovering the diversity and variability of specific cell types, both within individual patients and across patient populations ^10–15^. Such insights are important for understanding the molecular mechanisms that initiate and progress diseases, improving patient classification, and aiding in the development of medical treatments that are more closely aligned with individual characteristics and needs. ^16–20^.

Molecular signatures, including genes, mRNA transcripts, and proteins, help delineate diverse cell types and states ^21^. The advancement of omics technologies over the past two decades has enabled the simultaneous analysis of thousands of molecular entities, facilitating the characterization of various cells and diseases ^22^. This progress has led to improved capabilities for describing complex biological conditions, which in turn demands more detailed classification of both cells and diseases. However, the complexity of omics data, which is inherently high dimensional, poses a challenge to downstream computational processes. Moreover, identifying the molecular signatures integral to data interpretation becomes a formidable task ^23^. To circumvent this, a computational process, called feature selection should be employed to extract a subset of features that are the most informative and pertinent from high-dimensional omics data to discern the specific molecular patterns distinctive to each cell or disease subtype. This process offers several benefits for omics data analysis, including reducing noise and redundancy, improving sparsity and interpretability, and enhancing computational efficiency. Feature selection is proven by various theoretical and empirical studies to be effective for different tasks, such as classification, regression, clustering, and dimensionality reduction ^24^. Nonetheless, conventional feature selection methods face limitations in the area of subtype identification, primarily due to the ensuing reasons.

Traditionally, discriminative feature selection techniques have been employed to examine the molecular patterns of cells or patient samples under predefined conditions. The prevalent methods for this approach fall within the area of differential feature expression analysis. This category of methods identifies molecules, such as genes, that display contrasting expression levels or associations with conditions across two distinct groups, such as cancer and healthy individuals ^25–41^. It is important to recognize that features from these methods are discriminative and differentially expressed (DE). However, in numerous real-world scenarios, the complexity and heterogeneity of omics data are not readily captured by differential expression methods. Hence, while the discriminative feature selection paradigm centers on distinguishing established conditions and states, it can inadvertently stifle the exploration of diversity inherent within the data. As demonstrated in Supplementary Fig. 1(k)-(l), the discriminative DE features have limited ability to separate AML and MLL subtypes. However, injecting heterogeneity by adding HV features can reveal these subtypes (Supplementary Fig. 1(m)-(n)). This limitation of discriminative DE features can potentially simplify feature space, ultimately impeding the effectiveness of in-depth subtyping ^42^.

To address this issue, Tomlins and colleagues ^43^ introduced a novel statistical method called “cancer outlier profile analysis” (COPA). This method was designed to identify subtypes within cancer patients by focusing on the genes whose expression profiles exhibit minor outliers. This pioneering work inspired a series of studies that improved COPA with more sophisticated statistical and machine learning techniques for subgroup discovery within each experimental condition ^44–49^. These methods involve ranking features based on statistical metrics that gauge the degree of abnormal expression across two experimental scenarios. Nonetheless, they are not well-suited for tackling complex subtyping challenges, such as deciphering the population structure from single cells and detecting the cell state transitions across conditions such as development processes. The limitation lies in the fact that outlier genes do not encompass the full spectrum of sample heterogeneity and often lack the requisite discriminative potency. Conversely, the concept of selecting highly variable (HV) genes gained prominence, particularly in the context of single-cell RNA-seq analysis, for uncovering molecular signatures linked to diverse cell types ^50–52^. Often, these methods seek a set of HV features that exhibit significant variation across samples, regardless of experimental conditions, employing these features for subsequent clustering analysis. The downstream differential analysis among these clusters can be conducted to establish associations with distinct molecular signatures. However, it is important to note that HV features are not intended for uncovering unknown subtypes within established conditions. Moreover, they may lack the necessary discriminatory power and often incorporate substantial irrelevant heterogeneity for specific subtyping tasks. This phenomenon may significantly compromise the capability of subtype discovery.

Given the constraints inherent in current methods, the process of disease-specific subtyping demands a substantial investment of time and resources. Researchers must integrate differential expression analysis, domain expertise, manual marker selection, and subsequent verification to achieve accurate results. For example, a computational framework was developed to integrate single-cell RNA-seq data with other genetic information obtained from genome-wide association studies (GWASs) to infer cell types, especially in cases where genetic variants impact diseases ^53^. However, the absence of a comprehensive computational framework adept at identifying subtypes solely from gene expression data in a supervised setting involving two distinct disease conditions remains a significant challenge. The algorithm should possess the capability to identify features that exhibit both differential expression and variability across latent subtypes. To overcome this obstacle, we introduce a novel category of features that has largely eluded the notice of existing methods, termed Heterogeneity-preserving Discriminative (HD) features. Through our deep metric learning approach, focused on gene embedding derived from single-cell RNA seq datasets, we discovered the existence of a significant proportion of HD features, effectively possessing both DE and differentially variable (DV) features characterized by the distinct IQR differences present between the two pre-defined experimental conditions. Capitalizing on these HD features, we can achieve a more refined clustering of patient samples or cells, facilitating a deeper comprehension of the factors that influence their heterogeneity. Therefore, this approach can reveal new molecular insights that might otherwise remain obscured, effectively transcending the challenges inherent in conventional methodologies.

To facilitate the identification of HD features, we developed a novel computational framework, named PHet (Preserving Heterogeneity), that leverages an iterative subsampling approach to determine an appropriate and adequate set of features for subtype discovery tasks from omics data. Striking a balance between heterogeneity and discriminativeness proves to be challenging, as these aspects often counteract each other. However, PHet is designed to find this balance, optimizing both aspects while keeping the number of selected features low. We evaluated the effectiveness of PHet by benchmarking it against six single-cell transcriptomics, eleven microarray, and two simulated datasets. PHet was compared to 24 computational methods using a variety of performance metrics to ensure a fair and comprehensive analysis. Our results suggest that PHet performs favorably compared to other methods in identifying subtypes under binary experimental conditions. Moreover, PHet has shown the capability to reveal novel cell subtypes in both mouse and human airway epithelium single-cell transcriptomics datasets ^54^. PHet also demonstrates consistency and robustness in handling high-dimensional data, distinguishing it from other algorithms in our study.

## 2 Results

### 2.1 Identification of heterogeneity-preserving discriminative features in omics data by deep metric learning

To better understand the key statistical attributes that contribute to the preservation of subtype heterogeneity within features, we conducted a feature statistic embedding through the utilization of deep metric learning (DML) ^55,56^ (see Section 4.2 in Methods). In the typical data format, rows and columns of input data represent measurements and features, respectively. For our analysis, this structure is transposed, with features as rows and measurements as columns, previously used for transforming non-image data into an image ^57^. We then sorted feature measurements within each disease or cell condition in ascending order to remove biases from the original observation order, ensuring that comparisons between diseases or cell conditions rely solely on the statistical properties of the features, not the arbitrary order of the data. Following this, we employed UMAP (Uniform Manifold Approximation and Projection) ^58^ or PCA (Principal Component Analysis) using the feature values, both in case and control groups to cluster features across disease or cell conditions based on the similarity of their respective statistical distributions. These clusters served as a foundation for calculating the triplet loss ^56^ in our DML approach, where positive or negative samples are within the same or different clusters as anchor samples, respectively. This triplet loss brings the embedding of similar features close together while simultaneously maximizing the distance between dissimilar features (Fig. 1(a)). Our DML encoder performed the task of embedding feature statistics from each condition into a lower-dimensional space, facilitated by the acquisition of a meaningful distance between feature embeddings. The final step of our process involved the calculation of the differences between the feature embeddings of the same feature under different conditions. This final subtraction step allows us to derive insights into the variations present across different disease states or cell conditions, thereby enhancing our understanding of the heterogeneity preservation of each feature.

Our implementation of DML based on UMAP-based clustering was applied to the Patel data ^59^, a single-cell RNA-seq dataset encompassing heterogeneous subtypes of primary glioblastomas. We computed the Euclidean distance between feature (i.e., gene) embeddings across two different conditions to measure the distances between genes based on their statistical properties. In the context of the Patel dataset, we employed DML to compare the embeddings of gene expression for two disease conditions—progenitor states (MGH26 and MGH30) and differentiated states (MGH28, MGH29, and MGH31). Upon observing the resulting embeddings, it becomes apparent that genes with high Euclidean distances are notably situated at a considerable distance from the center (as depicted in Fig. 1(b)). While these embeddings effectively illustrate the areas occupied by DE genes with substantial mean differences between the two conditions (Fig. 1(c)), they fall short of providing a comprehensive explanation for the acquired distance metrics. Intriguingly, mapping the IQR differences between the two conditions onto the embedding demonstrates a clear pattern: genes with similar IQR differences cluster closely together, and a significant proportion of genes characterized by large IQR differences between the two conditions tend to cluster within regions of substantial distance (Fig. 1(d)). Moreover, the DML applied based on PCA-based clustering revealed the same insights regarding the IQR differences, suggesting our finding does not reply on specific dimensional reduction techniques (Supplementary Fig. 1(a)-(h)). This demonstrates the potential of IQR differences as a valuable statistical attribute capable of reflecting the underlying heterogeneity between the two conditions.

Based on our observations, we introduce a novel concept termed Heterogeneity-preserving Discriminative (HD) features (Fig. 1(e)), which combines the properties of DE ^60^ and DV ^61^ traits. DE features are characterized by significant differences in expression or abundance across distinct groups, while DV features are determined based on variability between conditions. While previous studies have leveraged DV features for cancer studies to uncover variations within a group ^61–64^, and for cell type classification using a cascade of additional features for enhanced accuracy ^65^, the specific applications of DV features for subtyping within distinct conditions have not been extensively addressed in the literature. HD features constitute a distinct category, uniquely characterized by their combined representation of both mean expression differences and variability discrepancies, as determined by IQR differences between two conditions (Fig. 1(e)). This intersection of attributes brings to light a previously unexplored feature class with potential implications.

To demonstrate the utility of HD features, we performed a comparative subtyping analysis using DE (identified through mean differences), DV (computed using IQR differences), and the novel HD features within the context of the Patel dataset ^59^ (Fig. 1(k)-(n)). We also included HV features identified using dispersion-based selection, as proposed by Satija and colleagues ^51^, capturing inherent data heterogeneity without considering predefined conditions. We applied the k-means algorithm to cluster the data based on the four different feature types: DE, HV, DV (IQR Diff.), and HD (Fig. 1(g)-(j)). Subsequently, we assessed the clustering performances using adjusted Rand index (ARI) and V-measure metrics on the reduced feature space following feature selection, without applying any additional dimensional reduction techniques such as UMAP or PCA. Our findings revealed that DE features demonstrated weak clustering performance, with an ARI of 36.44% and a V-measure of 48.59%, similar to HV features (Fig. 1(f)). This suggests that HV and DE features, which consisted of 1529 and 1486 features, respectively, were insufficient for accurately characterizing the subtypes. However, DV features based on IQR differences reduced the feature size to 904 and showed modest improvement in clustering over DE and HV features. The most promising outcomes were obtained when using the HD features, which comprise less than 15% (641 features) of the total features in the Patel data, achieving an ARI of 82.67% and a V-measure of 79.36%. When we visualized the cluster results using ordered distance maps, HD features exhibited clear separation of five clusters as opposed to the other feature sets (Fig. 1(j)), underscoring the effectiveness of HD features in capturing sample heterogeneity within the dataset. These HD features combine the properties of both DE and DV, offering a valuable resource for gaining a more refined understanding of the factors influencing disease states. These results motivated the development of PHet, our computational framework aimed at optimizing the integration of DE and DV, facilitating effective subtyping in omics data under two experimental conditions. The emergence of HD features and their incorporation into our methodology represents a promising step toward enhancing approaches for disease subtyping.

### 2.2 Overview of PHet

PHet is a method designed to detect informative features for subtype discovery from omics expression data. We employed iterative subsampling to enhance PHet’s capability to handle sample heterogeneity, based on Fisher’s method ^66^, which was previously used in DECO ^49^. The initial step involves annotating the data with two distinct conditions: control and case (e.g., healthy and cancerous conditions). Subsequently, the data undergoes a series of preprocessing steps, including removing low-quality samples and features and normalizing expression values to have zero mean and unit variance. This preprocessed data is then fed to the PHet pipeline which is composed of six major stages (Fig. 1(o) and Section 4.3). (1) Iterative subsampling ^66–68^: This step calculates both p-values, using t-test or z-test, and mean absolute interquartile range differences (ΔIQR) for each feature between two randomly chosen subsets of control and case samples. The size of these subsets is determined by the closest integer to the square root of the minimum number of samples in either the control or case group, i.e., min(*n, m*), where *n* and *m* correspond to the number of samples in control and case groups, respectively. This approach ensures even subsample sizes of both groups and has been applied to detect features containing intrinsic heterogeneity^49^ and rare cell subtypes, such as ionocyte cells which represent only 1 − 2% of airway epithelial cells ^54,69^. The p-values measure the statistical significance of the difference in expression levels between the two groups, while the ΔIQR values indicate the differences in expression variability between them. To capture sufficient features that help in subtype identification, the subsampling procedure is repeated for a predefined number of iterations (default is 1000). (2) Fisher’s combined probability test: Following subsampling, the collected p-values are summarized by the Fisher’s combined probability test. The results from this test serve as prior information to calculating feature statistics and ranking. (3) Enhancing discriminatory power of features: While iterative subsampling with Fisher’s score effectively detects mean differences in heterogeneous samples, its ability to discriminate can be limited when sample distributions deviate from unimodal patterns (Supplementary Fig. 1(o)). To address this issue, we have incorporated the nonparametric Kolmogorov-Smirnov (KS) test, which assesses differences between the cumulative distribution functions of control and case samples. This method helps identify features that significantly contribute to observed variations between the groups. Consequently, we enhance the discriminative power of Fisher’s scores by leveraging the results from the KS test. The p-values resulting from the KS test are grouped into pre-defined bins, each with uniform width (default is four bins with intervals defined as [0, 0.25], (0.25, 0.5], (0.5, 0.75], and (0.75, 1.0]). Each bin is assigned a specific weight based on the features present within that bin. These weights reflect the discriminatory power of respective features, allowing the regularization of the scores obtained from Fisher’s method. Importantly, these weights (default is (0.4, 0.3, 0.2, 0.1)) are empirically determined and may vary based on the specific analysis conducted. (4) Feature statistics and thresholding: Fisher’s scores (**f**) multiplied by the features discriminatory power (**o**) are then combined with the absolute mean values of ΔIQR (**r**) to estimate feature statistics, i.e., **r** + (**f** ⊙ **o**), where ⊙ corresponds to the Hadamard (element-wise) product. To ensure that the values of **r** and **f** ⊙ **o** are on the same scale, standardization is applied. This involves dividing the values of **r** by the sum of its values, and similarly, dividing **f** ⊙ **o** by the sum of its values. (5) Feature significance: the feature statistics are fitted using the gamma distribution, and features exceeding a user threshold (*α*) are trimmed. By default, *α* is set to 0.01. (6) Downstream analysis: In the final step, the selected features are used for various downstream tasks, such as clustering to reveal the heterogeneity within the dataset.

**Figure 1.**
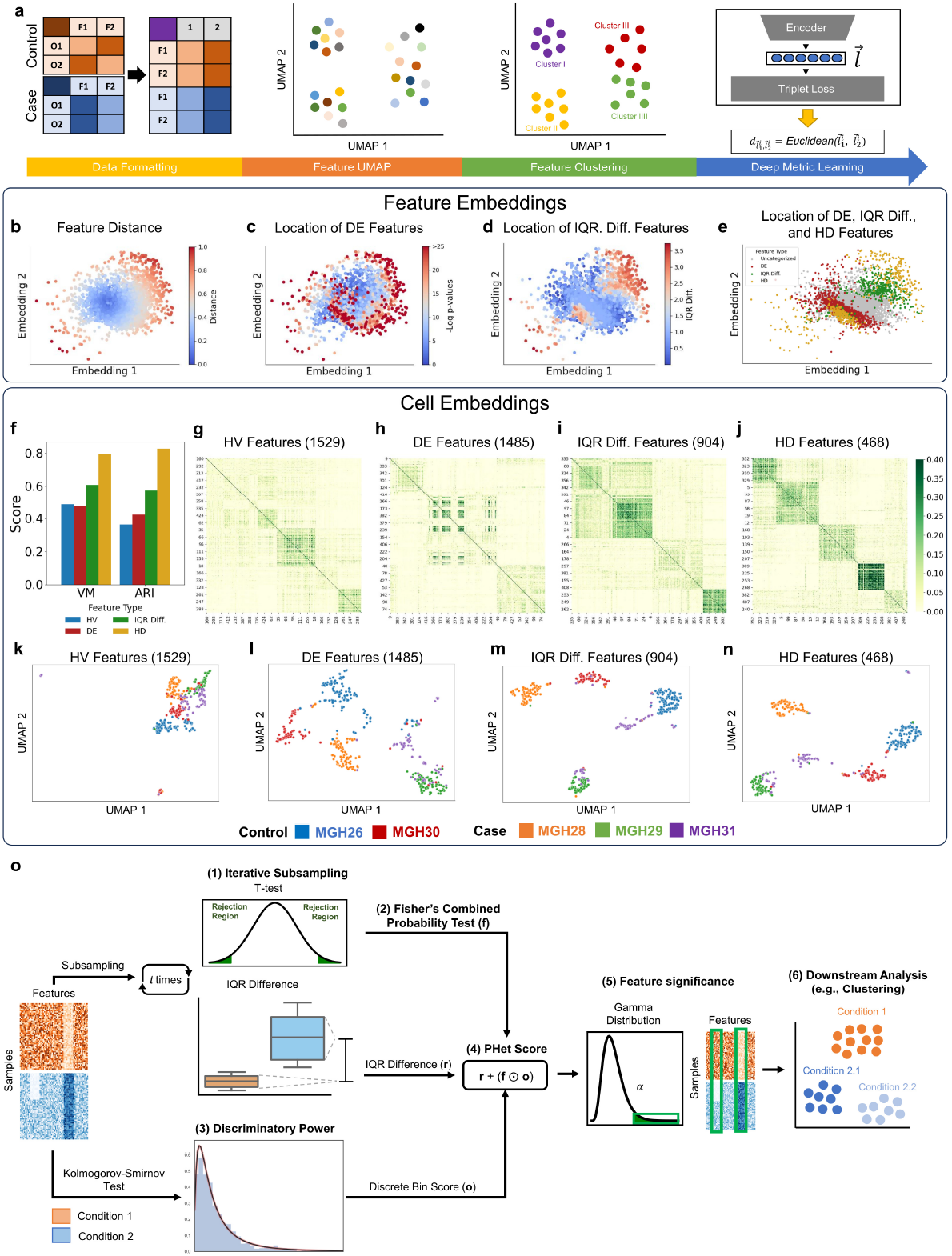
Characterization of features extracted by deep-metric learning (DML) and the overview of PHet framework. **a**, A schematic of DML approach with triplet loss, which is used to analyze the feature space. In this method, a UMAP is generated, where each point represents a feature from a specific condition. Clustering is then applied to the UMAP space. The resulting clusters and the original data are used as input for the triplet loss in DML. The embeddings from the encoder are used to calculate the Euclidean distance between the same feature of different classes, where 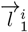 and 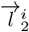 are the embeddings for the gene *i* in case and control conditions, respectively, and 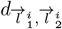 is the Euclidean distance between these embeddings. **b**,**c**,**d**,**e**, Scatter plots of feature embeddings using the Patel data with color representing the distance between the same feature of different conditions (**b**), logged p-values from mean differences (using z-test) between conditions of features (**c**), IQR difference between conditions (**d**), and location of respective feature types on the plot (**e**). **f**, Clustering performance results on the Patel data using the adjusted Rand index (ARI) and V-measure (VM) metrics based on DE, HV, IQR Diff., and HD features. **g**,**h**,**i**,**j** Heat maps of clustering results on the Patel data based on DE, HV, IQR Diff., and HD features, respectively. **k**,**l**,**m**,**n**, UMAP visualizations of 1529 HV features(**k**) (based on a dispersion threshold of *>* 0.5), 1485 DE features (**l**) (based on the z-test at significance level of 0.01), 904 ΔIQR features (**m**) (based on a threshold of *>* 0.4), and 468 HD features (**n**) that are intersection of both DE and ΔIQR features. **o**, PHet pipeline is composed of six major steps: (1)-an iterative subsampling to calculate p-values, using t-test or z-test, and maximum absolute interquartile range differences (ΔIQR) for each feature between two subsets of control and case samples, (2)-the Fisher’s combined probability test to summarize the collected p-values, (3)-the Kolmogorov-Smirnov test to adjust the Fisher’s scores. (4)-feature statistics estimation using a combination of the ΔIQR values, the Fisher’s combined probability scores, and the weighted features representing their discriminatory power, (5)-fitting feature statistics using the gamma distribution, and features exceeding a user threshold are trimmed (*<* 0.01), and (6)-downstream analysis, such as clustering analysis on the reduced omics data to reconstruct data heterogeneity.

The key hyperparameters in PHet consist of the binning weights **w** and the user threshold *α*. Through an extensive analysis using the ARI metric on separate test datasets (Supplementary Tables 1 and 2), we determined that setting *α* to 0.01 and **w** as (0.4, 0.3, 0.2, 0.1) led to empirically optimal clustering outcomes with an average ARI score of 61.02% across all test datasets (Supplementary Fig. 36). Furthermore, this configuration also resulted to a reduction in the number of features, with an average of 395.1 features retained across the test datasets (Supplementary Fig. 36). Therefore, these specific settings serve as the default hyperparameters for PHet throughout the entirety of this manuscript. For a more comprehensive understanding of these hyperparameters tuning process for PHet, please refer to Supplementary Note 1.

### 2.3 Evaluation of PHet’s performance in identifying subtypes of single cells and patients

To assess the effectiveness of PHet in subtyping tasks, we benchmarked PHet against six publicly available single-cell transcriptomic (scRNA-seq) datasets and eleven well-known microarray gene expression datasets (refer to Section 4.13; Supplementary Tables 4-5). We categorized cells in the scRNA-seq datasets and patient samples in the microarray data into two experimental conditions: control and case. An effective algorithm should accurately identify cell/patient subtypes while preserving biological integrity and heterogeneity with a small set of features. Prior to algorithm execution, we conducted clustering analysis on each scRNA-seq dataset using all features. The results revealed that cells in the Baron ^70^ dataset exhibited little sign of distinct clusters, with a low ARI score of 5.66% (Supplementary Fig. 20). Additionally, the Darmanis ^71^, and Patel ^59^ datasets showed moderate ARI scores of 25.88% and 19.72%, respectively, suggesting the presence of cell subclusters that partially aligned with true subclasses (Supplementary Figs 22 and 24). Conversely, the Camp ^72^, Li ^73^, and Yan ^74^ scRNA-seq data already displayed discernible cell subpopulations using all the features (Supplementary Figs 21, 23, and 25, respectively) with above average ARI scores of 53.22%, 58.07%, and 58.04%, respectively, allowing for evaluation of subtype detection methods to select a minimal meaningful feature set to retain the true cell heterogeneity. Similar analyses were performed on the microarray expression datasets, revealing that all datasets were composed of admixed patient subtypes (Supplementary Figs 9-19).

In our analysis of both types of datasets, we performed a comprehensive comparison of PHet and PHet (ΔDispersion) against 11 well-established methods as well as eight additional PHet variants introduced in this work (Section 4.4 and Table 1) for cell/patient subtypes identification across multiple performance metrics including F1 score for identifying DE features, ARI, adjusted mutual information, homogeneity, completeness, and V-measure for subtype discovery. The pre-annotated types of cells and patients in the benchmark datasets are considered the ground truth labels to quantify ARI (Supplementary Tables 4-5), and the top 100 DE features from LIMMA were used as the ground truth DE features to calculate the F1 scores of the algorithms (see Section 4.6 for details on metrics), because LIMMA is considered a standard benchmark for comparing DE features identified by each method (see Section 4.4 for LIMMA). This comparative analysis allows us to assess the extent of dissimilarity between the discriminatory features identified by each method and those identified by LIMMA. Moreover, this measurement aids in identifying the specific subset of DE features that facilitate the clustering of samples into two primary categories. The evaluated methods included four DE feature analysis tools and their variants: the standard Student t-test ^75^, Wilcoxon rank-sum test ^76^, Kolmogorov–Smirnov test ^77^, and LIMMA ^78,79^. Additionally, we evaluated the variants of PHet that utilize methods based on dispersion ^51^ and IQR ^80^. Furthermore, seven outlier detection algorithms were included in the evaluation: COPA ^81^, OS ^44^, ORT^45^, MOST ^46^, LSOSS ^47^, DIDS ^48^, and DECO ^49^. More elaboration about these methods are provided in Section 4.4 and Table 1.

**Table 1:** Methods used in the benchmark. MAD: median absolute deviation of expression values; RDA: recurrent differential analysis; and NSCA: non-symmetrical correspondence analysis.

| Method | Strategy | Down-regulated features | Outlier weighting | Subtype discovery | Multiple groups | Reference |
| --- | --- | --- | --- | --- | --- | --- |
| t-statistic | Mean difference | ✓ | × | × | × | 75 |
| t-statistic+Gamma | Mean difference and the gamma distribution | ✓ | × | × | × | 75 and this work |
| Wilcoxon | Sum of the ranks | ✓ | × | × | × | 76 |
| Wilcoxon+Gamma | Sum of the ranks and the gamma distribution | ✓ | × | × | × | 76 and this work |
| KS | Empirical cumulative distribution function | ✓ | × | × | × | 77 |
| KS+Gamma | Empirical cumulative distribution function and the gamma distribution | ✓ | × | × | × | 77 and this work |
| LIMMA | Feature-wise linear models and empirical Bayes to obtain posterior variance estimators | ✓ | × | × | ✓ | 78 |
| LIMMA+Gamma | Feature-wise linear models, empirical Bayes to obtain posterior variance estimators, and the gamma distribution | ✓ | × | × | ✓ | 78 and this work |
| COPA | Percentile MAD | × | × | × | × | 43 |
| OS | Quantile ordered & cut-off | ✓ | × | × | × | 44 |
| ORT | Robust t-statistic | × | × | × | × | 45 |
| MOST | Maximum ordered | ✓ | × | × | × | 46 |
| LSOSS | Least sum of ordered subset square t-statistic | ✓ | × | × | × | 47 |
| DIDS | Maximum value from control group | ✓ | ✓ | × | × | 48 |
| DECO | RDA and NSCA | ✓ | ✓ | ✓ | ✓ | 49 |
| Dispersion (composite) | Normalized dispersion of all samples | — | — | ✓ | — | 51 |
| Dispersion (per condition) | Normalized dispersion of samples based on each condition | — | — | ✓ | ✓ | 51 and this work |
| ΔDispersion | Differences of normalized dispersion between samples of each condition | — | — | ✓ | ✓ | 51 and this work |
| ΔDispersion+ΔMean | Differences of normalized dispersion and mean between samples of each condition | — | — | ✓ | ✓ | 51 and this work |
| IQR (composite) | Z-score and IQR of all samples | — | — | ✓ | — | 80 and this work |
| IQR (per condition) | Z-score and IQR of samples based on each condition | — | — | ✓ | ✓ | 80 and this work |
| ΔIQR | Z-score and ΔIQR | — | — | ✓ | ✓ | 80 and this work |
| ΔIQR+ΔMean | Z-score, ΔIQR, and ΔMean | — | — | ✓ | ✓ | 80 and this work |
| PHet (ΔDispersion) | Log transform, iterative subsampling, Fisher's method, and ΔDispersion | ✓ | ✓ | ✓ | ✓ | This work |
| PHet | Z-score, iterative subsampling, Fisher's method, and ΔIQR | ✓ | ✓ | ✓ | ✓ | This work |

The results showed that PHet consistently outperformed other baseline methods, achieving an average ARI score above 65.72% for subtype detection (Fig. 2(c)) while maintaining a competitive F1 score for identifying DE features (Fig. 2(a)) and selecting a smaller number of features (less than 300 on average; Fig. 2(b); see Supplementary Fig. 4 for additional results using multiple performance metrics). Statistical significance was assessed through a paired, two-tailed t-test, after obtaining no indication of non-normality in the data (D’Agostino’s *K*^2 82^ p-value: 0.498). This analysis demonstrated that PHet’s ARI scores (65.7%) are higher than those of the most competitive DE-based methods, KS+Gamma (54.32%, p-value of 0.023, Cohen’s *d* effect size of 0.57), LIMMA (50.55%, p-value of 0.007, Cohen’s *d* effect size of 0.69), and the most competitive outlier-based method, DECO (47.04%, p-value of 0.001, Cohen’s *d* effect size of 0.91) (see Supplementary Fig. 2(a)-(b)). PHet’s average F1 score (74.10%) was also higher than KS+Gamma (49.16%, p-value of 0.001, Cohen’s *d* effect size 1.14) and DECO (47.13%, p-value less than 0.001, Cohen’s *d* effect size of 1.1) (see Supplementary Fig. 2(c)-(d)). The comparison of F1 with LIMMA was not conducted because LIMMA’s features were regarded as ground truth DE features. While the average number of selected features of PHet (294.59) is less than those of KS+Gamma (793.71), it was much smaller than those for LIMMA (3783.59, p-value of 0.001, Cohen’s *d* effect size of 1.310) and DECO (3303.00, p-value of 0.002, Cohen’s *d* effect size of 1.218) (see Supplementary Fig. 2(e)-(f)).

**Figure 2.**
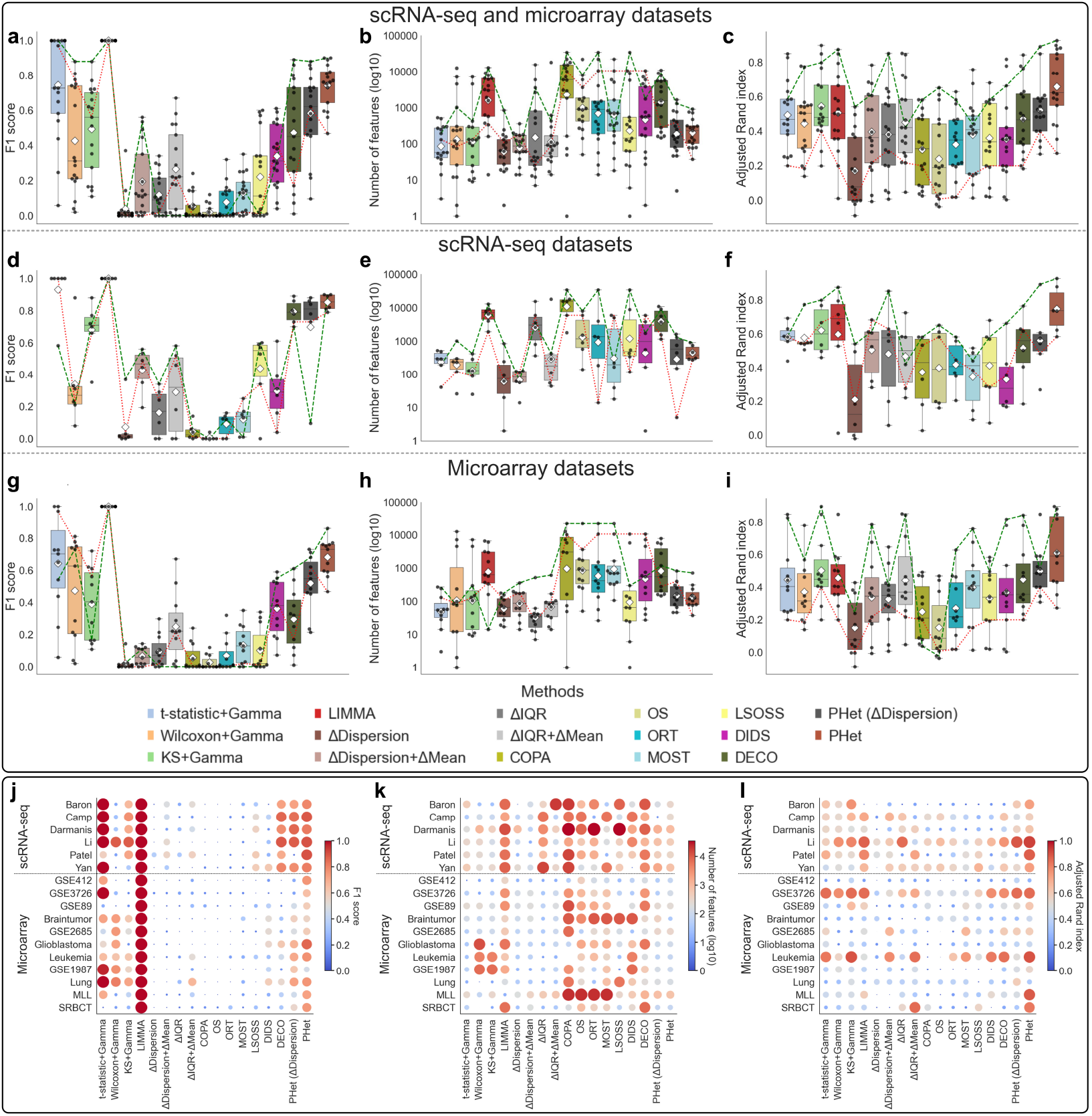
Performance comparison of PHet against 16 baseline methods across 6 single-cell transcriptomics and 11 microarray datasets. **a**,**d**,**g**, F1 scores of each method for detecting the top 100 DE features that are obtained using LIMMA for both microarray and scRNA-seq (**a**), six single-cell transcriptomics (**d**), and 11 microarray (**g**) datasets. **b**,**e**,**h**, The number of selected features by each method using both microarray and scRNA-seq (**b**), six single-cell transcriptomics (**e**), and 11 microarray (**h**) datasets. **c**,**f**,**i**, The adjusted Rand index of each method for both microarray and scRNA-seq (**c**), six single-cell transcriptomics (**f**), and 11 microarray (**i**) datasets (Supplementary Tables 2 and 3). The box plots show the medians (centerlines), first and third quartiles (bounds of boxes), and 1.5 × interquartile range (whiskers). A ◊ symbol represents a mean value. A green dashed line indicates the best-performing result on a dataset of PHet on each metric, while a red dashed line represents the worst-performing result on a dataset of PHet on each metric. **j**,**k**,**l**, Dot plots of F1 scores (**j**), number of selected features (**k**), and adjusted Rand index scores (**l**) are presented for each method applied to both microarray and single cell transcriptomics datasets.

Further investigation of PHet’s performance on an individual dataset basis revealed its consistency and robustness in terms of average ARI scores, particularly for the scRNA-seq datasets (Fig. 2(j)-(l)). While PHet is not the only method capable of detecting meaningful subtypes, it consistently exhibited high ARI and F1 values across all six scRNA-seq datasets (Fig. 2(j) and (l)). In contrast, other baseline methods performed well only on subsets of the scRNA-seq datasets. Even the competitive baseline methods did not demonstrate acceptable ARI performance with two or three datasets (KS+Gamma with Darmanis, Patel, and Yan; LIMMA with Baron and Darmanis; DECO with Baron, Darmanis, and Patel). Although PHet exhibited robust performance with scRNA-seq datasets, it performed poorly in detecting subtypes in four microarray datasets, including GSE412, Braintumor, Glioblastoma, and Lung. Similarly, none of the baseline methods were able to sufficiently identify subtypes in these datasets. This may be attributed to the limited information available in these microarray datasets for subtype discovery. When PHet successfully detected subtypes in the microarray datasets, some other baseline methods also detected subtypes. However, their performances were still inconsistent and highly specific to individual datasets, similar to the scRNA-seq data case.

Due to differences in sample numbers and the levels of heterogeneity, we performed quantitative comparisons among the methods with scRNA-seq and microarray datasets, separately. For scRNA-seq datasets (Supplementary Table 5), we observed that PHet outperformed established outliers detection methods (e.g., DECO) across all scRNA-seq datasets with a mean F1 score exceeding 85% (Fig. 2(d)) with respect to discriminative performance. Notably, PHet was able to strike a balance between keeping the feature number low with an average of fewer than 450 features and maximizing the discriminative performance for all single-cell transcriptomic datasets (Fig. 2(e)). Since the DE features were derived from LIMMA, the average F1 score for LIMMA is 1. The results obtained from a basic t-statistic method were found to be consistent with those obtained from LIMMA, which is expected given that LIMMA’s approach is similar to the t-statistic but incorporates different variance calculation and advanced functionalities. As a result, there is a high level of agreement in the top 100 differentially expressed features identified by these two methods. By fitting feature statistics using the gamma distribution, most DE-based methods struggled to match PHet’s performance.

In terms of average ARI, which measures the agreement between ground truth and clustering results, PHet outperformed existing methods across six scRNA-seq datasets and achieved *>* 10% and *>* 14% gain over the competitive performing algorithm, KS+Gamma and LIMMA, respectively (Fig. 2(f)), and over 15% and 20% increase from PHet (ΔDispersion) and DECO, respectively. This suggests that the IQR-based approach is more effective than dispersion-based feature selection, leading to improved clustering quality and better recovery of cell types. The KS test is useful for detecting DE features because this test does not require any assumptions about the shape or parameters of data distributions. Instead, the KS test compares the cumulative distributions of features, which makes it sensitive to any changes in the data ^83^. This sensitivity allows the KS to effectively detect variations in expression levels that other tests (e.g., t-statistic and Wilcoxon rank-sum test) may not be able to capture ^65^. PHet leverages the benefits from both ΔIQR and KS, leading to better clustering results as indicated by multiple performance metrics, such as adjusted mutual information, homogeneity, completeness, and V-measure, across six scRNA-seq datasets (Supplementary Fig. 6).

To visualize the effectiveness of the PHet’s feature selection, we compared pairwise similarity heatmaps using the selected features from each method and annotated cell types in the scRNA-seq datasets against other competing methods (KS+Gamma, LIMMA, DECO). Notably, among the 14 cell types in the Baron dataset (Supplementary Table 5), PHet selected 748 features that contribute to at least 7 clusters of cells (Fig. 3(a)). Moreover, in this dataset, four cell types—alpha, beta, gamma, and delta—representing the endocrine cells, exhibit inherent hierarchical relationships due to their shared expression profiles ^70^(Fig. 3(a)). Specifically, the alpha and gamma cells were observed to form a closely related group. Furthermore, delta cells were positioned as a group connected with the alpha/gamma cell group in the subsequent hierarchy, followed by the integration of beta cells into this configuration. PHet’s similarity heatmap preserved this hierarchical information (Supplementary 20). In contrast, the three baseline methods lost it entirely, even if KS+Gamma produced a higher ARI value (Fig. 3(b)-(c)). In the Camp dataset, while PHet achieved a lower ARI value, it also preserved the inter-cluster distances with much fewer number of features than LIMMA and DECO (Fig. 3(e)-(h)). In the other four scRNA-seq datasets where PHet achieved the highest ARI values, PHet produced more distinct clusters and better retained the hierarchy of cell types compared to the three baseline methods (Fig. 3(i)-(x)). This suggests that PHet can not only identify cell subtypes but also excels at revealing their hierarchical structures.

**Figure 3.**
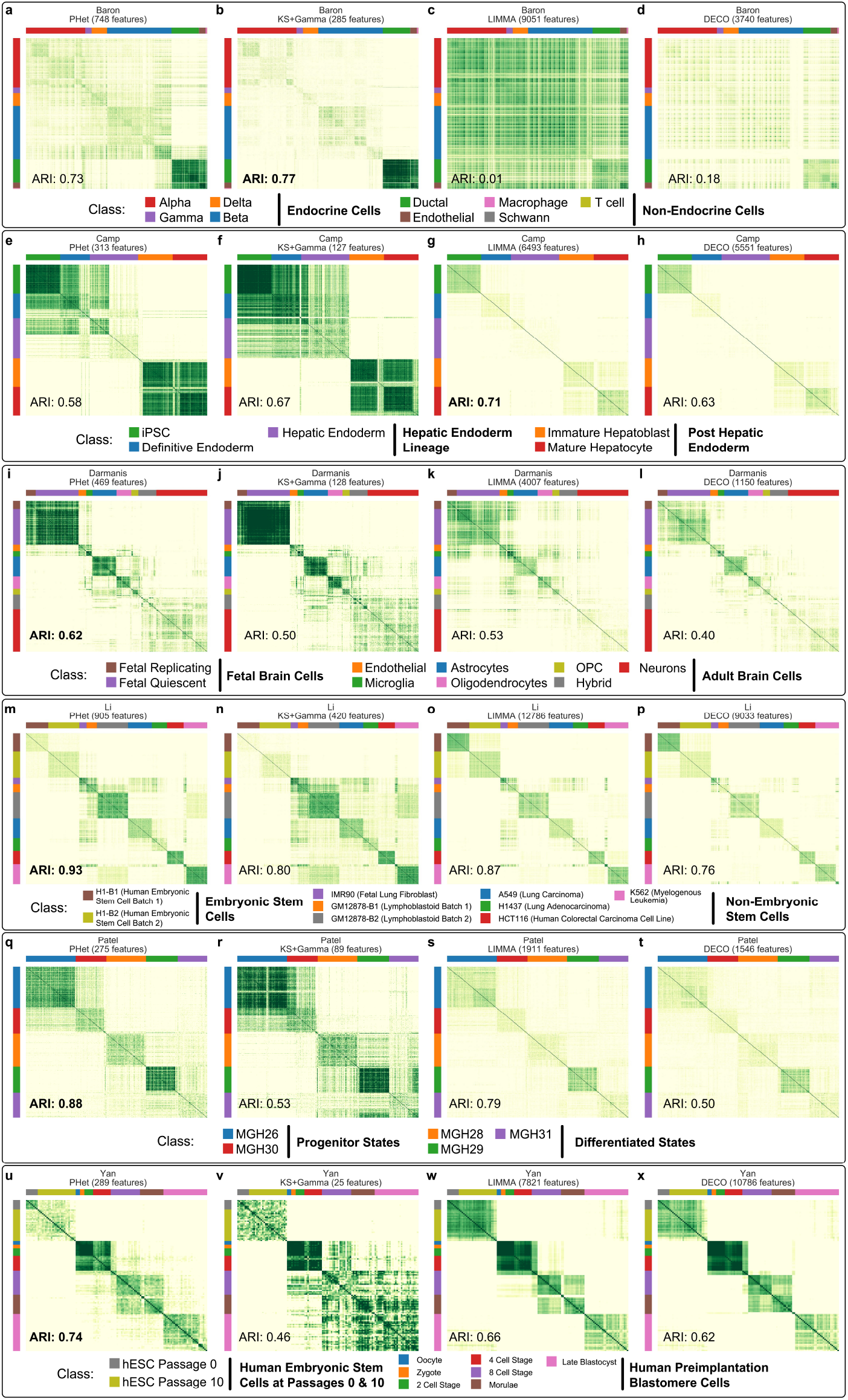
Heatmaps displaying the clustering results of PHet and the top three methods are presented for six single cell transcriptomics datasets. The datasets are: Baron (**a**-**d**), Camp (**e**-**h**), Darmanis (**i**-**l**), Li (**m**-**p**), Patel (**q**-**t**), and Yan (**u**-**x**). For each method, the selected features are used. The bold font ARI score indicates the best performing method for the corresponding data in the comparison.

In the microarray gene expression datasets (Section 4.13; Supplementary Table 4), which aim to uncover patient subtypes, the performance of all algorithms was generally lower in terms of F1 and ARI scores compared to the scRNA-seq datasets. Additionally, the similarity heatmaps exhibited less distinct cluster structures and hierarchical information than those in the scRNA-seq dataset (Supplementary Figs 7-8). This is likely due to the limited heterogeneity and sample numbers within these datasets. Nonetheless, in terms of clustering, we observed that PHet outperformed other baseline methods, achieving an average ARI score exceeding 60% while selecting fewer features (less than 130 on average; Fig. 2(g)-(i)). For example, in the analysis of the MLL dataset ^84^ (Supplementary Table 4), 72 patient samples were examined and categorized into three types of leukemia: acute lymphoblastic leukemia (ALL), mixed-lineage leukemia (MLL), and acute myeloid leukemia (AML). The ALL patients were considered the control group, while the MLL and AML patients were categorized as the case group. PHet recognized three clusters of patients with an ARI score of 88% that closely matched their true sample types (Fig. 2(l)). In contrast, none of the competing algorithms, including LIMMA, were unable to clearly identify these three patient subtypes (Supplementary Fig. 8(n)-(p)). For the Leukemia ^20^ dataset, KS+Gamma and DECO displayed three distinct clusters on par with PHet (Supplementary Fig. 8(a)-(d)) with similar ARI scores in the range of 84 ∼ 90%. In the evaluation of ten other microarray datasets, different algorithms displayed varying levels of effectiveness in identifying sample heterogeneity and subtypes. For example, KS+Gamma exhibited strong performances in the GSE89^85^ dataset, as evidenced by average ARI scores of 65.48%, while PHet demonstrated slightly weaker performance with average ARI scores of 61.38% (Supplementary Fig. 7(i)-(j)). DECO, an outlier detection method, performed remarkably well in the GSE2685^86^ dataset with an average ARI score of 60%, marginally surpassing PHet’s 59% (Supplementary Fig. 7(q)-(t)). Across the other benchmark microarray datasets, PHet consistently demonstrated competitive or superior results across various metrics (Supplementary Figs 7 and 8). Despite PHet’s slight underperformance in certain specific datasets, it is important to note that other baseline methods exhibited varying results across all scRNA-seq and microarray datasets, whereas PHet consistently demonstrated consistent performance across various metrics, such as F1 and the number of predicted features.

### 2.4 Ablation studies of PHet’s components

To further explore the impact of PHet’s components on feature selection for subtype detection, we conducted ablation studies on the same microarray and single-cell transcriptomics datasets discussed in Section 2.3. We systematically examined the impact of removing and reintegrating three main components of PHet while keeping *α* and **w** at their optimal values: Fisher’s scores, the absolute mean values of ΔIQR, and feature discriminatory power (Section 2.2). The integration of the first two components necessitates PHet to employ iterative subsampling, while the last component does not entail the subsampling process. The outcomes of these experiments revealed that relying solely on ΔIQR (PHet(+ΔIQR,-Fisher,-Discriminatory)), while disabling the other two components, led to suboptimal ARI scores for microarray and (Supplementary Fig. 37). This suggests that the exclusive reliance on ΔIQR limits the discriminatory power in extracting DE features, thereby impeding the accurate delineation of subtypes within conditions. Conversely, when exclusively utilizing Fisher’s method (PHet(-ΔIQR,+Fisher,-Discriminatory)), notable improvements were observed, with gains over 10% and 30% average ARI scores on microarray and scRNA-seq datasets. These improvements can be attributed to the sub-sampling process, which effectively captured variations in condition and case samples. Importantly, when all three components were incorporated, i.e., PHet(+ΔIQR,+Fisher,+Discriminatory) or PHet, the most favorable outcomes were achieved, with average ARI scores exceeding 60% and 75% on microarray and scRNA datasets, respectively. These results demonstrate statistical significance based on ARI scores, supported by a paired two-tailed t-test with a p-value of 0.026 when compared to the second-best method, PHet (+ΔIQR,+Fisher,-Discriminatory). These findings underscore the significance of feature scores derived from discriminatory and ΔIQR components, highlighting their valuable contribution to subtype detection.

We also conducted test experiments to assess the importance of the iterative subsampling component of PHet on clustering outcomes by excluding it while maintaining the same hyperparameters. The findings demonstrated that the exclusion of iterative subsampling resulted in suboptimal performance (with a p-value of 0.0004 using a paired two-tailed t-test), with an average ARI score of 48.78%, in comparison to the standard configuration of PHet, which achieved an average ARI score of 65.72% (Supplementary Fig. 38). These results underscore the significant role of iterative subsampling in subtype detection.

### 2.5 Evaluation of PHet’s discriminative performance on simulated data

While PHet exhibited competitive discriminative performance in our previous benchmark test, the evaluation was confined to comparing the selected features with LIMMA-based DE features. To bolster the validation of PHet’s ability to capture features that can discriminate the two conditions while also retaining a small set of these features, we conducted a comprehensive evaluation of 25 algorithms (outlined in Section 4.4 and Table 1) using two sets of simulated data where ground truth of DE features is available. These datasets correspond to the ‘minority’ and ‘mixed’ model schemes, as proposed by Campos and colleagues ^49^, and are also designed to capture sample heterogeneity under the supervised settings. In the ‘minority’ model, a small fraction of case samples exhibited changes in specific features, while the ‘mixed’ model displayed substantial intra-group variation in both case and control samples for those features. Each dataset consists of 40 samples, evenly split between control and case groups with 1100 features, including 93 DE features between the two groups. We generated 5 datasets based on the minority model, varying the proportions of perturbed samples in the case group from 5% (1 in 20) to 45% (9 in 20). Those perturbed samples were introduced by modifying the expressions of 100 randomly selected features, which encompassed DE features. Similarly, we constructed five datasets following the same procedure for the mixed model, with perturbed samples evenly distributed between a subset of both case and control groups. Further details about these datasets are provided in Section 4.13. For quantitative analysis, the F1 score was used to compare the top 100 predicted features of each method with the top 100 true features. Furthermore, we recorded the number of informative features with high scores (at *α <* 0.01; see Section 4.4) for each method. We followed the model-specific parameters recommended by the respective authors of each method, thus avoiding any bias or error stemming from inappropriate parameter choices. A best performing algorithm should attain high F1 scores across both model schemes, while also exhibiting the capability to predict a small set of important features that contribute to the perturbed samples. Both PHet and PHet (ΔDispersion) retained a fewer set of informative features (at *α <* 0.01) for both minority and mixed changes under all settings (Fig. 4(c) and (d); Supplementary Fig. 3(c) and (d)), indicating that both methods have a robust ability to detect important features, contributing to perturbed samples. The unsupervised Dispersion (composite) method performed on par with PHet (F1 score of 0.92 on average) for both model schemes, achieving an F1 score of 0.88 on average. However, the supervised approach of the ΔDispersion+ΔMean method resulted in lower F1 scores (averaging 0.36) compared to the ΔIQR+ΔMean method, which achieved an average F1 score of 0.88 (Fig. 4(a) and (b); Supplementary Fig. 3(a) and (b)). These findings indicate that the IQR statistic is more effective in capturing DE features than dispersion under the two group comparisons. Despite the suboptimal performance of ΔDispersion, the similar performances across both model schemes between PHet and PHet (ΔDispersion), can be attributed to the Fisher’s scores and discriminative power, which was also observed in the previous Section 2.4. A majority of DE-based methods, DIDS, and DECO demonstrated comparable performance with PHet, suggesting that PHet has a sufficient discriminative performance (Supplementary Fig. 3).

**Figure 4.**
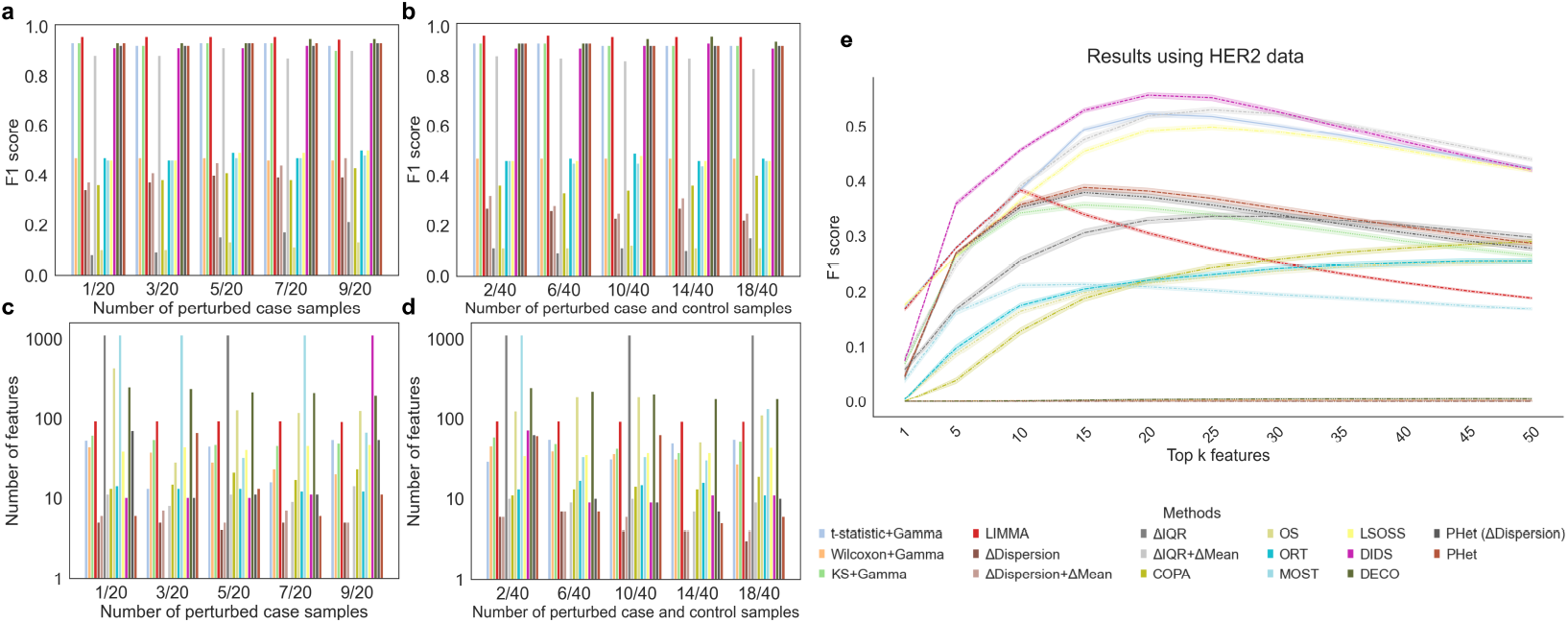
Performance comparison of PHet against 16 baseline methods using simulated and HER2 datasets. **a, b**, F1 scores of 17 different methods to detect the true DE features in perturbed samples. **c, d**, The number of selected features for each method. **a, c**, The performance of the 17 methods is compared under five scenarios, where the number of perturbed case samples varies from 1 to 9 ({1, 3, 5, 7, 9}). **b, d** The performance of the 17 methods is compared under five scenarios, which correspond to an increasing number of perturbed samples, from 2 to 18, in both case and control groups. Results (**a, b, c**, and **d**) are obtained using 10 simulated datasets (Supplementary Table 1). **e**, The performance of the 17 algorithms, using the F1 metric, in identifying the true 20 biomarkers on HER2 data (Supplementary Table 1). The algorithms were assessed based on their top k predicted features, ranging from 1 to 50.

### 2.6 Analysis of PHet’s ability to identify markers with low signals

The presence of outlier DE genes with low signals creates unique challenges in cancer studies, as they contribute to the observed heterogeneity in tumor samples. Specific algorithms such as COPA ^48^, categorized as “outlier detection”, have been proposed to address this issue. In this study, we further assessed PHet’s capability to identify DE features with low signals, referred to as outlier features. To achieve this, we evaluated the effectiveness of 25 algorithms, including seven outlier detection algorithms (Table 1), in identifying biomarkers using 1000 batches of HER2 (human epidermal growth factor receptor 2) data (Section 4.13; Supplementary Table 3). Each batch consisted of 188 samples, with 178 fixed HER2 non-amplified samples in the control group and 10 randomly drawn samples from 60 HER2-amplified samples in the case group, following the methodology described by de Ronde and colleagues ^48^. The objective is to identify the true 20 biomarkers, located on ch17q12 or ch17q21 chromosome regions, from a pool of 27,506 features in each batch. It is worth noting that these 20 specific features have limited signals. The performance of 25 algorithms was evaluated using the F1 score, which measures the algorithm’s ability to accurately identify the true 20 markers among its top predicted features. The study involved varying the number of top features from 1 to 50.

Despite not specifically designed to detect outlier features, PHet displayed a competitive performance with a mean F1 score ranging from 4% to 39% across batches (Fig. 4(e)). PHet was able to identify 10 true biomarkers within its top 20 features. This ability to detect true biomarkers sets PHet apart from dispersion based features selection and several DE and outlier detection algorithms, such as KS, Wilcoxon, OS, ORT, and COPA (Supplementary Fig. 3(e)). The performance of the DIDS method in terms of mean F1 score over batches was impressive, surpassing all other models with a score greater than 44%. However, DIDS demonstrated suboptimal performance in extracting features for subtype detection, especially when dealing with single-cell RNA-seq datasets (Section 2.3). This is not surprising considering that DIDS was primarily designed to extract features contributing to outliers in tumor samples, rather than for subtype detection. Although DECO has proven its ability to identify DE features under various complex scenarios, and outperformed DIDS in subtype detection (Section 2.3), its performance fell short and achieved a mean F1 score of less than 1% over 1000 batches. This underperformance can be attributed to expression values of 20 true markers, which impedes DECO’s ability to accurately identify these features. Consequently, DECO prioritizes other strong features, which have distinct expression profiles that differ significantly from the features of interest. Furthermore, the potential influence of batch effects may have adversely impacted the results, which represents a limitation of this experimental study.

### 2.7 PHet uncovers distinct differentiation lineages in airway epithelium

Preserving inherent complex relationships among cells is a fundamental challenge in the analysis of cell differentiation. To gain insights the mechanisms of cell differentiation, it is essential to capture the interactions and dependencies that occur among various cell types in terms of changes in gene expression. In order to assess the capability of PHet in detecting subpopulations constituting differentiation trajectories, we utilized two established scRNA-seq data of the respiratory airway epithelium ^54^: 14,163 mouse tracheal epithelial cells (MTECs) and 2,970 primary human bronchial epithelial cells (HBECs). The MTECs dataset comprises cells collected from injured and uninjured mice at different time points (1, 2, 3, and 7 days) after polidocanol-induced injury. Previous studies of lineage tracing and the regeneration process of post-injury have confirmed that basal cells differentiate into a heterogeneous population consisting of secretory, ciliated, and tuft cells, as well as other rare cell populations, such as PNECs, brush cells, and pulmonary ionocytes ^54,69,87^. For both datasets, basal cells were considered as the control group while the remaining cell types were grouped under the case category (Supplementary Table 6). Similar to the results obtained using pre-annotated markers in the previous study ^54^ (Fig. 5(a) and (h); Supplementary Figs 26(a) and 28(a)), the UMAP visualizations of PHet’s features revealed distinct cell clusters including basal and secretory cells, within both HBECs and MTECs data (Fig. 5(b) and (i); Supplementary Figs 26(b) and 27(a)). These visualizations also identified clusters corresponding to rare cell populations encompassing brush and PNECs, pulmonary ionocytes, and SLC16A7.

**Figure 5.**
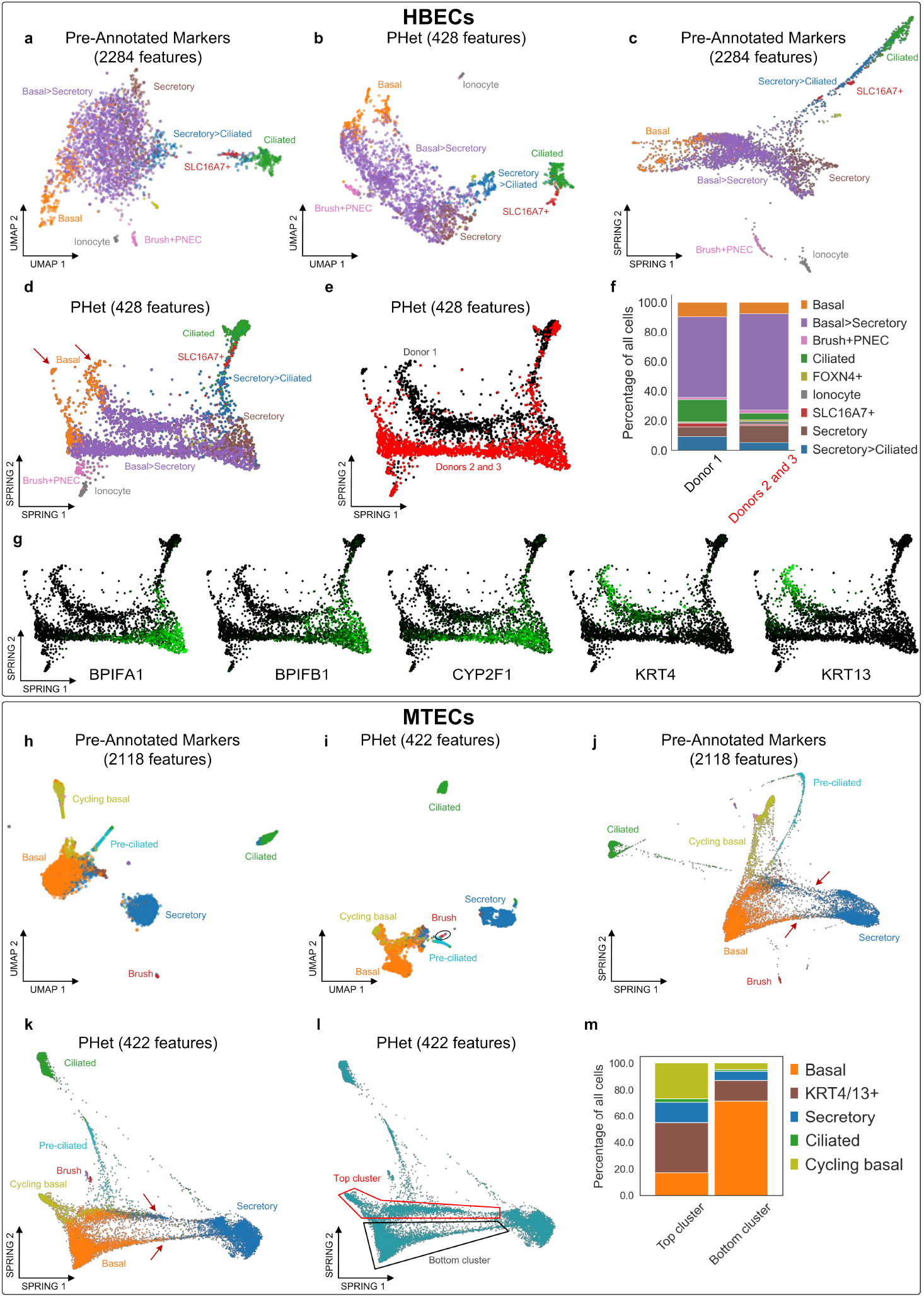
Differentiation trajectory reconstruction of HBECs and MTECs single-cell transcriptomics datasets. **a**,**b**,**c**,**d**, UMAP visualizations using pre-annotated markers and PHet’s selected features and their corresponding SPRING plots, respectively. Coloring represents the previously annotated cell types. The trajectories are visually represented by red colored arrows. **e**, A SPRING plot using PHet’s features displaying two distinct trajectories: Donor 1 (black) and Donors 2 and 3 (red). **f**, A bar plot representing the relative abundance of cell types grouped by donors. **g**, SPRING plots of the selected top features predicted by PHet. The color gradient from black to green indicated cells enriched with the corresponding feature. **h**,**i**,**j**,**k**, UMAP visualizations using pre-annotated markers and PHet’s selected features and their corresponding SPRING plots, respectively. Coloring represents the previously annotated cell types. The trajectories are visually represented by red colored arrows. **l**, Two distinct clusters are displayed in the SPRING plot, with one cluster (injured) located at the top and another (uninjured) at the bottom. **m**, The relative abundance of cells between these

To assess the role of PHet’s HD features in delineating cell differentiation trajectories, we performed SPRING visualization ^88^. SPRING is a graph-based method that constructs a cell similarity network based on their expression profiles. This aids in uncovering the structures underlying cellular differentiation trajectories. We employed the known pre-annotated markers and PHet features to generate the SPRING plots from HBECs and MTECs datasets. This comparative analysis allowed us to examine which approach more effectively captures the differentiation trajectories of cell populations. While the pre-annotated markers and PHet features displayed similar basal cell differentiation processes, only PHet revealed the presence of two distinct trajectories for the HBECs spanning basal-to-luminal differentiation, including rare cels, such as ionocytes (indicated by two red arrows in Fig. 5(c)-(d); Supplementary Fig. 26(c)-(f)). By examining the metadata of HBECs, we found that each trajectory was closely linked to a subset of donors (Fig. 5(e); Supplementary Fig. 26(f)). For instance, cells in the black cluster were specifically associated with Donor 1 that exhibited enrichment in cytokeratin genes KRT4 and KRT13^89^ (Fig. 5(g); Supplementary Fig. 26(i)). These cells could potentially serve as progenitors initiating luminal differentiation. Furthermore, an assessment of the relative abundance of cell types revealed that the secretory and basal to secretory cells were more prevalent in the red cluster (Donors 2 and 3; Fig. 5(f); Supplementary Fig. 26(g)). These cells have a significantly higher expression of the CYP2F1 gene compared to those in the black cluster (Fig. 5(g), Supplementary Fig. 26(h)-(i)). The CYP2F1 gene encodes a cytochrome P450 enzyme crucial for the metabolism of xenobiotics and endogenous compounds, aligning with the primary roles of the secretory club cells ^90^. Moreover, we also pinpointed several genes with diverse functions that were differentially expressed between the clusters (Fig. 5(g); Supplementary Fig. 26(h)-(i)). These include a member of a BPI fold protein (BPIFA1) that plays a role in the innate immune responses of the conducting airways ^91^.

UMAP visualizations of the MTECs data using both pre-annotated markers and PHet features yielded comparable observations regarding cell types (Fig. 5(h)-(i); Supplementary Figs 27(a) and 28(a)). Similar to the HBECs analysis, when SPRING was applied to both known pre-annotated markers and PHet features, two identified distinguishable cell trajectories that emerged within the MTECs dataset (Fig. 5(j)-(k); Supplementary Figs 27(b) and 28(b)). Subsequently, we characterized cell populations within these two trajectories (Fig. 5(l); Supplementary Fig. 27(c)) using cell-specific gene signatures. The lower cluster of cells was enriched with basal cells while the upper cluster contained a significant proportion of cycling basal cells (*>* 20%) (Fig. 5(m); Supplementary Fig. 27(d)). This implies a regenerative function for these cells, responding to injury by undergoing proliferation and differentiation into other cell types (Supplementary Figs 27(e)-(h) and 29). Moreover, the upper cluster contains abundant transitional cells uniquely expressing KRT4 and KRT13 genes. These cells may represent an intermediate population positioned between tracheal basal stem cells and differentiated secretory cells as suggested by previous studies ^54,69,89^. Similar cell subpopulation analysis was performed using the pre-annotated markers (Supplementary Fig. 28). The results suggest that the cycling basal cells exhibited a lower proportion in the top cluster compared to PHet’s features, as the pre-annotated markers placed cycling basal cells farther from basal cells. Therefore, this top cluster does not fully capture the information on airway regeneration following injury. In contrast, PHet’s features brought cycling basal cells closer to basal cells on both UMAP and SPRING plots (Fig. 5(i) and (k)). This is not only biologically more plausible but also makes the upper cluster more comprehensive in representing the post-injury differentiation trajectory. Taken together, these results suggest that PHet’s feature selection may provide additional insights into discovering cell subpopulations for HBECs and MTECs compared to manual marker-based analysis.

### 2.8 PHet effectively identifies subpopulations of basal cells in the MTECs dataset

Cell subtype identification is one of the most fundamental applications in single-cell data analysis. This endeavor involves assigning each cell to a specific group based on its feature expression profile, thereby shedding light on the heterogeneity and diversity inherent within cell populations in complex biological contexts, such as tissues, organs, or tumors. Moreover, the process of cell subtype identification holds great potential to uncover new biomarkers, while enhancing the understanding of cellular functions and interactions. Hence, it becomes paramount to investigate PHet’s features for the discovery of cell subtypes. In this specific case study, we reexamined the heterogeneity of basal cells in the mouse tracheal epithelial cells (MTECs) ^54^. The research findings reported by Carraro and colleagues ^92^ served as a catalyst for our exploration into the existence of potential basal cell subtypes in MTECs data. We used PHet’s features to detect basal cell clusters and compared them with dispersion based HV features and pre-annotated basal cell markers ^54^. To perform the clustering analysis, the Leiden community detection algorithm from the SCANPY package ^50^ was utilized, and the resolution hyperparameter was fixed to 0.5. The cluster quality was evaluated using the silhouette score, which measures how well each cell belongs to its assigned cluster. This metric was used due to the absence of labeled information pertaining to the basal cell subpopulations. A higher silhouette score implies a better quality of clustering.

PHet based features revealed four distinct clusters of basal cells (Fig. 6(a); Supplementary Figs 30 and 33(a)), thereby achieving the highest silhouette score (47%) (Fig. 6(h)). Each basal cluster is characterized by distinct gene expression profiles and biological functions, as observed by Carraro and colleagues ^92^. Basal-1 and Basal-3 clusters for PHet exhibited elevated expression of the canonical basal cell markers, including TPR63 (tumor protein P63) and the cytokeratin 5 (KRT5) (Fig. 6(d) and (f); Supplementary Fig. 33(d)-(f)). Conversely, cells in Basal-2 and Basal-4 clusters exhibited the reduced expression of the basal cell markers and showed enrichment for SCGB3A2/BPIFA1 for Basal-2 and MSLN/AGR2 and members of the serpin family (e.g. TSPAN1) for Basal-4, respectively (Fig. 6(d) and (f); Supplementary Fig. 33(d)-(f)). This finding suggests that these clusters represent two distinct basal cell subtypes undergoing transitions toward a luminal secretory phenotype (Supplementary Figs 30 and 33(b)-(c)). Of note, BPIFA1 was expressed in Basal-2 predominantly and was also differentially expressed between the two donor groups in the HBECs dataset (Section 2.7; Fig. 6(i)), providing further evidence that these cells are undergoing differentiation into secretory cells. A previous study has established a connection between BPIFA1 secretion and the appearance of secretory cells during mucociliary differentiation of airway epithelial cells ^93^. Furthermore, BPIFA1 is known to be upregulated in one of the secretory cell subtypes in cystic fibrosis lungs ^92^. PHet effectively identified the specific basal cell subtype linked to these phenomena. It also underscores that PHet’s features can accurately reflect the developmental trajectories of cell differentiation.

In contrast to the results obtained from PHet, the pre-annotated markers displayed a less distinct four basal subtypes, with a relatively low silhouette score of 41% (Fig. 6(b) and (h); Supplementary Fig. 34(a)). This suggests that the pre-annotated markers may not accurately capture the unique characteristics associated with basal subtypes. One particular challenge was observed in identifying the Basal-3 subpopulation, as it appeared to be situated between the Basal-1 and Basal-4 subtypes (Fig. 6(b)). This arrangement made it difficult to identify unique characteristics associated with the Basal-3 cells. Moreover, cells in the Basal-2 cluster exhibited a complex expression pattern, where both basal and secretory cell markers are being enriched in these cells, while simultaneously losing expression of BPIFA1 (Fig. 6(e), (g), and (j); Supplementary Figs 31 and 34(d)-(f)). This presents a challenge in accurately interpreting biologically meaningful signals related to cells in this cluster. These findings highlight the necessity for updating the annotated markers to characterize basal cell subtypes. The current markers may not adequately capture the diversity and distinctiveness of these subpopulations. The dispersion based HV features displayed three basal clusters with a silhouette score below 40% (Fig. 6(c) and (i); Supplementary Fig. 35(a)). A subpopulation of basal cells is observed to have a mixture of signatures from both Basal-2 and Basal-3, so it is referred to as Basal-2/3 (Supplementary Figs 32 and 35(d)-(f)). Overall, these HV features were not adequate in delineating basal subpopulations.

**Figure 6.**
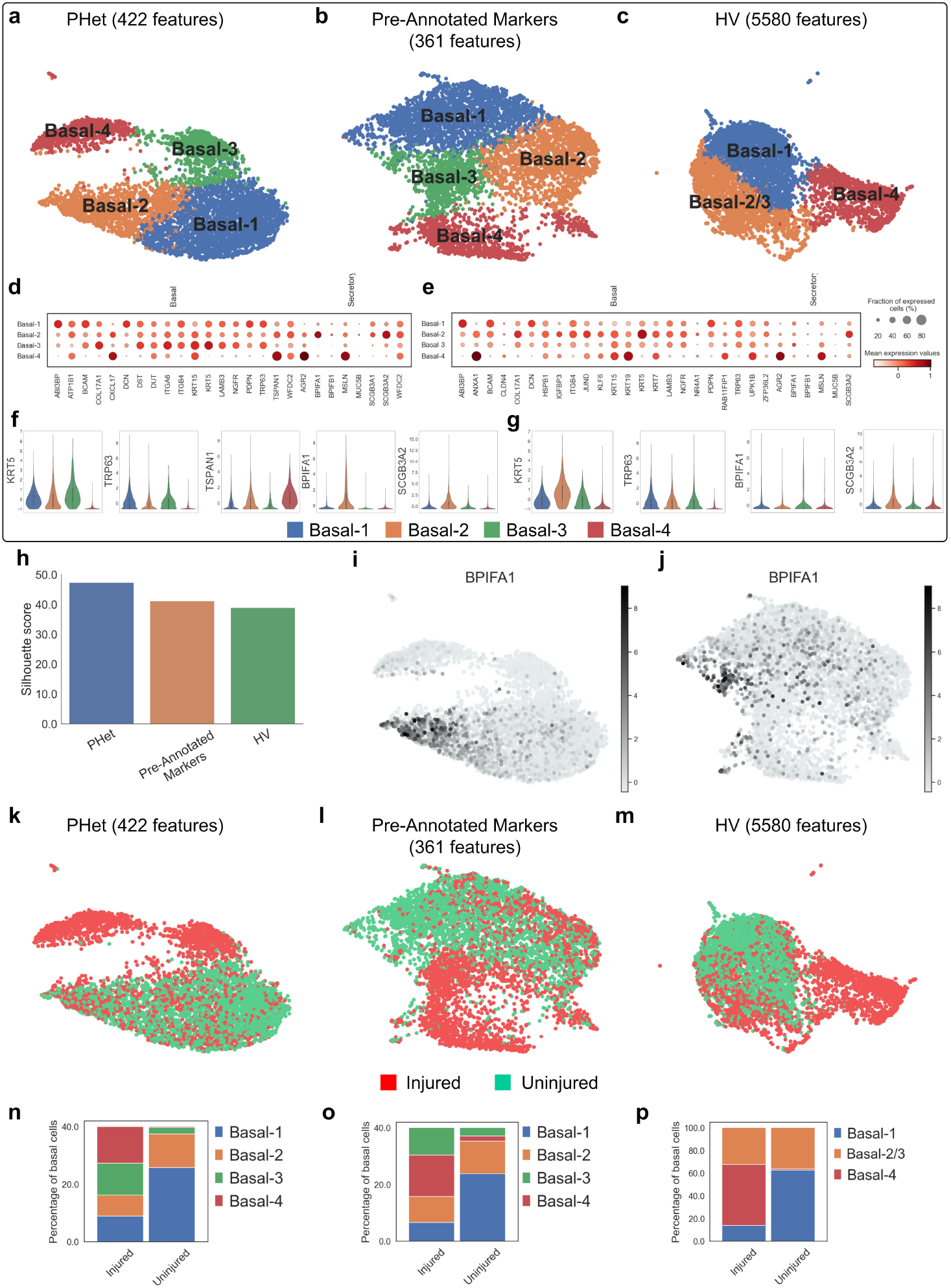
Basal cell subtype discovery in MTECs single-cell transcriptomics data. **a**,**b**,**c**, UMAP visualizations using PHet’s selected features (**a**), pre-annotated markers (**b**), and HV features (**c**). Coloring represents the detected basal cell types. **d, e**, dot plots of the expression levels of selected basal and secretory cell markers among the four basal subtypes for PHet (**d**) and pre-annotated markers (**e**). **f, g**, Violin plots of the distribution of feature expressions of selected basal and secretory cell markers among the four basal cell subpopulations for PHet (**f**) and pre-annotated markers (**g**). **h**, Silhouette scores for clustering results based on PHet’s features, pre-annotated markers, and HV features. **i**,**j**, UMAP plots of basal cells for PHet (**i**) and pre-annotated markers (**j**). These plots provide insights into the expression patterns of the secretory cell marker “BRIFA1”. **k**,**l**,**m**, UMAP visualizations of injured vs uninjured conditions using PHet’s selected features (**k**), pre-annotated markers (**l**), and HV features (**m**). **n**,**o**,**p**, The proportion of each basal subtype in the injured and uninjured cell groups given PHet (**n**), pre-annotated markers (**o**), and HV features (**p**).

In an attempt to gain insights into the occurrence of basal subtypes in both injured and uninjured conditions of MTECs data (Fig. 6(k)-(m)), we analyzed the distribution of basal cell clusters. Leveraging the PHet’s features, the Basal-4 population was observed to be almost exclusively present in the injured condition (Fig. 6(n)). In contrast, when examining the pre-annotated markers, we found that a small fraction (5%) of the Basal-4 population was present in the uninjured case (Fig. 6(o)), reassuring that these markers are insufficient for uncovering accurate subtypes at a higher resolution. The HV features constituted three clusters of basal, but their significance and biological relevance are unclear (Fig. 6(p)).

## 3 Discussion

Subtype discovery within the growing number of omics expression datasets is important for studying tissue heterogeneity, understanding cellular differentiation pathways, and identifying molecular signatures linked to different biological states or complex diseases like cancer and diabetes. Inaccurate detection of subtypes could negatively affect clinical decision-making, the development of targeted therapies, and patient treatment planning. Additionally, finding molecular signatures in high-dimensional omics data presents challenges due to factors like noise, sparseness, and heterogeneity. A crucial initial step is the selection of features associated with specific subtypes. Many current methods focus on identifying DE features, which are features that show distinct expression levels across certain biological conditions. Recent single-cell RNA-seq analyses have introduced a novel category known as HV features, representing genes with high variability regardless of conditions. In this paper, based on our deep metric learning, we introduce a new feature set called HD features, characterized by both mean expression differences and variability (IQR) discrepancies between conditions. These HD features are important for capturing the heterogeneity and diversity of subtypes while remaining discriminative between known biological conditions.

Several approaches, such as DIDS ^48^ and DECO ^49^, have been developed to extract relevant features from expression data and to identify subtypes. However, these methods are limited in capturing subtype-related features because they focus on a specific type of outlier features, which means they may miss important information that could be useful for subtyping. PHet overcomes these limitations by using IQR differences, iterative subsampling, and statistical tests. PHet assigns scores to features based on their heterogeneity and discriminability across experimental conditions and filters out irrelevant features by fitting a gamma distribution to the scores. The resulting features can then be used to cluster the data into subtypes that reflect the underlying biological heterogeneity. Based on our benchmark studies, PHet tended to outperform the existing methods by retaining a small set of features while ensuring high clustering quality for subtypes detection. Furthermore, PHet’s versatility allows for extension to multiple conditions (e.g., basal, secretory, and ciliated cells), over different omics measurements (e.g., single-cell RNA-seq, proteomics, or metabolomics), and different experimental designs (e.g., time-series or multi-factorial experiments). For optimum subtypes detection using PHet, the data should be batch-corrected and prepossessed beforehand, and the framework is not designed to account for confounding factors or artifact noise that may affect the expression measurements ^94–97^. In evaluating the performance of algorithms using DE features, the absence of verified ground truth DE features necessitates relying on the top DE features identified by LIMMA. This dependence on LIMMA-selected features introduces uncertainty, as these features may not accurately represent the true differential expression within the data.

The establishment of comprehensive atlas datasets for various organisms and tissues, exemplified by initiatives like the Human Cell Atlas ^98^ and the Mouse Cell Atlas ^99^, has paved the way for discovering and characterizing novel cell types ^100–102^. These endeavors provide invaluable insights into the diversity and functions of cellular phenotypes across diverse biological contexts and conditions, illuminating how cell populations change in disease conditions ^103,104^. PHet is well-positioned to support this initiative by enhancing the identification of unknown cell subtypes and providing deep insights into the cellular diversity of both healthy and diseased tissues. This contribution is facilitated by its detailed analysis of the essential heterogeneity in large-scale omics expression data at single-cell resolution, aligning with the goals of comprehensive atlas initiatives.

## 4 Methods

Throughout this paper, the default vector is considered to be a column vector and is represented by a boldface lowercase letter (e.g., **x**) while matrices are denoted by boldface uppercase letters (e.g., **X**). If a subscript letter *i* is attached to a matrix (e.g., **X**_*i*_), it indicates the *i*-th row of **X**, which is a row vector. A subscript character to a vector (e.g., **x**_*i*_) denotes an *i*-th cell of **x**. Occasional superscript, **x**^(*i*)^, suggests a sample or an iteration index. An uppercase calligraphy character (e.g., X) indicates a set.

### 4.1 Data preprocessing

Omics data, denoted by **M** ∈ ℝ ^*n*+*m,p*^, takes expression profile (e.g., gene) as the input and is categorized into *control* (*n* samples) and *case* (*m* samples) experimental groups, where both groups share the same set of features (*p* ∈ N). Formally, let the data matrix **X** ⊂ **M** be control samples of size *n* ∈ ℕ where each element in **X**_*i,j*_ represents the expression value for sample *i* ∈ {1, …, *n*} and feature *j* ∈ {1, …, *p*} while the data matrix **Y** ⊂ **M** be a set of case samples of size *m* ∈ ℕ where **Y**_*k,j*_ represents the expression value for sample *k* ∈ {1, …, *m*} and feature *j* ∈ {1, …, *p*}. We filtered out low-quality samples and features in the omics data. Specifically, features expressed (as non-zero) in more than 1% of samples and samples expressed as non-zero in more than 1% of features were retained. All the data were log-transformed. We did not scale datasets to unit variance and zero mean, as scaling is an intrinsic property of methods. No other additional preprocessing and normalization was performed on the data. It is important to note that throughout the manuscript, the terms “samples” and “cells” were utilized interchangeably.

### 4.2 Deep metric learning for feature embeddings

Deep metric learning (DML) aims to learn a distance metric that can measure the similarity or dissimilarity between data samples ^105^. In the context of triplet loss that considers three types of sample types: positive, negative, and an anchor, the goal of DML is to maximize the distance between the anchor-negative samples while minimizing the distance between the anchor-positive samples by a predetermined margin. This way, DML can generate low-dimensional embeddings that effectively represent the original high-dimensional features.

In order to leverage DML for obtaining feature embeddings within specific disease or cell conditions, a series of steps were followed. Initially, the preprocessed data **M** was transposed, and the values of each sample were sorted in ascending order 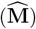. This resulted in a modified omics dataset encompassing both control and case conditions. However, due to the uneven size of control and case samples in **M**, we have utilized the “RandomUnderSampler” function from the imblearn package ^106^ to downsample from a condition that had a larger sample size. This allowed us to balance the dataset and ensure that our results were not biased towards one condition.

Prior to the DML approach, it is essential to reduce the dimensionality of the features and subsequently perform clustering. In this study, we employed UMAP ^107^ or PCA to reduce their dimensions. For UMAP, the minimum distance parameter and the number of neighbors were set to 0 and 15, respectively while for PCA the number of principal components was set to 5. After applying UMAP or PCA, we performed clustering using the k-means algorithm to partition features with the number of clusters being set to 200. This choice was informed by empirical studies considering a range of cluster numbers 100, 200, 300, 400, 500 using the Patel dataset. The analysis revealed distinct patterns when using 200 clusters from UMAP or PCA, as depicted in Supplementary Fig. 1 (i)-(j). Specifically, the HD features with elevated Euclidean distances were observed to be situated at a considerable distance from the center when employing 200 clusters. Conversely, the use of under or over 200 clusters was not able to exhibit this distinct pattern. Based on these findings, we have opted to utilize 200 clusters in our analysis for discovering feature types using the Patel data. It is important to note that the number of clusters may differ across datasets; therefore, it is essential to consider the unique characteristics of each dataset when determining the appropriate number of clusters. The resulting clustering labels were then used to construct triplets for deep metric learning. Each triplet consisted of three features: an anchor 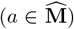, a positive 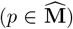, and a negative 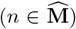. The process begins with the selection of a randomly chosen anchor feature *a*^*i*^ from a specific cluster *i*, which serves as the reference point for the triplet. Next, a positive feature *p*^*i*^ is randomly selected from the pool of features sharing the same cluster label as the anchor. Then, a negative feature *n*^*j*^ is chosen from features that have a different cluster label than the anchor, i.e., *I ≠j*. It is noteworthy to mention that the selected negative feature (*n*^*j*^) possesses the property of being situated within a specified margin (*m*) while still maintaining a substantial distance from the anchor-positive features. As a result, this negative feature is referred to as semi-hard. The process of constructing triplets is repeated for each individual feature. Finally, these triplets are fed to DML to learn feature embeddings using the triplet loss function:

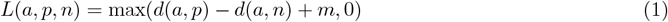

This function utilizes the Euclidean distance *d* between anchor, positive, and negative samples, along with a margin hyperparameter *m* (default is 1) to control the separation between similar and dissimilar pairs. To optimize the triplet loss function, the semi-hard triplet loss is used. This method selects triplets in which the negative feature is farther from the anchor than the positive feature, but still within a margin, i.e. *d*(*a, p*) *< d*(*a, n*) *< d*(*a, p*)+*m*. By doing so, meaningful embeddings can be generated that accurately distinguish between similar and dissimilar pairs. The semi-hard triplet loss ensures that the negative feature is neither too easy nor too hard, which further enhances the quality of the embeddings.

The architecture of our DML is a simple fully connected three-layer neural network. The input layer has a dimension of 25, followed by a hidden layer with a dimension of 10, and an output or embedding layer with a dimension of 2. To optimize the learning process for DML with triplet loss, we employed a mini-batch strategy with a batch size of 128. The Adam optimizer is utilized, and the activation function used is ReLU. We trained the network for 200 epochs to ensure convergence and optimal performance. The resulted embeddings, which have a dimension of 2, were used to calculate the Euclidean distance between any two features belonging to the same class (Fig. 1(a)). The entire process starting from sorting value was repeated 30 rounds to account for uneven sample size between case and control groups and the Euclidean distances were averaged. The implementation of our DML system is based on the TensorFlow framework ^108^.

### 4.3 PHet framework

PHet is a framework that identifies features that can separate different groups of samples in a control-case study. PHet accomplishes this goal in a pipeline procedure that is composed of six steps (see Fig. 1(o)): (1) iterative subsampling, (2) Fisher’s combined probability test, (3) measuring features discriminatory power, (4) computing feature statistics, (5) features ranking and selection, and (6) downstream analysis, such as clustering.

#### 4.3.1 Iterative subsampling

Given an annotated omics dataset (**M**), PHet computes p-values of mean differences and absolute differences in the interquartile range (ΔIQR) between control and case samples to collect feature signals. A p-value indicates the probability that, under the null hypothesis of no difference between groups, the difference calculated from the data is equal to or greater than the difference actually observed. Under the null hypothesis of no difference between groups, the p-value follows a uniform distribution between 0 and 1. Therefore, a low p-value relative to a predetermined threshold associated with statistical significance suggests that the observed difference is unlikely to be explained by chance. This, in turn, indicates that there is support for rejecting the null hypothesis in favor of an alternative hypothesis that the feature, under consideration, is indeed differentially expressed ^26^.

In order to identify DE features that contribute to subtypes determination within and across conditions, we employed the iterative subsampling with even subsampling size. The size of both subsets is determined by the closest integer to the square root of min(*n, m*). This process, called “iterative subsampling”, is repeated for a predefined number of iterations *t* ∈ ℕ to obtain a distribution of p-values for each feature. These p-values are stored in a matrix, denoted as **P**, which has dimensions *p*× *t*, where *p* represents the number of features and *t* represents the number of iterations. PHet employs the two-sample Student t-test (or Z-test if the subset size exceeds 30) as a test statistic to compute p-values.

However, p-values alone are not sufficient to extract features by their biological relevance, as they do not fully capture the heterogeneity of changes in expression for subtype detection. To address this issue, PHet utilizes ΔIQR between each condition as a measure of data variability. The IQR is defined as the difference between the first and third quartiles of the data, denoted as *q*_1_ and *q*_3_ respectively ^109^. By considering both p-values and ΔIQR, features are scored in a way that addresses complex sample distributions, which may deviate from conventional unimodal patterns, thereby improving subtype detection. The formula for ΔIQR of a subset *s* is:

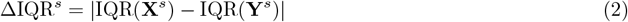

The ΔIQR values are stored in **R** ∈ ℝ ^*p×t*^, and then the mean operation for each feature is applied to obtain a vector of size *p*, i.e., **r** ∈ ℝ ^*p*^. This vector and the corresponding p-values (**P**) serve as inputs in the PHet pipeline.

#### 4.3.2 Fisher’s combined probability test

In the second step of the pipeline, PHet summarizes the obtained p-values from the previous step into test statistics using Fisher’s method for combined probability ^110^:

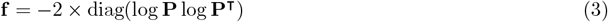

where `diag` corresponds the matrix-to-vector diagonal operation. The Fisher’s scores, represented by **f** ∈ ℝ^*p*^, provide a measure of differential consistency of probabilities across the number *t* of tests or iterations. When p-values for a feature, across *t* iterations, are small, the corresponding Fisher’s score will be large, implying that this feature is likely to be differentially expressed, and vice versa. By ranking and filtering out the features with low Fisher scores, we can reduce the dimensionality of the omics data and obtain a smaller set of DE features. The output of this step is a set of Fisher’s combined probability scores (**f**).

#### 4.3.3 Determining features discriminatory power

The obtained Fisher’s scores can effectively detect mean differences between samples in two experimental conditions; however, its discriminative ability may be constrained when confronted with complex sample distributions that deviate from the typical unimodal patterns, as alluded in Section 2.2. To address this issue, PHet uses the nonparametric two-sample Kolmogorov–Smirnov (KS) test between control and case samples to readjust the Fisher’s values. The KS test identifies a set of features by measuring the maximum difference for each feature between the two cumulative distributions without considering the type of distribution of control and case samples ^111^. To address the potential disparity in sample sizes, PHet employs a fixed sample size that is determined as the nearest integer to the square root of the smaller sample size between the control and case groups. Then, PHet compartments (binning) p-values into a predetermined number of bins *b* ∈ ℕ, each with uniform width (default is four bins with intervals defined as [0, 0.25], (0.25, 0.5], (0.5, 0.75], and (0.75, 1.0]) and assigns each bin *i* ∈ *b* a weight *w*_*i*_ ∈ [0, 1]. These weights are then associated with a specific subset of features falling within that particular interval. This weighting mechanism serves as an indicator of the discriminatory strength of each feature, enabling PHet to identify and prioritize the most relevant features for subtype detection:

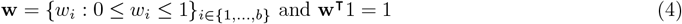

By default, the weights are set to **w** = (0.4, 0.3, 0.2, 0.1). Finally, PHet obtains the feature profile matrix as:

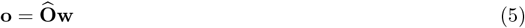

where **o ∈** ℝ ^*p*^ and 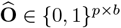. Each row in 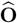 represents a one-hot encoding vector with only one element equal to 1 and the rest equal to 0. This element indicates the discriminatory strength that the corresponding feature displays. The output of this step is a vector representing the discriminatory power of each feature (**o**).

#### 4.3.4 Estimating feature statistics

In this step, PHet computes overall feature scores, referred to as the **s** statistics, using the outputs from previous steps: (1)-the average ΔIQR scores (**r**), which measures the difference in interquartile range between conditions, (2)-the Fisher’s scores (**f**), which indicates how discriminant each feature is, and (3)-the features discriminatory power (**o**), which links features to its discriminatory weight:

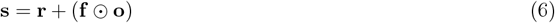

where ⊙ corresponds to the Hadamard (element-wise) product. To ensure that the values of **r** and **f** ⊙ **o** are on the same scale, we standardize both metrics. This involves dividing the values of **r** by the sum of its values and similarly, dividing **f** ⊙ **o** by the sum of its values. As can be seen in Eq. 6, **o** also adjusts Fisher’s scores to prevent the overestimation of DE features. The matrix **s** ∈ ℝ ^*p*^ holds the final statistics, where a high feature score indicates its importance for subtype detection.

#### 4.3.5 Features selection

The statistics, **s**, from the previous step are used to rank and select features. These statistics exhibit a distribution that can be effectively represented by a gamma distribution with shape *γ* and scale *β* parameters, denoted as Gamma (*γ, β*). The estimation of these parameters is facilitated using the SciPy package ^112^. Subsequently, we identify features whose scores are higher than the 1-*α* quantile of the distribution, where *α* ∈ (0, 1) is a user-defined threshold (set to 0.01 by default). The output of this step is a reduced set of features, i.e., *𝒫*^*′*^ ⊆ {1, ‥, *p*}.

#### 4.3.6 Downstream analysis

In this step, PHet leverages the identified features (*𝒫* ^*′*^) to perform a range of subsequent tasks. One of the key applications is employing clustering techniques, such as k-means ^113^ and spectral clustering ^114^, to detect subtypes within sample groups. Other applications include visualizing high-dimensional single-cell data by compressing the reduced expression matrix into a low-dimensional representation with an appropriate dimensionality reduction algorithm, enabling the identification of clusters, trajectories, or patterns that are difficult to discern in high-dimensional space. The default dimensionality reduction algorithm in PHet is UMAP (Uniform Manifold Approximation and Projection) ^58^ which is currently considered to be the superior method to collapse high-dimensional features in omics data. Other alternative methods are PCA ^115^ and t-SNE ^116^. The UMAP algorithm uses the stochastic gradient descent approach to minimize the difference between distances among samples in a higher-dimensional space and their projected lower-dimensional space. PHet also provides outputs in a tabular format that can be used in SPRING application ^88^ to explore cell differentiation trajectories.

### 4.4 Benchmark evaluation compared to existing tools

The performance of the PHet algorithm was evaluated in comparison with four DE feature analysis tools and their variants: the standard Student t-test ^75^, Wilcoxon rank-sum test ^76^, Kolmogorov–Smirnov test ^77^, and LIMMA ^78^; a dispersion-based method and its variants ^51^; an IQR-based approach and its variations ^80^; and seven outlier detection algorithms: COPA ^81^, OS ^44^, ORT^45^, MOST ^46^, LSOSS ^47^, DIDS ^48^, and DECO ^49^. In addition, we also assessed the performance of PHet using the ΔDispersion metric instead of ΔIQR. For all these methods, features were selected using either p-values or their scores with a cutoff threshold determined by the gamma distribution. Descriptions of each method are included below.

#### Student t-test (t-statistic)

The t-statistic, also known as the Student t-test, is a conventional statistical method that measures the difference in mean values between two groups of samples for each feature ^75^. This test assumes that the data follows a normal distribution and that the variance is equal across the two groups. The t-statistic can be used to find DE features between the two groups, which can help to understand the biological processes that are associated with the disease or the treatment outcome. The two-class *t* -statistic for a feature *j* is defined as:

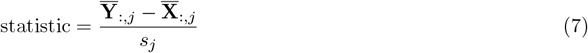

where 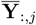 and 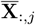 are the mean expression values of case and control samples, respectively. *s*_*j*_ is the pooled standard error estimate as:

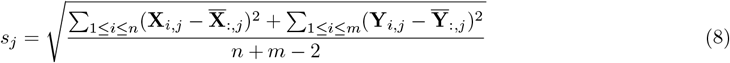

A high statistic means that the feature *j* has a large difference in mean expression between the groups, while a low statistic means that the feature *j* has a similar expression in both groups. To select important features, we followed two approaches: 1)-a p-value cutoff based on controlling false discovery rate (FDR) at *<* 0.01 significance level using the Benjamini & Hochberg method ^117^ and 2)-fit a gamma distribution to statistics and then trim features at *<* 0.01 significance level. The first approach is termed *t-statistic* and the later method is called *t-statistic+Gamma*. For both approaches, we applied a two-tailed test using the SciPy package ^112^.

#### Wilcoxon rank-sum test

This is a nonparametric statistical approach to determine whether samples from control and case groups are derived from the same distribution. The test statistic is computed as the sum of the ranks for the *n* samples from the control group. If the null hypothesis of identical population distributions holds true, these ranks represent a random sample from the *n* + *m* integers. It is important to note that under this null hypothesis, and for a larger sample size exceeding 7 for both groups ^118^, a normal approximation can be applied to report the Wilcoxon statistic:

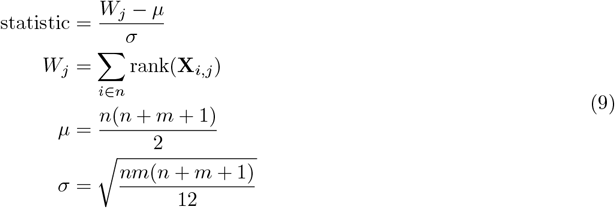

For more details, please refer to Hogg and colleagues ^118^. Having obtained p-values for all features, the important features can be determined based on controlling FDR at *<* 0.01 significance level using the Benjamini & Hochberg method ^117^. Alternatively, we can fit a gamma distribution to statistics and then select features at *<* 0.01 significance level. These two approaches are referred to as *Wilcoxon* and *Wilcoxon+Gamma*, respectively. For both approaches, we applied a two-tailed test using the SciPy package ^112^. It is worth mentioning that the Wilcoxon rank-sum test is robust to outliers and does not assume any specific distribution of the data; however, this test may exhibit reduced effectiveness when the sample size per condition is small ^119^.

#### Kolmogorov–Smirnov (KS) test

This is a nonparametric two-sample test that is used to compare the distributions of two groups of samples, such as control and case, without making any assumptions about their shapes or parameters ^77^. The test statistic for the KS test, for a given feature *j*, is the maximum absolute difference between the empirical cumulative distributions of control and case samples ^111^:

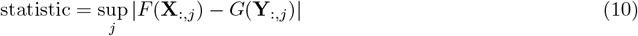

where *F* (.) and *G*(.) are the empirical distribution functions of control and case samples, respectively, and sup is the supremum function. To identify important features, we employed two approaches: *KS* and *KS+Gamma*. The KS method involves setting a p-value threshold based on false discovery rate (FDR) at a significance level of less than 0.01 using the Benjamini & Hochberg method ^117^. In contrast, the KS+Gamma method involves fitting a gamma distribution to the test statistics and trimming features at a significance level of less than 0.01. The two-tailed test was performed using the SciPy package ^112^ for these two strategies.

#### LIMMA

LIMMA (Linear Models for Microarray Data) ^78^ is a package designed to perform differential expression analysis for large-scale expression studies while remaining reasonably easy to utilize for simple experiments and small sample sizes. The package takes expression data as input, which can be log ratios or log-intensity values from microarray technologies. LIMMA then fits a linear model for each feature to the expression data and applies empirical Bayes (eBayes) and other shrinkage methods to improve the statistical inference. We applied the *limma-trend* ^79^ *in our benchmark analysis. Similar to the t-statistic, features are determined using two approaches: 1)-a p-value cutoff at <* 0.01 significance level implemented by LIMMA, termed as *LIMMA* method, and 2)-fit a gamma distribution to statistics, obtained from LIMMA, and then trim features at *<* 0.01 significance level, which is named as *LIMMA+Gamma*.

LIMMA has been a popular choice for feature detection through differential expression analyses of microarray, RNA-seq, and other types of data. However, as with the t-statistic, LIMMA assumes that the data follows a normal distribution, which may not be true for large-scale population studies. Furthermore, the increase in sample size raises the presence of potential outliers, which may violate the normality assumption and affect the p-value estimation and the false discovery rate control.

#### Dispersion based HV Features

Satija and colleagues ^51^ developed a method based on the normalized dispersion analysis to identify HV features. The method computes dispersion measures (variance/mean) for each feature from all samples and then groups features, based on their mean expression, into a fixed number of bins (20 by default). Within each bin, the dispersion measures of features are standardized to identify features that have higher variability than the great majority of the features in the bin. Typically, selecting the top HV features requires the configuration of various hyperparameters. In our methodology, we adopt an alternative approach. Rather than specifying hyperparameters, we collect the normalized dispersion values for all features and fit a gamma distribution to these values. We then select features at a significance level of less than 0.01. This approach allows us to determine HV features without the need for manual parameter tuning. We apply this method to a combination of control and case samples, which is why we refer to it as *Dispersion (composite)*. Furthermore, we extend this process by introducing three additional strategies to collect statistics or dispersion values. The first strategy, *Dispersion (per condition)*, computes dispersion values for features separately for each condition. The second strategy, Δ*Dispersion*, calculates the absolute differences of dispersion values between samples of control and case groups for all features. The third strategy, Δ*Dispersion+*Δ*Mean*, takes the feature-wise sum of the absolute differences of both dispersion and mean values between samples of the two conditions. For all these methodologies, we fit a gamma distribution to the collected values and select features at a significance level of less than 0.01.

#### Interquartile range (IQR)

This is a statistical measure that provides insight into the spread of the middle half of a feature. We propose four methods to calculate the IQR for each feature: 1)-*IQR (composite)* which involves combining the control and case samples and computing the IQR of their composite distribution. This approach takes into account the overall variability in the data; 2)-*IQR (per condition)* which calculates the IQRs of samples for each condition separately. By considering the IQRs within each condition, this method provides insights into the variability specific to each condition; 3)-Δ*IQR*, which finds the absolute differences of the IQRs between samples of the two conditions. This method highlights the features that exhibit the greatest variation between the control and case samples; and 4)-Δ*IQR+*Δ*Mean*, which combines the absolute differences in IQRs with the absolute differences in means between samples of the two conditions. By considering both IQR and mean differences, this method captures both variability and shifts in the data. The formula for ΔIQR+ΔMean for a feature *j* is defined as:

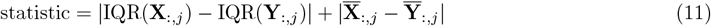

where 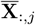 and 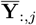 represent the mean expression values for control and case samples, respectively, for the feature *j*. To ensure that the absolute IQR difference and the absolute mean difference for each feature are on the same scale, we standardize both metrics. This process involves dividing the absolute IQR difference for each feature by the sum of the absolute IQR differences across all features. Similarly, the absolute mean difference for each feature is divided by the sum of the absolute mean differences across all features. For all IQR-based methods, a precondition to computing scores is that the expression values for each feature should be normalized to a unit variance and zero mean.

#### COPA

While the *t* -statistic is an effective way to determine the DE features, this method does not account for the heterogeneity of samples within each group. This is because the *t* -statistic computes the mean difference of features between control and case samples without providing a way to discern outliers. Typically, outlier samples have high expression values for a set of features within a group. This phenomenon is common for cancer studies where mutations can either amplify or down-regulate feature expression in only a minority of cancer (or case) samples. To address these extreme values of feature expression in the case group, Tomlins and colleagues ^43^ proposed Cancer Outlier Profile Analysis (COPA) which is defined for a feature *j* as:

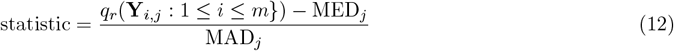

where *q*_*r*_(.) is the *rth* percentile of the expression data. MED_*j*_ is the median expression value for all samples for the feature *j* and is defined as:

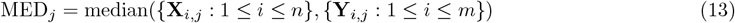

MAD_*j*_ is the median absolute deviation of expression values in all samples and is defined as:

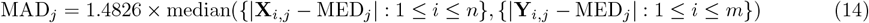

The COPA statistic utilizes the median and median absolute deviation to define outliers, where samples with high outlier scores for many features are considered to be outliers.

#### OS

A major limitation in calculating the COPA statistic is the *rth* percentile, which is a tuning parameter that depends on the data. To overcome this problem, Tibshirani and colleagues ^44^ proposed the Outlier Sum (OS) statistic. For a feature *j*, it is defined to be the sum of expressions in case samples that have values beyond *q*_75_(.) + IQR(.) for that feature as:

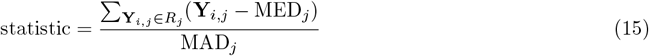

where,

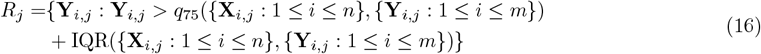

#### ORT

Wu and colleagues ^45^ proposed another variant to the OS statistic where the median of expression values for a feature *j* is computed per group based as:

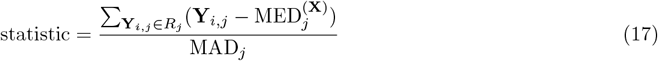

The definition of *R*_*j*_ is:

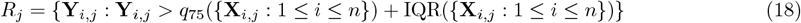

The set *R*_*j*_ is comprised of those samples in the case condition that deviate significantly from the control samples, based on the 75th percentile (*q*_75_) and the IQR values of the control samples for the feature *j*. The MAD_*j*_ is defined as:

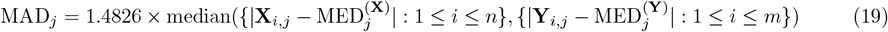

where 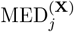 and 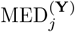 are the median expression values for control and case samples, respectively. Here, the medians are computed separately for each condition, rather than using all the samples to get a global median value. This method, called Outlier Robust T-statistic (ORT), showed good performance compared to OS and COPA on a breast cancer microarray dataset ^120^.

#### MOST

Following the ORT statistic, Lian and colleagues ^46^ proposed the Maximum Ordered Subset T-statistics (MOST) as a method to identify a subset of case samples exhibiting aberrant feature expressions. This method leverages the median expression value for control samples pertaining to a specific feature *j*, in order to locate outliers within the case condition that display abnormal signals for that particular feature. The detection of outliers is achieved by arranging the gene expression values of the case condition for a feature *j* in descending order 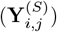 and determining an outlier threshold *k* that maximizes the statistic value (see below). Once the *k* value is determined, the top *k* sorted case samples are considered outliers. The MOST statistic is calculated as:

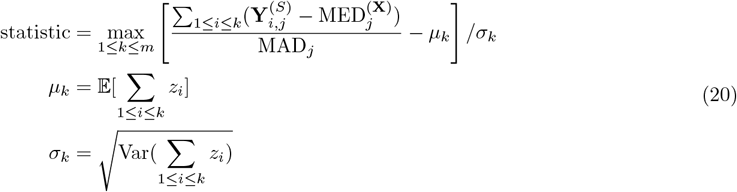

where *z*_1_ *>* … *> z*_*m*_ is the order statistics of *m* case samples generated from a standard normal distribution, as described in the original paper by Lian and colleagues ^46^. The median absolute deviation of expressions(MAD_*j*_) has the exact formula as in the ORT statistic (Eq. 19). In practical implementation, a naive iterative approach is employed to collect statistics for the *m* case samples for each feature. Subsequently, the optimal MOST statistic associated with a feature is determined by identifying the maximum statistic.

#### LSOSS

Wang and colleagues ^47^ proposed an approach for identifying outliers in case samples by detecting distinct peaks (or modes) of expression values compared to the rest of the inactivated case samples. Their approach is similar to MOST, where they found that a higher peak corresponds to activated case samples, while a lower peak indicates inactivated case samples. This outlier issue can be addressed through the concept of detecting a “change point” or “break point” in the *ordered gene expression values* of the case condition. This observation led them to introduce a novel statistical method, the Least Sum of Ordered Subset Square (LSOSS) method. The primary objective of this method is to identify the specific peak in expression values that corresponds to the outliers in case samples. By doing so, the LSOSS method endeavors to maximize the disparity between case and control samples. The LSOSS statistic for a feature *j* is defined as follows:

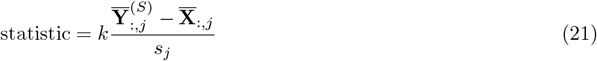

where 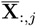 represents the mean expression value for control samples for the feature *j*. 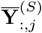 is the mean expression value of the *k* largest expression values of the case samples and is calculated based on the sorted case samples in descending order of their expression values for the feature *j*. This approach aims to facilitate the identification of distinct peaks within the case samples. To determine an optimal *k* value, which corresponds to the break point for the feature *j*, the case samples are divided into two distinct groups: *S* = {**Y**_*i,j*_ : 1 ≤ *i* ≤ *k*} and *T* = {**Y**_*i,j*_ : *k* + 1 ≤ *i* ≤ *m*}. Then, the mean and sum of squares for the feature *j* are calculated as:

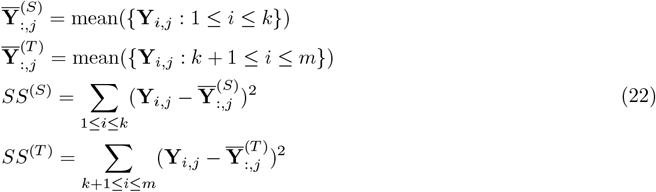

The optimal *k*(≤ *m* − 1) is determined by selecting the minimum value representing the pooled sum of squares:

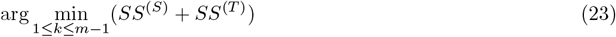

The *s*_*j*_ in the denominator of the LSOSS statistic corresponds the pooled standard error:

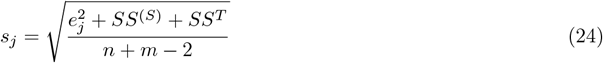

where 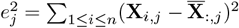 represents the sum of squares for the control samples for the feature *j*.

#### DIDS

de Ronde and colleagues ^48^ developed the Detection of Imbalanced Differential Signal (DIDS) algorithm that calculates the absolute differences in expression values by comparing the case samples with the maximum expression value observed in the control samples. These differences are then aggregated using one of three statistical functions: tanh, square, or square root to obtain the DIDS statistics:

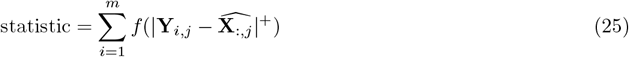

where 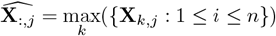 and |.| is:

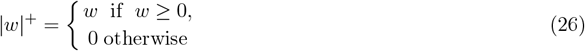

The *f* in the DIDS statistics is a strictly increasing function of either:

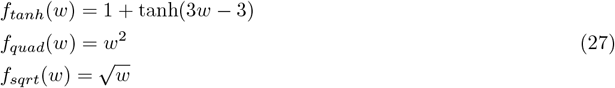

#### DECO

Campos-Laborie and colleagues ^49^ developed a method, called DEcomposing heterogeneous Cohorts using Omic data profiling (DECO), for studying the changes in feature profiles between control and case samples. DECO leverages recurrent differential analysis in combination with non-symmetrical correspondence analysis to categorize each feature into one of four model-type schemes to facilitate subtypes detection: complete, majority, minority, and mixed. The classification of a feature into those four categories is applied using the raw omics data before normalization. For a more comprehensive understanding of this method, please review the paper by Campos-Laborie and colleagues ^49^. DECO was shown to deliver promising results for the analysis of heterogeneity, biomarker identification, and subtype discovery ^49^. Furthermore, DECO has the capability to detect subpopulations without the need for labeled samples, thereby making it particularly valuable for unsupervised outlier detection.

### 4.5 Parameter settings for benchmark algorithms

We implemented the standard Student t-test, Wilcoxon rank-sum test, Kolmogorov–Smirnov test, COPA, OS, ORT, MOST, LSOSS, DIDS, ΔIQR, and PHet in Python. The same settings were utilized on all benchmark problems. For the Student t-test, Wilcoxon rank-sum test, Kolmogorov–Smirnov test, and LIMMA, we used a two-tailed test with default settings. For COPA and OS, we assigned *q*_*r*_ to 75%. We employed the ‘tanh’ scoring function for DIDS and IQR for OS, ORT, the four variants of IQR, and PHet. Additionally, we set the vector of feature weight *w* for discriminatory power to [0.4, 0.3, 0.2, 0.1] for PHet. To ensure a comprehensive analysis, we applied up and down regulations for OS, ORT, MOST, LSOSS, DIDS, all variants of IQR, and PHet. For LIMMA and Dispersion-based methods, we used the LIMMA (*limma-trend*) ^78,79^ and the SCANPY^50^ packages with default settings, respectively. To collect p-values and statistics from the Student t-test, Wilcoxon ranksum test, and Kolmogorov–Smirnov test, we utilized the SciPy package^112^. Finally, for DECO analysis, we employed the deco package written in R and applied the decoRDA module with 1000 iterations while keeping the remaining parameters at their default configurations.

### 4.6 Evaluation metrics

Performances of algorithms on different datasets were compared using seven evaluation metrics: F1, adjusted Rand index (ARI), the adjusted mutual information (AMI), silhouette, homogeneity, completeness, and Vmeasure. We applied these metrics to the original or reduced feature space before performing dimensionality reduction.

**F1**. The F1 metric is a widely used measure of the effectiveness of a classifier. It is calculated as the harmonic mean of the precision and recall, which are two metrics that evaluate how well the classifier can identify the positive and negative classes. Precision is defined as the ratio of true positive predictions to the total number of positive predictions, while recall is the ratio of true positive predictions to the total number of actual positive instances. The F1 metric is defined as:

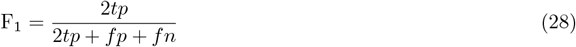

where tp, fp, and fn correspond to the true positive, false positive, and false negative features, respectively. The F1 metric ranges in [0, 1], and higher values indicate higher classification accuracy. We used the F1 score to compare the performances of algorithms on the microarray, single-cell RNA-seq, simulated, and HER2 datasets in Figs 2(a), (d), (g), and (j) and 4(a)-(b) and (e), and Supplementary Figs 2(c)-(d), 3(a)-(b) and (e), and 4-6.

#### Adjusted Rand index (ARI)

This metric is used to quantify the similarity between two different clustering results, such as comparing the current clusters with previously clustering outcomes or pre-annotated true labels. This measure is adjusted to account for random permutations, as follows:

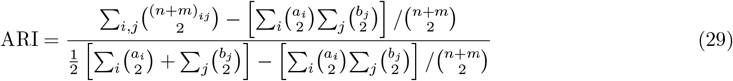

where (*n* + *m*)_*i,j*_ represents the number of samples that are concurrently classified into both clusters *i* and *j* based on the ground truth labels and the clustering results, respectively. *a*_*i*_ signifies the total number of samples belonging to cluster *i* as per the true labels, whereas *b*_*i*_ denotes the count of samples assigned to cluster *j* in accordance with the clustering labels. The ARI ranges in [0,1] with higher values indicating greater similarity between the current clustering results and either previous clustering or pre-annotated true labels. The ARI metric was used in Figs 1(f) and 2(c), (f), (i), and (l), and Supplementary Figs 2(a)-(b), 4-6 and 36-38.

#### Adjusted mutual information (AMI)

This metric is based on the mutual information (MI) between true labeled subtypes and clustering labels of a dataset, adjusted against chance. The AMI is calculated using the following formula:

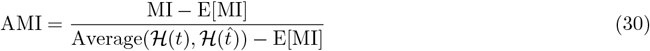

where *t* ∈ ℕ^*n*+*m*^ and 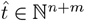 are the true subtypes and predicted clusters, respectively. E[.] is the expectation of random labeling. MI is defined as:

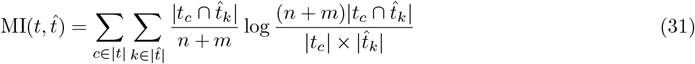

where *ℋ* (*t*) is the entropy of the truly labeled subtypes, which measures how much information is needed to identify the true labels of a sample without any clustering information while the term H(*t*^) is the entropy of the predicted clusters:

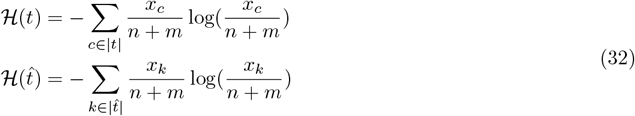

where *x*_*k*_ represents the total number of samples in the predicted cluster *k* and *x*_*c*_ is the number of samples that have the true subtype *c*. The AMI ranges in [0,1] where a higher score entails that both true labeled subtypes and clustering labels are near identical. We used the AMI metric to compare the clustering performances of algorithms for microarray and single-cell RNA-seq datasets in Supplementary Figs 4-6 and 36-38.

#### Silhouette

This metric aims to determine whether a clustering solution has successfully minimized the within-cluster dissimilarity while maximizing the inter-cluster dissimilarity. The formula for calculating the silhouette score is as follows:

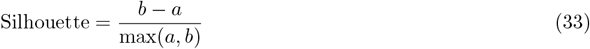

where *a* is the average intra-cluster distance, i.e., the average distance between each sample within a cluster and *b* is the average inter-cluster distance, i.e., the average distance between all clusters. The silhouette score ranges from − 1 to 1, where a high silhouette value indicates that samples are close to their own cluster and far from other clusters. We used this metric to compare the clustering performances of PHet, pre-annotated markers, and HV features for MTECs single-cell RNA-seq data in Fig. 6(h).

#### Homogeneity

This metric measures the distribution of subtypes within each predicted cluster:

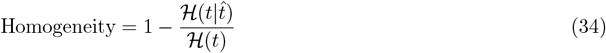

where *t* and 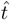 are the true subtypes and predicted clusters, respectively, and (*t*) is defined in Eq. 32. The term 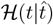 is the conditional empirical entropy, which quantifies how much information is required to determine the true labels of a sample based on its predicted cluster assignment:

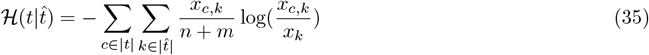

where *x*_*c,k*_ corresponds to the number of samples in cluster *k* that belongs to the true label *c* and *x*_*k*_ represents the total number of samples in the predicted cluster *k*. The homogeneity score ranges in [0, 1], where 1 denotes that each cluster contains only samples from a single true subtype (i.e., zero entropy), and 0 entails that each cluster contains samples from all subtypes. A higher homogeneity value indicates a better clustering performance. This metric was applied to compare the clustering performances of algorithms for microarray and single-cell RNA-seq datasets in Supplementary Figs 4-6 and 36-38.

#### Completeness

This metric measures the clustering distribution over each true subtype. A clustering result satisfies completeness if all samples that are members of a given subtype belong to the same cluster:

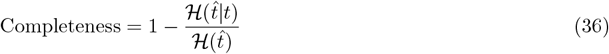

where *t* and 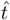 are the true subtypes and predicted clusters, respectively, and 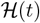 is defined in Eq. 32. 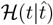 is the conditional entropy of the predicted cluster distribution given the true subtypes:

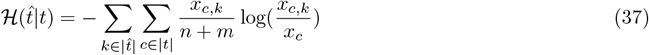

where *x*_*c,k*_ corresponds to the number of samples in cluster *k* that belongs to the true label *c* and *x*_*c*_ is the number number of samples that have the true subtype *c*. The completeness score ranges in [0, 1] where a higher value indicates a better clustering completeness. This metric was applied to compare the clustering performances of algorithms for microarray and single-cell RNA-seq datasets in Supplementary Figs 4-6 and 36-38.

#### V-measure

This metric is the harmonic mean of two scores, homogeneity and completeness:

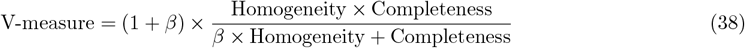

where Homogeneity and Completeness are defined in Eq. 34 and 36, respectively. *β* ∈ ℝ is a tuning parameter to favor the contributions of homogeneity or completeness. By default, *β* is set to 1. The V-measure score ranges in [0, 1] where a higher value quantifies the goodness of the clustering result. This metric was applied to compare the clustering performances of algorithms for microarray and single-cell RNA-seq datasets in Fig. 1(f) and Supplementary Figs 4-6 and 36-38.

### 4.7 Cohen’s *d* statistic

To quantify the effect size, we used results metrics, such as ARI scores, F1 scores, and predicted features from 25 methods (Section 2.3; Supplementary Fig. 2(b), (d), and (f)). These metrics allowed us to calculate the absolute values of Cohen’s *d* statistic ^121^ to assess the magnitude of differences between the compared methods, and is defined as:

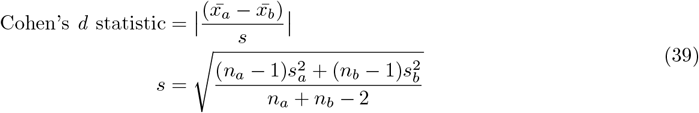

where *x*_*a*_ and *x*_*b*_ represent means of a clustering metric results for methods *a* and *b*, respectively, and *s* represents the pooled standard deviation. *n*_*a*_ and *n*_*b*_ represent the number of data points.

### 4.8 Clustering

We applied spectral ^114^ and k-means ^113^ clustering algorithms to two types of data: microarray gene expression and single-cell RNA-seq (Supplementary Table 7). The clustering process was conducted using the Scikit-learn package ^122^. To ensure robust evaluation and performance comparison against pre-annotated true labels, we set the number of clusters equal to the original patient sample types and cell types in both datasets. Other hyperparameters were maintained at their default settings. The quality of clustering results was assessed using a range of metrics including ARI, AMI, silhouette, homogeneity, completeness, and V-measure (refer to Section 4.6 for details).

### 4.9 UMAP settings

We utilized the UMAP package ^107^ in Python and fine-tuned several hyperparameters to optimize the generation of embeddings. Specifically, we set the minimum distance to 0 in order to improve the differentiation of subtypes, utilized 5 neighbors to effectively capture smaller subtypes, and executed 1000 iterations to ensure the acquisition of high-quality embeddings. The remaining UMAP hyperparameters were fixed to their default settings. We applied UMAP to generate several figures, such as Figs 1(k)-(n), 5(a)-(b) and (h)-(i), 6(a)-(c) and (k)-(l), and Supplementary Figs 9-25, 26(a)-(b), 27(a), 28(a), 30-32, 33(a)-(b), 34(a)-(b), and 35(a)-(b).

### 4.10 SCANPY settings

We used SCANPY^50^ to normalize cells, select HV features based on the minimum dispersion criterion of 0.5, perform clustering using the Leiden algorithm at resolution ranges in [0.4, 0.8], apply differential expression analysis between cell clusters using Wilcoxon rank-sum test, and visualize results using various plots, such as heatmaps. We employed SCANPY to obtain the results for Dispersion (composite), Dispersion (per condition), ΔDispersion, ΔDispersion+ΔMean, and PHet (ΔDispersion) methods in Sections 2.3, 2.6, and 2.8. Additionally, we created several figures using this library in Fig. 6(a)-(g) and (i)-(p) and Supplementary Figs 27(e)-(h), 28(e)-(h), 30-32, 33(a)-(b) and (d)-(f), 34(a)-(b) and (d)-(f), and 35(a)-(b) and (d)-(f).

### 4.11 SPRING settings

We employed SPRING ^88^ to uncover complex high-dimensional structures within single-cell gene expression data. Specifically, we configured the number of PCA dimensions to 50, the minimum number of cells for each gene to 3, the gene variability percentile to 0, the number of nearest neighbors to 5, and the minimum number of genes for each cell to 0. SPRING was applied to generate several figures, such as Fig. 5(c)-(e), (g), and (j)-(l), and Supplementary Figs 26(c)-(f) and (i), 27(b)-(c), 28(b)-(c), and 29(a)-(b) and (e).

### 4.12 Differential feature expression

In the analysis of Fig. 4(a)-(b) and Supplementary Fig. 3(a)-(b), we used LIMMA (*limma-trend*) ^78,79^ to conduct the differential expression analysis. We specifically focused on identifying the top 100 DE features with adjusted p-values below 0.01 for evaluating the performance of various algorithms.

### 4.13 Datasets

We used the following benchmark datasets: i)-two sets of simulated data (Supplementary Table 3), ii)-1000 subsets of the HER2 breast cancer gene expression data with varying sample sizes and proportions (Supplementary Table 3), iii)-17 gene expression datasets from microarray platforms with known clinical labels (Supplementary Tables 1 and 4), iv)-ten single-cell RNA-seq datasets with annotated cell identities (Supplementary Tables 2 and 5), and v)-two single-cell RNA-seq datasets from lung airway epithelial cells with mixed known and unknown cell subtypes (Supplementary Table 6).

#### Simulated datasets

A base synthetic data was generated using the madsim package ^123^ with specific configurations. The lower and upper bounds of log_2_ intensity values were set to 2 and 16, respectively. The proportion of DE features was set to 0.1. The remaining parameters in the madsim package were set to their default values. The resulting dataset consisted of 1100 features, with 93 of them being DE features. There were a total of 40 samples, evenly distributed between control and case conditions (Supplementary Table 3). The purpose of creating this base dataset was to establish a prior distribution for simulating two additional sets of data. Each of these sets consisted of five datasets drawn from the same underlying base synthetic data. The first set represented the minority model type, where random perturbations were applied to 100 features in the case condition, including some DE features. The number of perturbed case samples ranged from 5% to 45%, corresponding to 1 to 9 out of the 20 samples. The second set represented the mixed model scheme, which followed a similar procedure as the minority model. However, in this case, the perturbed samples were distributed equally among a subset of cases and controls. These two sets of simulated data were designed to represent balanced situations, where the number of case and control samples was exactly the same. We used these datasets to evaluate the performance of 25 algorithms in Table 1 as discussed in Section 2.5.

#### HER2 datasets

We followed the approach described by de Ronde and colleagues ^48^ to produce 1000 synthetic batches of HER2 (human epidermal growth factor receptor 2) data. Each batch consisted of 27,506 features and 188 samples. To create these batches, we incorporated 178 fixed control samples from a breast cancer dataset (NCBI Gene Expression Omnibus (GEO) accession ID GSE34138) representing the HER2-negative group, and 10 case samples randomly selected from 60 HER2-positive tumor samples (GEO accession ID GSE41656). It’s important to note that the case samples varied for each batch, while the control samples remained constant. The main objective behind generating this dataset was to assess the effectiveness of algorithms in predicting 20 true outlier genes. These genes are specifically located on either ch17q12 or ch17q21 (see Section 2.6). These regions were chosen due to their well-established association with HER2 amplification and breast cancer.

#### Microarray datasets

We downloaded twelve microarray datasets from the GEO database with accession numbers of GSE412; GSE3726; GSE89; GSE68956 (the Braintumor data); GSE2685; GSE83227 (the Lung data); GSE1987; GSE25055 (the BCCA data); GSE2191; GSE2535; GSE19429 (the MDS data); and GSE68907 (the Prostate data). The Glioblastoma, Leukemia, MLL, SRBCT, and DLBCL data were downloaded from Bioinformatics Laboratory at the University of Ljubljana from https://file.biolab.si/biolab/supp/bi-cancer/projections/index.html.

#### Single-cell transcriptomics datasets with known sub-types

We downloaded eight scRNA-seq datasets from the GEO repository with accession numbers of GSE67835 (the Darmanis data); GSE36552 (the Yan data); GSE81252 and GSE96981 (the Camp data); GSE84133 (the Baron data); GSE81861 (the Li data); GSE57872 (the Patel data); GSE75140 (the Knoblich data); and GSE83139 (the Wang data). The Segerstolpe data were downloaded from the EMBL’s European Bioinformatics Institute database with accessions number of E-MTAB-5060 and E-MTAB-5061. The Lake data was downloaded from the dbGaP repository with the accession number of phs000833.v7.p1

#### Single-cell transcriptomics datasets from airway epithelium

We downloaded two scRNA-seq datasets corresponding the airway epithelial cells from the GEO repository with accession numbers of GSE102580 (the HBECs and MTECs datasets).

## Data availability

All datasets analyzed in this manuscript are publicly available and were obtained from public data repositories. See Section 4.13 and Supplementary Tables 1-6 for detailed information on microarray and single-cell omics datasets used in this study. The manuscript includes the source data necessary for generating key figures.

## Code availability

The codebase for PHet is publicly available at https://github.com/kleelab-bch/phet. PHet is licensed under the MIT License. For reproducibility, all the scripts for benchmarks and case studies presented in this manuscript are available at https://zenodo.org/records/14460056.

## Acknowledgements

This work was supported by NIH, United States (Grant Numbers: R35GM133725 and R01HL163513).

## Author contributions

A.R.M.A.B., C.H., and K.L. conceived the research. A.R.M.A.B. and C.H. wrote the PHet package, performed computational analysis and experimental work, and analyzed and produced figures using the discussed data. A.R.M.A.B., C.H., and K.L. wrote the manuscript.

