## Supplementary Figure, Supplementary Table for "Heterogeneity-Preserving Discriminative Feature Selection for Disease-Specific Subtype Discovery"


| Dataset | #Samples | #Features | Subtypes | Control | Case | Description |
|---|---|---|---|---|---|---|
| BCCA [16] | 285 | 22283 | Basal, HER2+, LumA, and LumB | Basal and HER2+ | LumA and LumB | Patient samples of diagnosed tumours divided in oestrogen receptor positive and negative (ESR1+/ESR1–) |
| DLBCL [27] | 77 | 7070 | Diffuse large B-cell lymphoma (DLBCL) and Follicular lymphoma (FL) | DLBCL | FL | DLBCL and FL patient samples |
| GSE2191 [31] | 54 | 12625 | Remission and Relapse | Remission | Relapse | Patient samples with acute myeloid leukemia (AML) according to their prognosis after treatment (remission or relapse of disease) |
| GSE2535 [8] | 28 | 12625 | Responder and non-responder to imatinib treatment | Non-responder | Responder | Patients with chronic myeloid leukemia (CML) in chronic phase and their responses to the standard treatment with imatinib |
| MDS [23] | 80 | 54675 | RAEB1 and RAEB2 | RAEB1 | RAEB2 | CD34+ selected cells from bone marrow of patients suffering two closely related myelodysplastic syndrome (MDS) subtypes:RAEB1 and RAEB2 |
| Prostate [28] | 102 | 12533 | Normal tissue and Prostate tumor | Normal | Prostate | Patient samples of prostate tumors and adjacent prostate tissue not containing tumor |

**Supplementary Table 1:** Summary of six microarray datasets used for PHet's hyperparameters tuning. These datasets differ with respect to patient types, the number of samples for each group, and the feature size, offering a robust and practical framework for validating and fine-tuning PHet's hyperparameters.

| Dataset | #Cells | #Features | Subtypes | Control | Case | Description |
|---|---|---|---|---|---|---|
| Knoblich [4] | 734 | 18927 | Fetal neocortex, Dissociated whole cerebral organoid, and Microdissected cortical-like ventricle from cerebral organoid | Fetal neocortex | Dissociated and Microdissected | Cells in human cerebral organoids and fetal neocortex |
| Lake [19] | 1320 | 25051 | Excitatory and inhibitory subtypes | Excitatory subtypes: Ex1-8 | Inhibitory subtypes: In1-8 | Cells from human brain representing frontal, prefrontal, and visual cortices |
| Segerstolpe [26] | 1017 | 25525 | Alpha, Beta, Delta, Gamma, Epsilon, Ductal, Acinar, Endothelial, and Mast | Endocrine cells: Alpha, Beta, Delta, Gamma, and Epsilon | Non-Endocrine cells: Ductal, Acinar, Endothelial, and Mast | Cells from healthy human pancreatic islets |
| Wang [30] | 251 | 19950 | Alpha, Beta, Delta, Gamma, Epsilon, Ductal, Acinar, and Mesenchyme | Endocrine cells: Alpha, Beta, Delta, Gamma, and Epsilon | Non-Endocrine cells: Ductal, Acinar, and Mesenchyme | Cells from human endocrine pancreas in type 1 and type 2 diabetes |

**Supplementary Table 2:** Summary of four single-cell transcriptomics datasets used for PHet's hyperparameters tuning. These datasets cover different biological systems (e.g., human pancreatic and brain cells). In addition, the data vary in the number of cells, cell types, and feature size, offering a robust and practical framework for validating and fine-tuning PHet's hyperparameters.

| Dataset | #Samples | #Features | Subtypes | Control | Case | Description |
|---|---|---|---|---|---|---|
| Synthetic [12] | 40 | 1100 | Class 0 & 1 | Class 0 | Class 1 | Synthetic microarray dataset |
| HER2 [10] | 188 | 27,506 | HER2 negative & positive | HER2-negative | HER2-positive | Breast cancer |

**Supplementary Table 3:** Summary of simulated datasets used for benchmark evaluation.

| Dataset | #Samples | #Features | Subtypes | Control | Case | Description |
|---|---|---|---|---|---|---|
| GSE412 [6] | 110 | 8266 | Before and After therapies | Before Therapy | After Therapy | Childhood acute lymphoblastic leukemia |
| GSE3726 [7] | 52 | 22283 | Breast and Colon cancer | Breast | Colon | Breast and colon cancer patients |
| GSE89 [13] | 40 | 5724 | Ta, T1, and T2-T4 tumors | Ta & T1 tumors | T2-T4 tumors | The stage of bladder carcinoma |
| Braintumor [25] | 40 | 7129 | Normal, Medulloblastoma Glioma (Malignant Glioma), RhabdoidTu (Rhabdoid Tumor), and PNET (Primitive Neuroectodermal Tumor) | Normal (Normal Cerebella) | Glioma, Medulloblastoma RhabdoidTu, and PNET | Different embryonal tumors of the central nervous system |
| GSE2685 [17] | 30 | 4522 | Normal (Noncancerous), Diffuse Gastric, and Intestinal Gastric | Normal | Diffuse and Intestinal Gastric Tumors | Diffuse and Intestinal tumor gastric and noncancerous tissue samples |
| Glioblastoma [21] | 50 | 12625 | CG (Classic glioblastoma), CO (Classic oligodendroglioma), NG (Non-classic glioblastoma), and NO (Non-classic Oligodendroglioma) | CG and CO | NG and NO | Malignant glioma and oligodendroglioma patients |
| Leukemia [14] | 72 | 7129 | ALL-B (B Cell Acute Lymphoblastic Leukemia), ALL-T (T Cell Acute Lymphoblastic Leukemia), and AML (Acute Myeloid Leukemia) | ALL-T and ALL-B | AML | Samples from human acute myeloid and acute lymphoblastic leukemias |
| GSE1987 [11] | 34 | 10541 | Normal (Normal Lung), Squamous (Squamous Cell Lung Carcinomas), and Adenocarcinoma | Normal | Squamous and Adenocarcinoma | Adenocarcinoma, squamous cell carcinoma, and normal lung tissue samples |
| Lung [3] | 203 | 12600 | NL (Normal Lung), AD (Adenocarcinoma), SMCL (Small-cell Lung Carcinomas), SQ (Squamous Cell Lung Carcinomas), and COID (Pulmonary Carcinoids) | NL | AD, SMCL, SQ, and COID | Four different lung tumors (adenocarcinomas, small-cell lung carcinomas, squamous cell carcinomas and carcinoids) and normal lung tissue |
| MLL [1] | 72 | 12533 | ALL (Acute Lymphoblastic Leukemia), MLL (Mixed-Lineage Leukemia), and AML (Acute Myeloid Leukemia) | ALL | MLL and AML | Samples from acute lymphoblastic leukemia, mixed-lineage leukemia, and acute myeloid leukemia patients |
| SRBCT [18] | 83 | 2308 | EWS (Ewing's Sarcoma), RMS (Rhabdomyosarcoma), BL (Burkitt's Lymphoma), and NB (Neuroblastoma) | EWS and RMS | NB and BL | The small round blue cell tumors (SRBCTs) are samples from 4 different childhood tumor patients: Ewing's sarcoma, Burkitt's lymphoma, neuroblastoma, and rhabdomyosarcoma |

**Supplementary Table 4:** Summary of the eleven microarray datasets used for benchmark evaluation. These datasets differ with respect to patient types, the number of samples for each group, and the feature size, providing a comprehensive and realistic benchmark for sample subtypes detection.

| Dataset | #Cells | #Features | Subtypes | Control | Case | Description |
|---|---|---|---|---|---|---|
| Baron [2] | 7111 | 20125 | Alpha, Beta, Delta, Gamma, Ductal, Endothelial, Macrophage, Schwann, and T_cell | Endocrine cells: Alpha, Beta, Delta, and Gamma | Non-Endocrine cells: Ductal, Endothelial, Macrophage, Schwann, and T_cell | Human pancreas |
| Camp [5] | 425 | 19020 | iPSC, Definitive Endoderm, Mature Hepatocyte, Immature Hepatoblast, and Hepatic Endoderm | Hepatic Endoderm Lineage: iPSC, Definitive Endoderm, and Hepatic Endoderm | Post Hepatic Endoderm: Immature Hepatoblast and Mature Hepatocyte | Hepatocyte differentiation in fetal and adult liver |
| Darmanis [9] | 466 | 22088 | Aastrocytes, Endothelial, Microglia, Neurons, Oligodendrocytes, Hybrid, OPC (Oligodendrocyte Precursor Cells), Fetal Quiescent, and Fetal Replicating | Adult Brain Cells: Aastrocytes, Endothelial, Microglia, Neurons, Oligodendrocytes, Hybrid, and OPC | Fetal Brain Cells: Fetal Quiescent and Fetal Replicating | Adult and fetal human brain cells |
| Li [20] | 561 | 55186 | A549 (Lung Carcinoma), GM12878-B1 (Lymphoblastoid Batch 1), GM12878-B2 (Lymphoblastoid Batch 2), HCT116 (Human Colorectal Carcinoma Cell Line) H1437 (Lung Adenocarcinoma), IMR90 (Fetal Lung Fibroblast) K562 (Myelogenous Leukemia), H1-B1 (Human Embryonic Stem Cell Batch 1), and H1-B2 (Human Embryonic Stem Cell Batch 2) | Embryonic Stem Cells: H1-B1 and H1-B2 | Non-Embryonic Stem Cells: A549, GM12878_B1, H1437, HCT116, IMR90, K562, and GM12878_B2 | Two batches of human embryonic stem & lymphoblastoid cells and several tumor tissues from patients with colorectal and lung cancers |
| Patel [22] | 430 | 5948 | MGH26, MGH30, MGH28, MGH29, and MGH31 | Progenitor States: MGH26 and MGH30 | Differentiated States: MGH28, MGH29, and MGH31 | Primary human glioblastomas |
| Yan [32] | 124 | 20214 | Oocyte, Zygote, 2 Cell Stage, 4 Cell Stage, 8 Cell Stage, Morulae, Late Blastocyst, hESC Passage 0, and hESC Passage 10 | Human Preimplantation Blastomere Cells: Oocyte, Zygote, 2 Cell Stage, 4 Cell Stage, 8 Cell Stage, Morulae, Late Blastocyst | Human Embryonic Stem Cells at Passages 0 and 10: hESC Passage 0 and hESC Passage 10 | Cells from human preimplantation embryos and human embryonic stem cells (hESCs) at different passages |

**Supplementary Table 5:** Summary of six single-cell transcriptomics datasets used for benchmark evaluation. These datasets cover different biological systems (e.g., human pancreatic cells, human brain cells, and liver bud cells). In addition, the data vary in the number of cells, cell types, and feature size, providing a comprehensive and realistic benchmark for cell subpopulations recovery.

| Dataset | #Cells | #Features | Subtypes | Control | Case | Description |
|---|---|---|---|---|---|---|
| HBECs [24] | 2970 | 25475 | Basal, Basal>Secretory, Secretory, Secretory>Ciliated, Ciliated, SLC16A7+, Brush+PNEC, Ionocytes, and FOXN4+ | Basal | Basal>Secretory, Secretory, Secretory>Ciliated, Ciliated, SLC16A7+, Brush+PNEC, Ionocytes, and FOXN4+ | Human bronchial epithelial cells |
| MTECs [24] | 14163 | 28205 | Basal, Cycling Basal (homeostasis), Cycling Basal (regeneration), Krt4/13+, Secretory, Pre-ciliated, Ciliated, Brush, PNEC, and Ionocytes | Basal, Cycling Basal (homeostasis), and Cycling Basal (regeneration) | Krt4/13+, Secretory, Pre-ciliated, Ciliated, Brush, PNEC, and Ionocytes | Mouse tracheal epithelial cells |

**Supplementary Table 6:** Summary of the lung airway epithelium transcriptomics datasets.

| Dataset | Clustering method |
|---|---|
| GSE412 [6], GSE2685 [17], GSE1987 [11], MLL [1], SRBCT [18], Camp [5], Darmanis [9], Li [20], Patel [22], and Yan [32] | K-means [15] |
| GSE3726 [7], GSE89 [13], Braintumor [25], Glioblastoma [21], Leukemia [14], Lung [3], and Baron [2] | Spectral [29] |

**Supplementary Table 7:** Types of clustering applied to each dataset for benchmark evaluation.

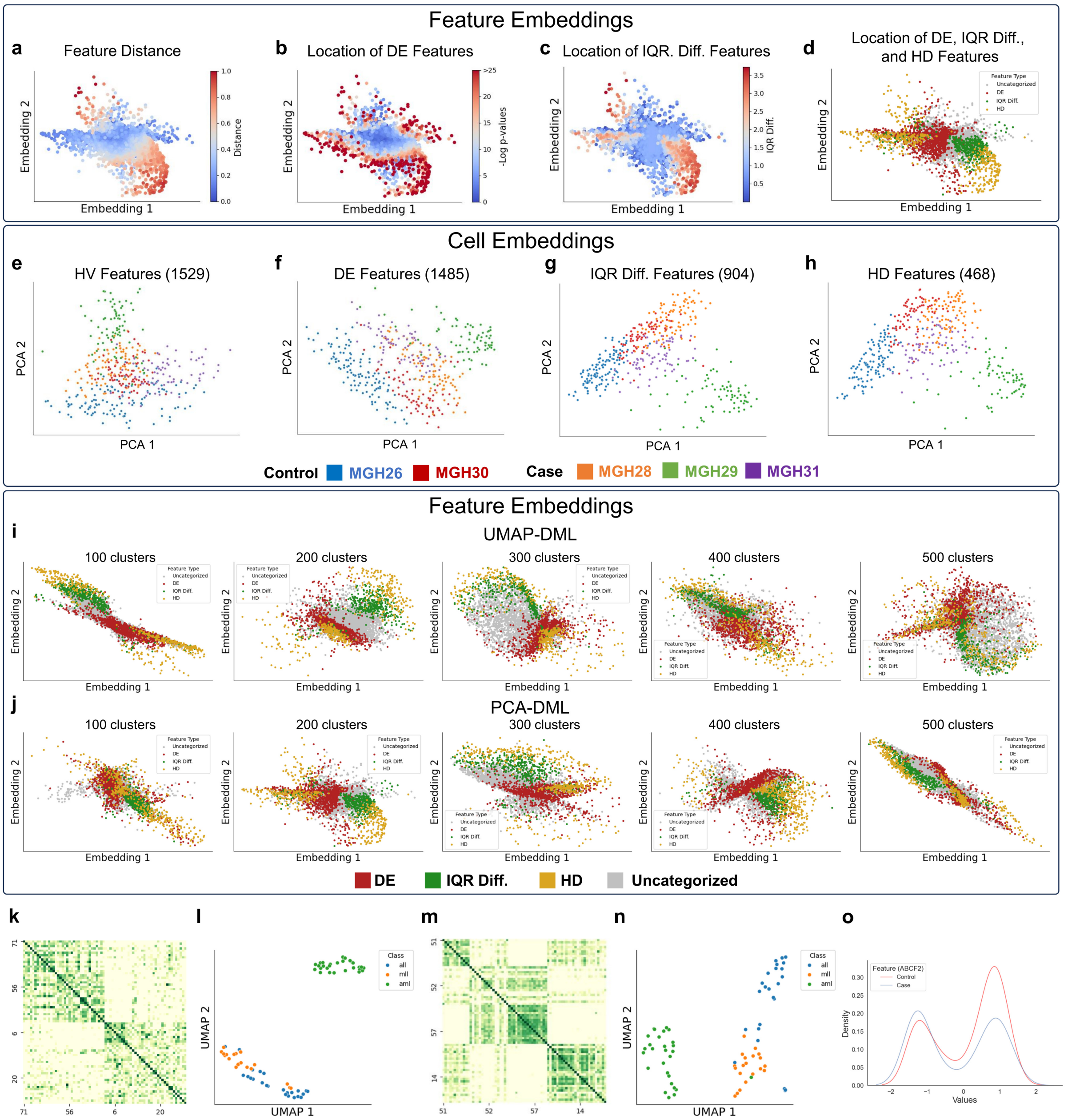


**Supplementary Figure 1:** Characterization of features obtained from deep-metric learning (DML) and limitation of discriminative DE features. **a**,**b**,**c**,**d**, Scatter plots of feature embeddings with color representing the distance between the same feature of different conditions (**a**), logged p-values from mean differences (using z-test) between conditions of features (**b**), IQR difference between conditions (**c**), and location of respective feature types on the plot (**d**). The original feature space of the Patel data underwent dimensionality reduction through principal component analysis (PCA), resulting in the extraction of 5 principal components with the highest explained variance ratio. These reduced features serve as the input for the DML process to generate feature embeddings. **e**,**f**,**g**,**h**, PCA visualizations of 1529 HV features (**e**) (based on a dispersion threshold of $> 0.5$), 1485 DE features (**f**) (based on the z-test at significance level of 0.01), 904 $\Delta$IQR features (**g**) (based on a threshold of $> 0.4$), and 468 HD features (**h**) that are intersection of both DE and $\Delta$IQR features. **i**,**j**, Scatter plots depicting the UMAP-DML and PCA-DML feature embeddings, colored according to the location of the respective feature types across a range of clusters 100, 200, 300, 400, 500. **k**,**l**,**m**,**n**, Limitation of discriminative DE features in detecting subtypes from the MLL microarray dataset. Similarity heatmap (**k**) and UMAP (**l**) of discriminative DE features. Similarity heatmap (**m**) and UMAP (**n**) of discriminative DE and HV features. **o**, the distribution of "ABCF2" feature expression values in the Patel data.

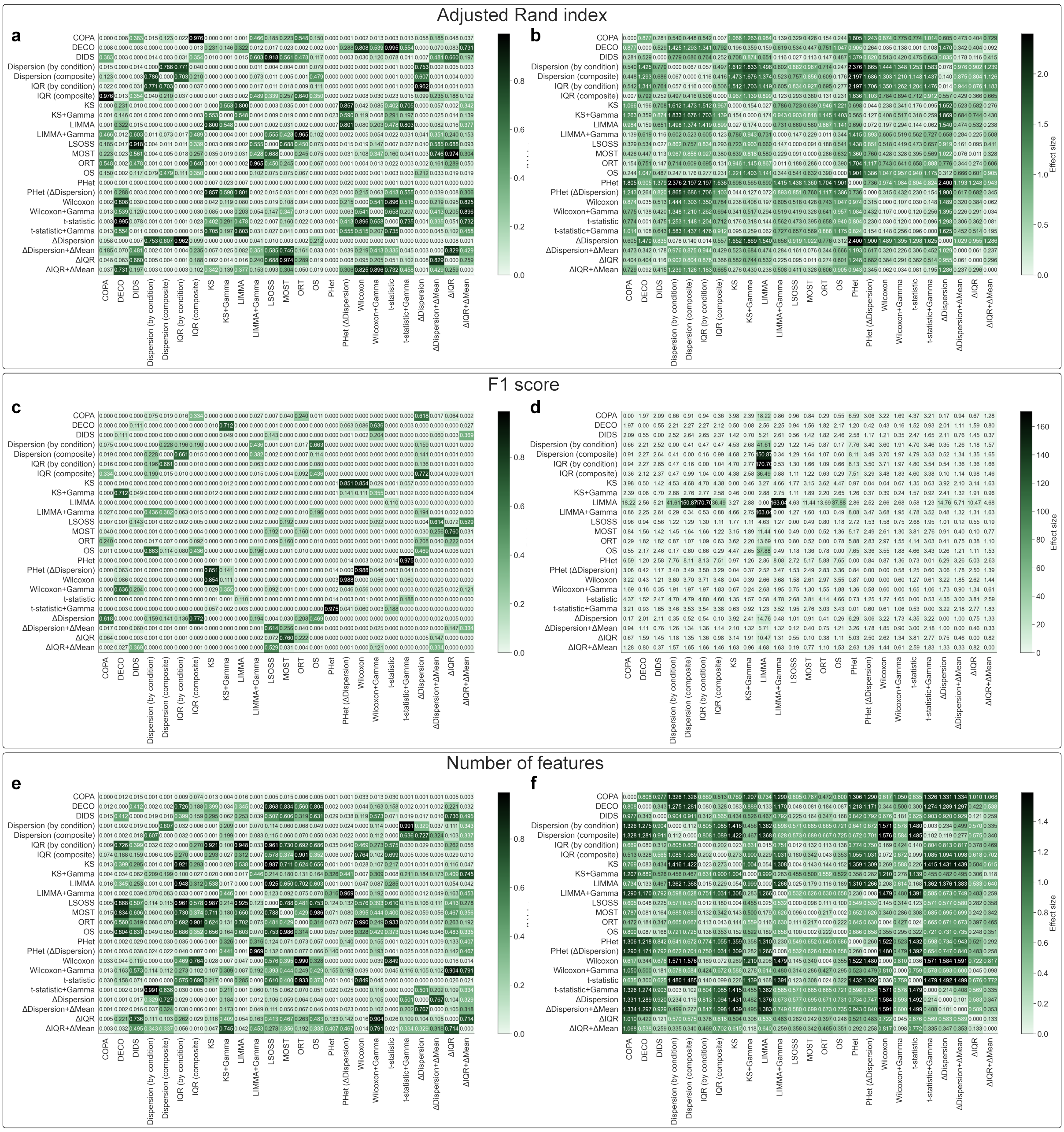


**Supplementary Figure 2:** The results of the paired two-sided t-test and Cohen's $d$ effect size of 25 methods using eleven microarray and six single-cell transcriptomics datasets (Supplementary Tables 4 and 5). Paired two-sided t-test and Cohen's $d$ effect size based on ARI scores ((**a**) and (**b**)), F1 scores ((**c**) and (**d**)), and the number of selected features ((**e**) and (**f**)) for each method.

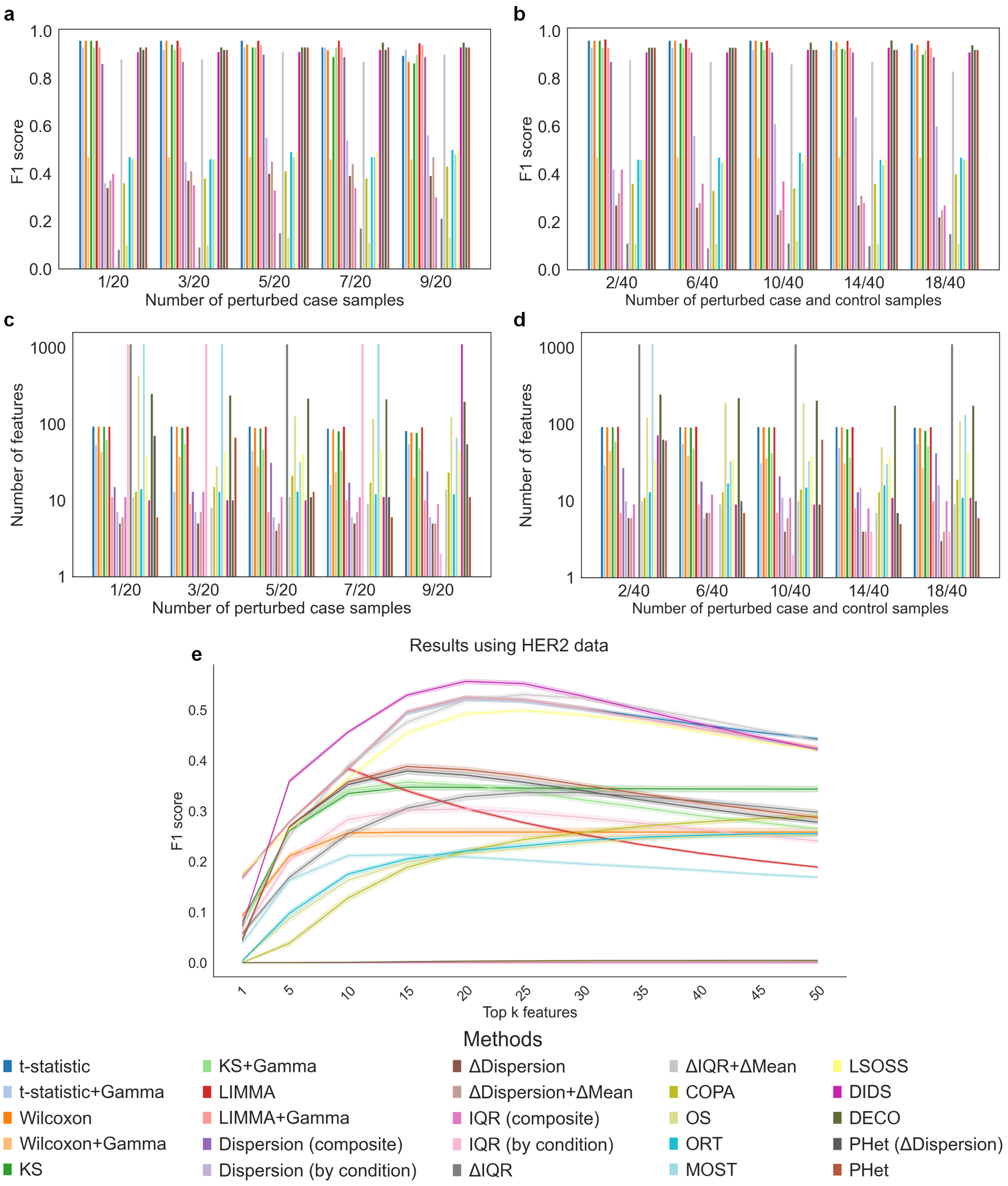


**Supplementary Figure 3:** Performance comparison of PHet against 24 baseline methods using simulated and HER2 datasets. **a**,**b**, F1 scores of 25 different methods to detect the true DE features in perturbed samples. **c**,**d**, The number of selected features for each method. **a**,**c**, The performance of the 25 methods is compared under five scenarios, where the number of perturbed case samples varies from 1 to 9 ($\{1, 3, 5, 7, 9\}$). **b**,**d** The performance of the 25 methods is compared under five scenarios, which correspond to an increasing number of perturbed samples, from 2 to 18, in both case and control groups. Results (**a**,**b**,**c**, and **d**) are obtained using 10 simulated datasets (Supplementary Table 3). **e**, The performance of the 25 algorithms, using the F1 metric, in identifying the true 20 biomarkers on HER2 data. The algorithms were assessed based on their top k selected features, ranging from 1 to 50.

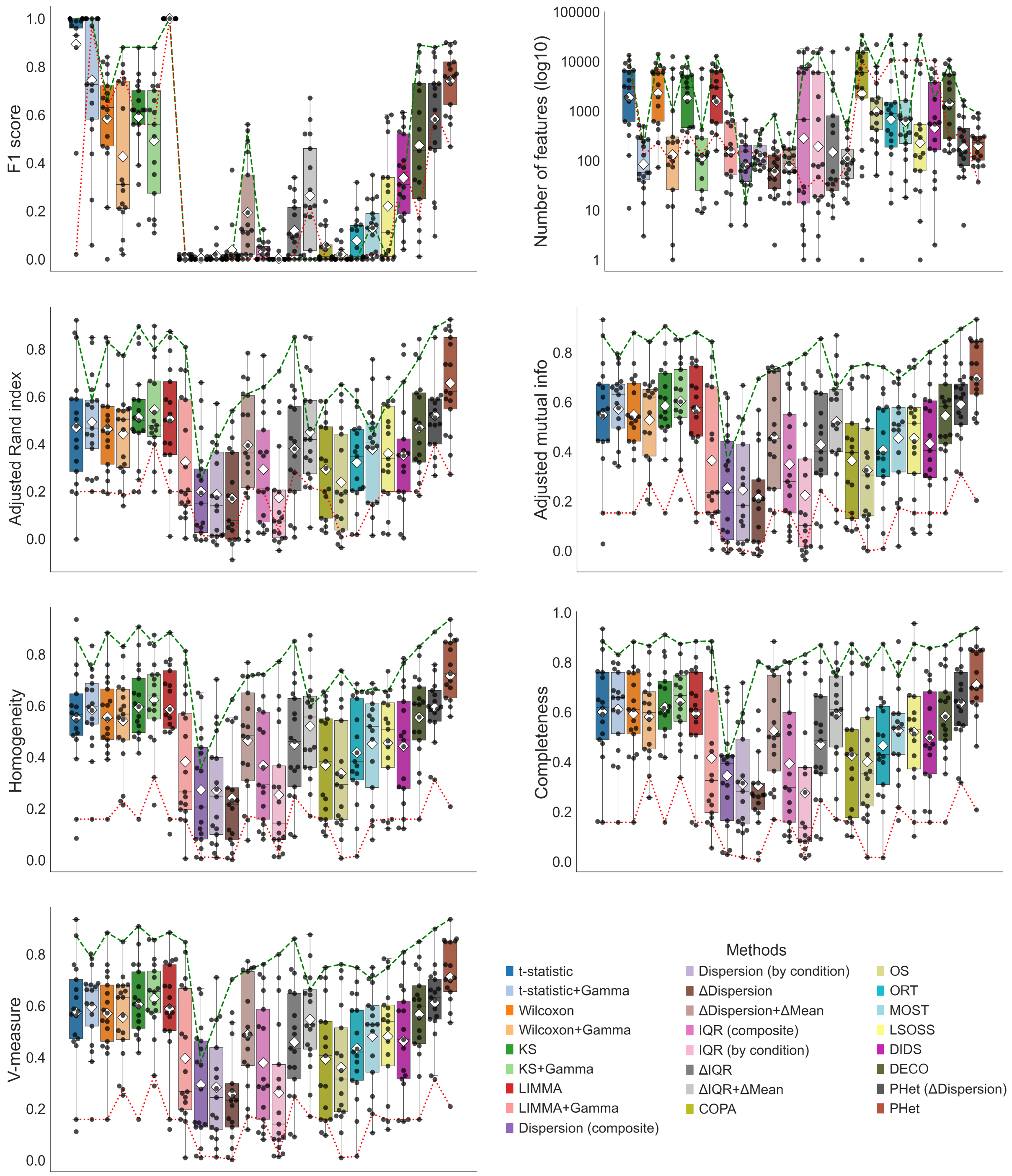


**Supplementary Figure 4:** PHet demonstrates competitive performance when compared to other baseline methods across eleven microarray and six single-cell transcriptomics datasets. Boxplots showing the performance of PHet and baseline methods evaluated using the number of selected features, F1 score, adjusted Rand index, adjusted mutual information, homogeneity, completeness, and V-measure. The box plots show the medians (centerlines), first and third quartiles (bounds of boxes), and 1.5× interquartile range (whiskers). A ◊ symbol represents a mean value. A green dashed line highlights the best-performing result achieved by PHet on each dataset for each metric, while a red dashed line signifies the worst-performing result.

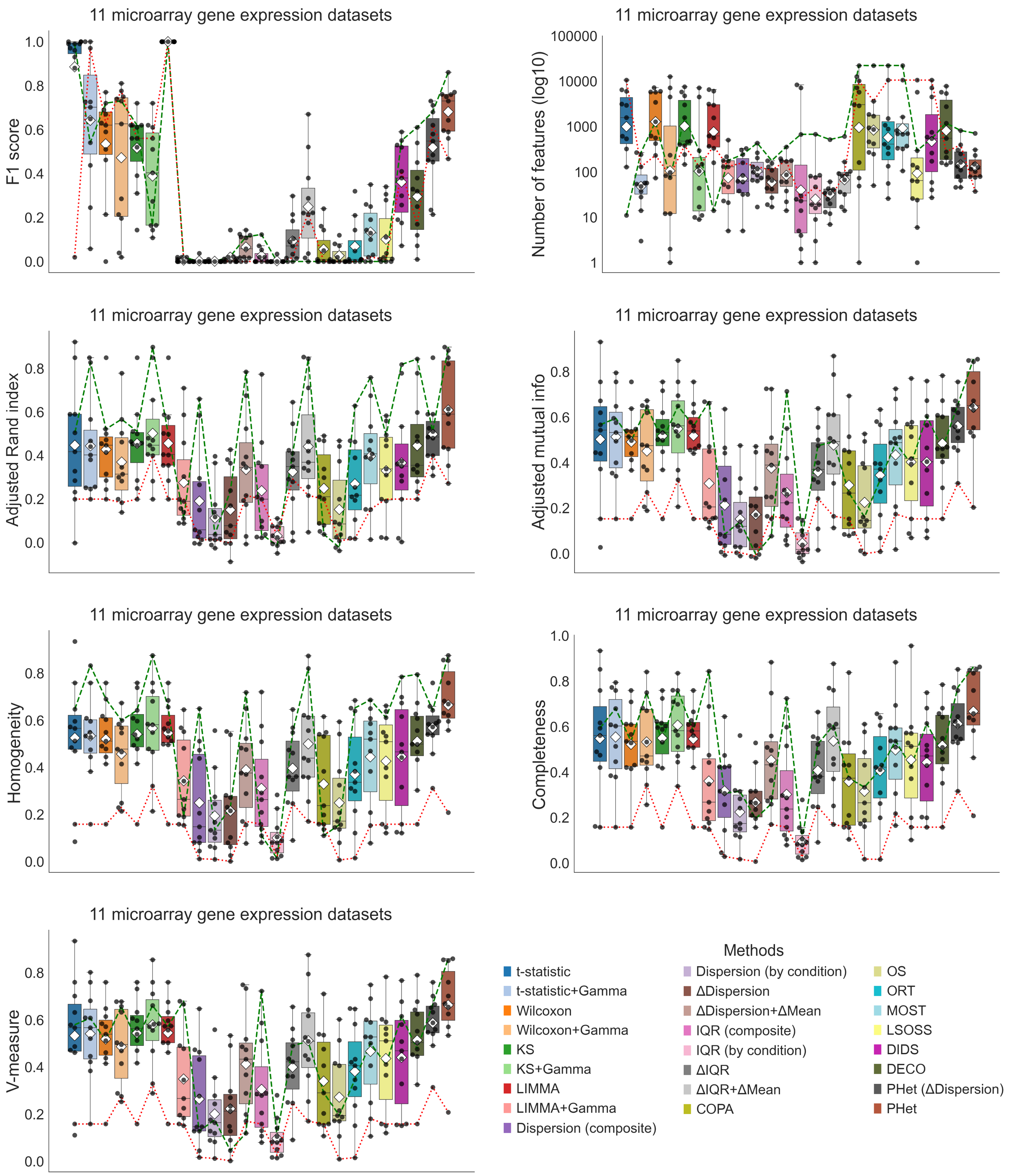


**Supplementary Figure 5:** PHet demonstrates competitive performance when compared to other baseline methods across eleven microarray datasets. Boxplots showing the performance of PHet and baseline methods evaluated using the number of selected features, F1 score, adjusted Rand index, adjusted mutual information, homogeneity, completeness, and V-measure. The box plots show the medians (centerlines), first and third quartiles (bounds of boxes), and 1.5× interquartile range (whiskers). A ◊ symbol represents a mean value. A green dashed line highlights the best-performing result achieved by PHet on each dataset for each metric, while a red dashed line signifies the worst-performing result.

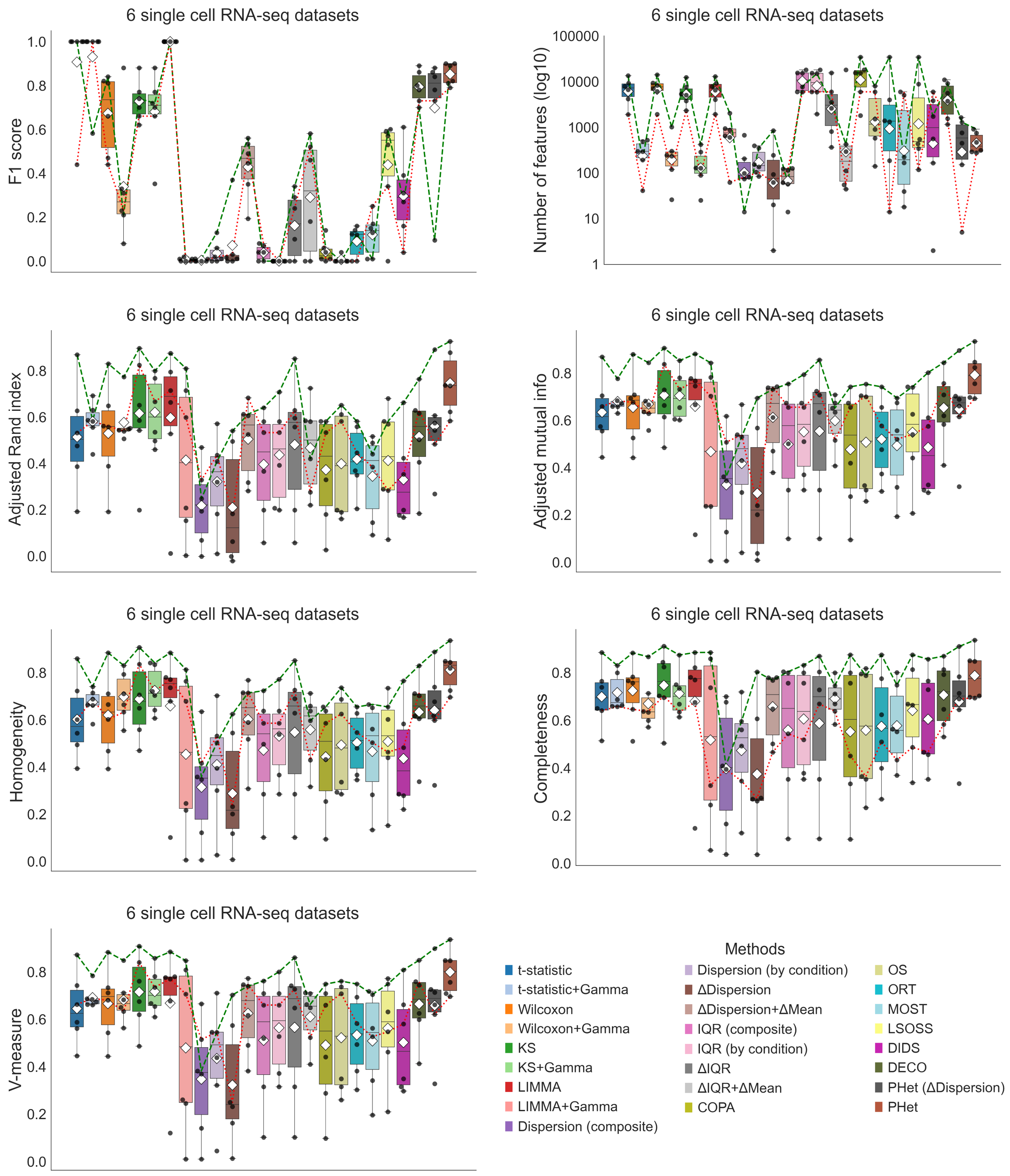


**Supplementary Figure 6:** PHet demonstrates competitive performance when compared to other baseline methods across six single-cell transcriptomic datasets. Boxplots showing the performance of PHet and baseline methods evaluated using the number of selected features, F1 score, adjusted Rand index, adjusted mutual information, homogeneity, completeness, and V-measure. The box plots show the medians (centerlines), first and third quartiles (bounds of boxes), and 1.5× interquartile range (whiskers). A ♢ symbol represents a mean value. A green dashed line highlights the best-performing result achieved by PHet on each dataset for each metric, while a red dashed line signifies the worst-performing result.

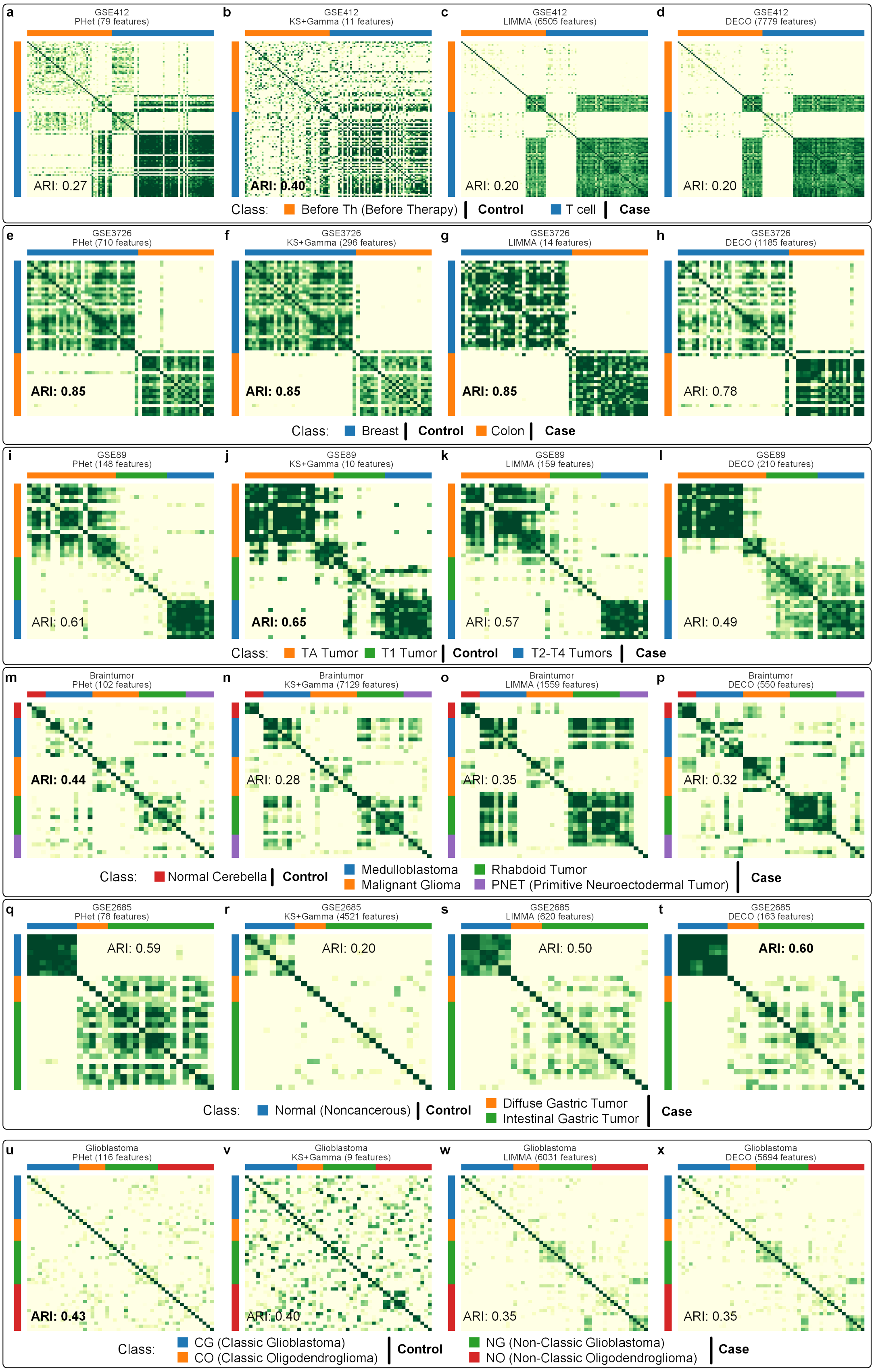


**Supplementary Figure 7:** Heatmaps displaying the clustering results of PHet and the top three methods are presented for six microarray datasets. The datasets are: GSE412 (**a-d**), GSE3726 (**e-h**), GSE89 (**i-l**), Braintumor (**m-p**), GSE2685 (**q-t**), and Glioblastoma (**u-x**). For each method, the selected features are used. The bold font ARI score indicates the best performing method for the corresponding data in the comparison.

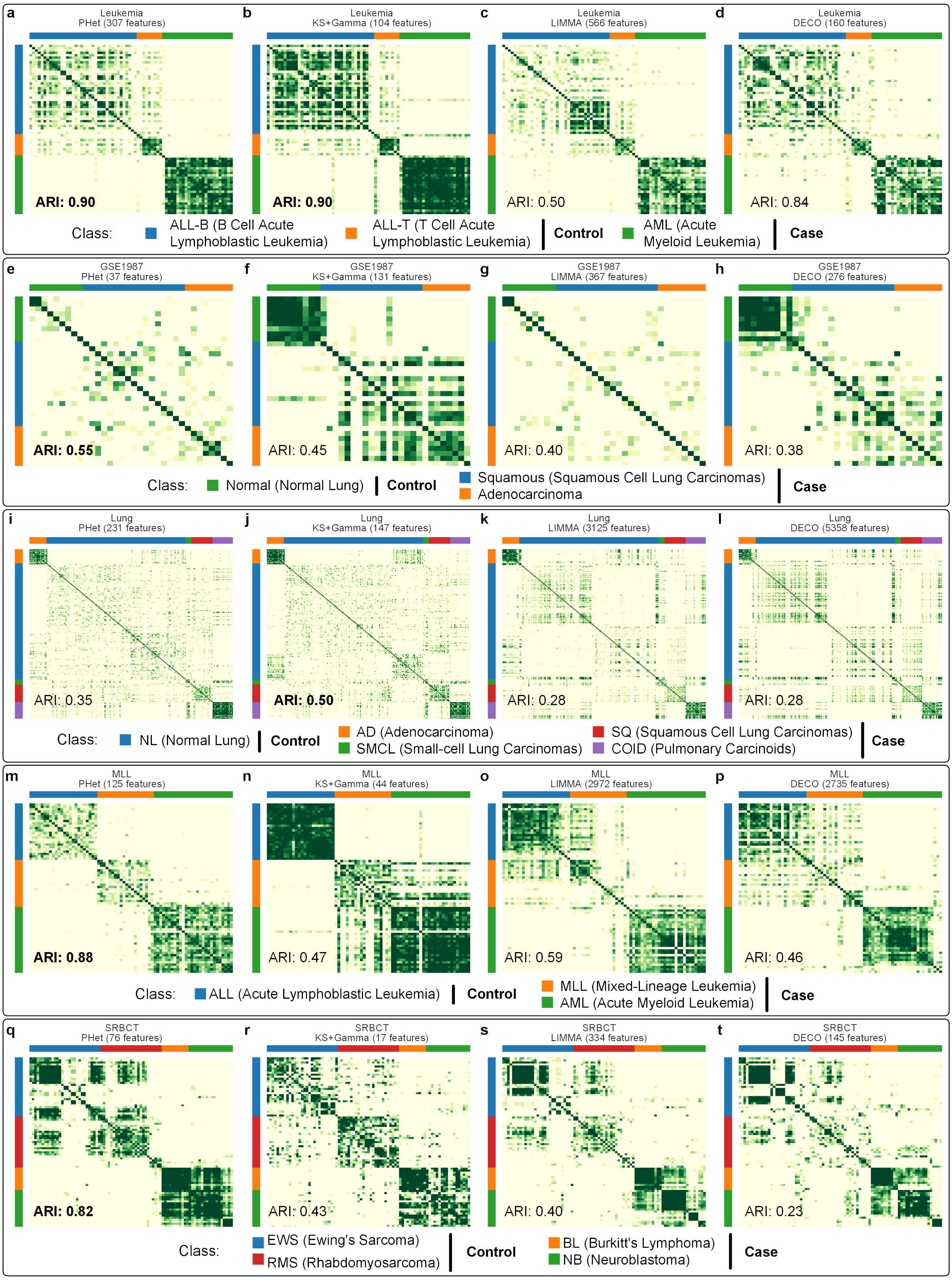


**Supplementary Figure 8:** Heatmaps displaying the clustering results of PHet and the top three methods are presented for five microarray datasets. The datasets are: Leukemia (**a-d**), GSE1987 (**e-h**), Lung (**i-l**), MLL (**m-p**), and SRBCT (**q-t**). For each method, the selected features are used. The bold font ARI score indicates the best performing method for the corresponding data in the comparison.

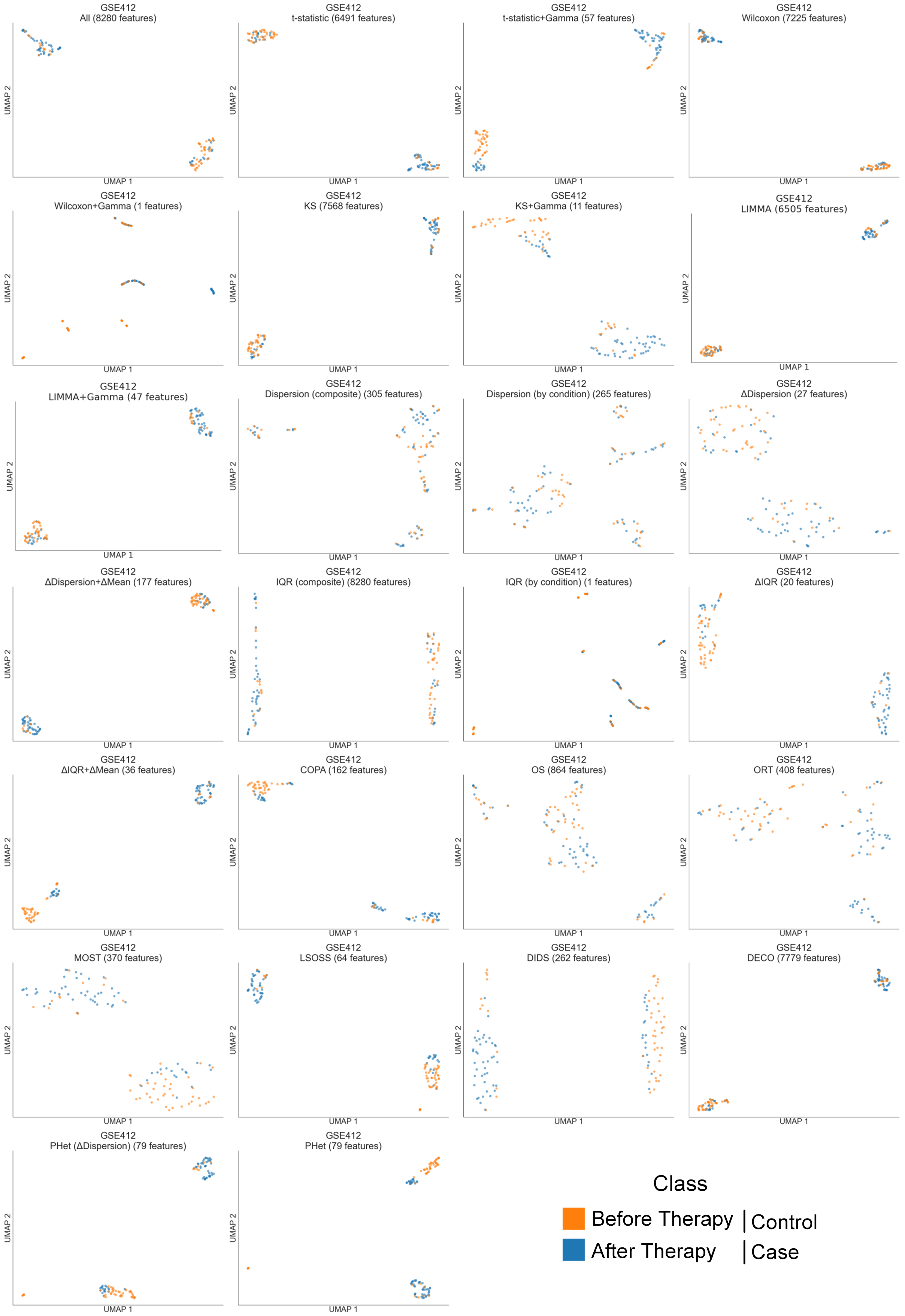


**Supplementary Figure 9:** UMAP visualizations of the childhood acute lymphoblastic leukemia microarray gene expression dataset (GSE412) [6] using all and selected features by each method, colored by original patient types (Supplementary Table 4).

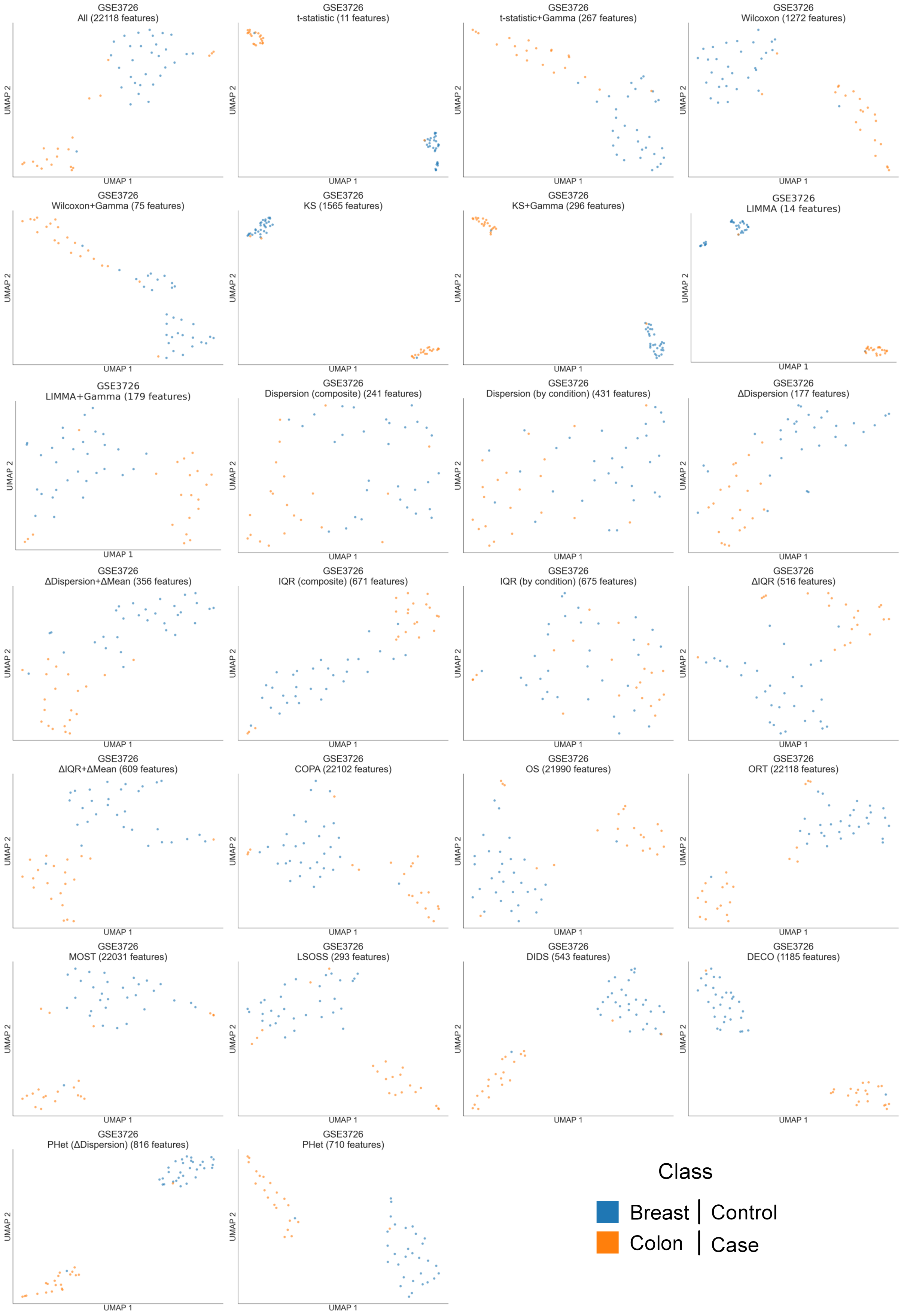


**Supplementary Figure 10:** UMAP visualizations of the breast and colon microarray gene expression dataset (GSE3726) [7] using all and selected features by each method, colored by original patient types (Supplementary Table 4).

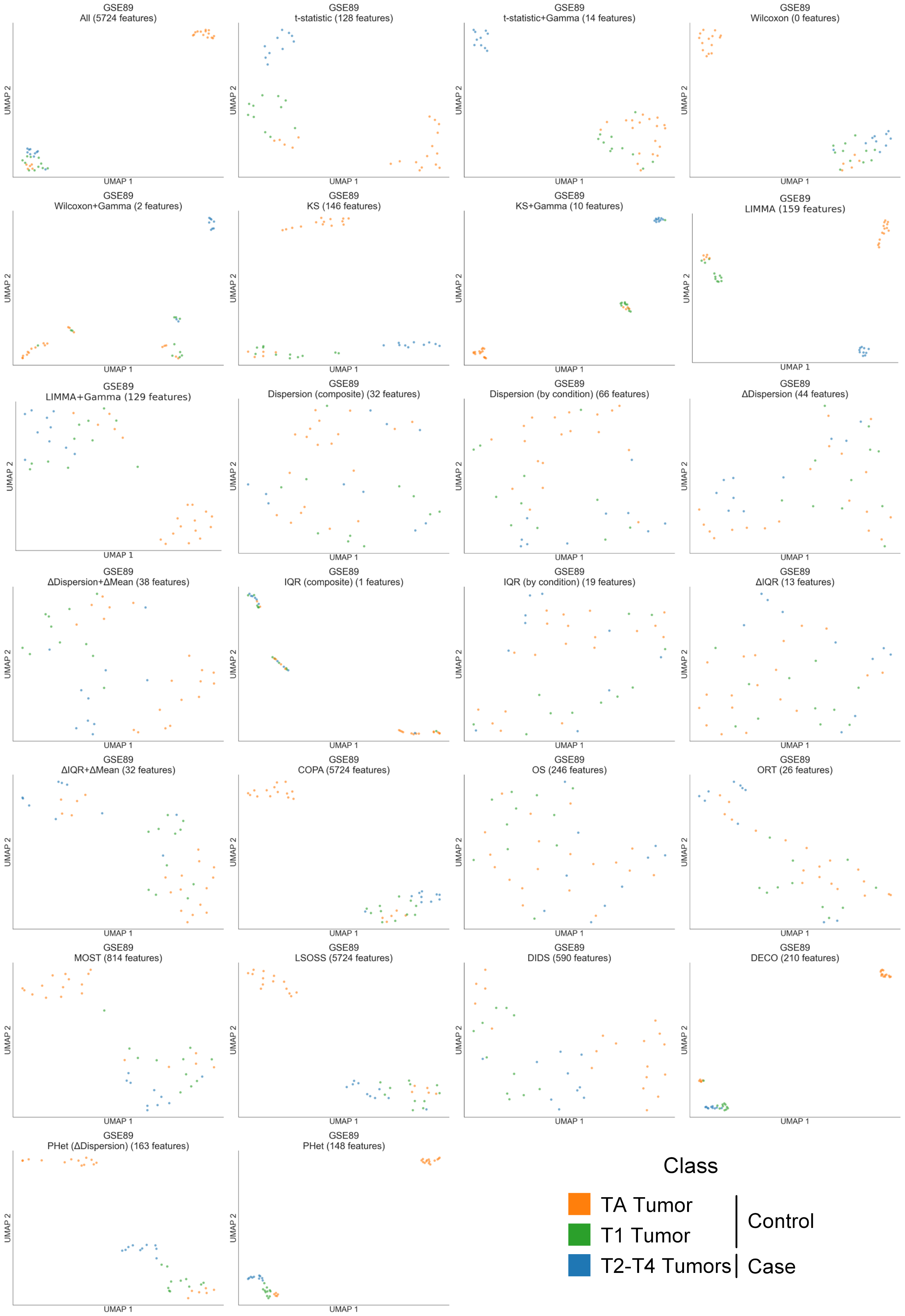


**Supplementary Figure 11:** UMAP visualizations of different stages of bladder carcinoma (Ta, T1, and T2-T4) from the bladder cancer microarray gene expression dataset (GSE89) [13] using all and selected features by each method, colored by original patient types (Supplementary Table 4).

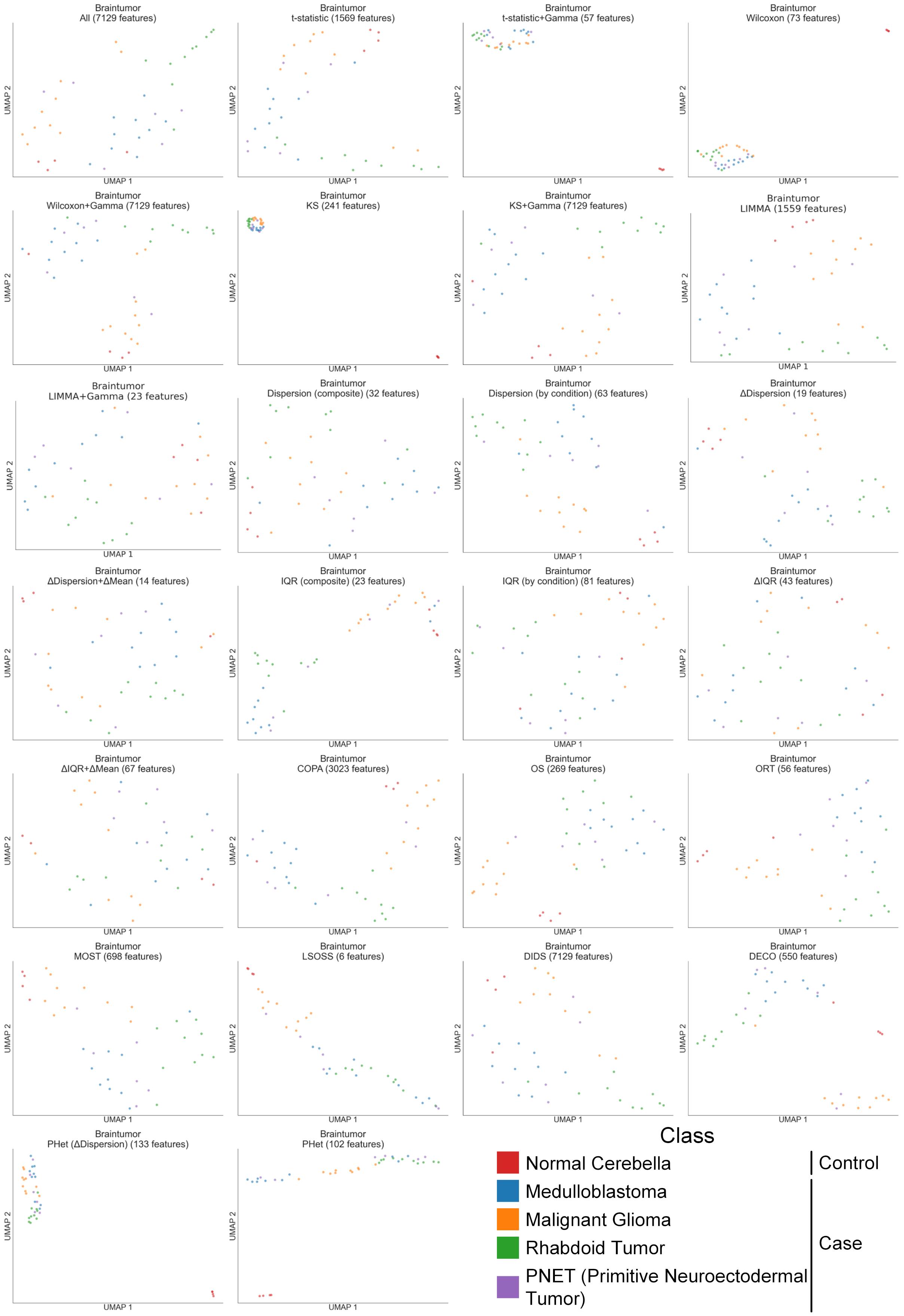


**Supplementary Figure 12:** UMAP visualizations of the braintumor microarray gene expression dataset (Braintumor) [25] using all and selected features by each method, colored by original patient types (Supplementary Table 4).

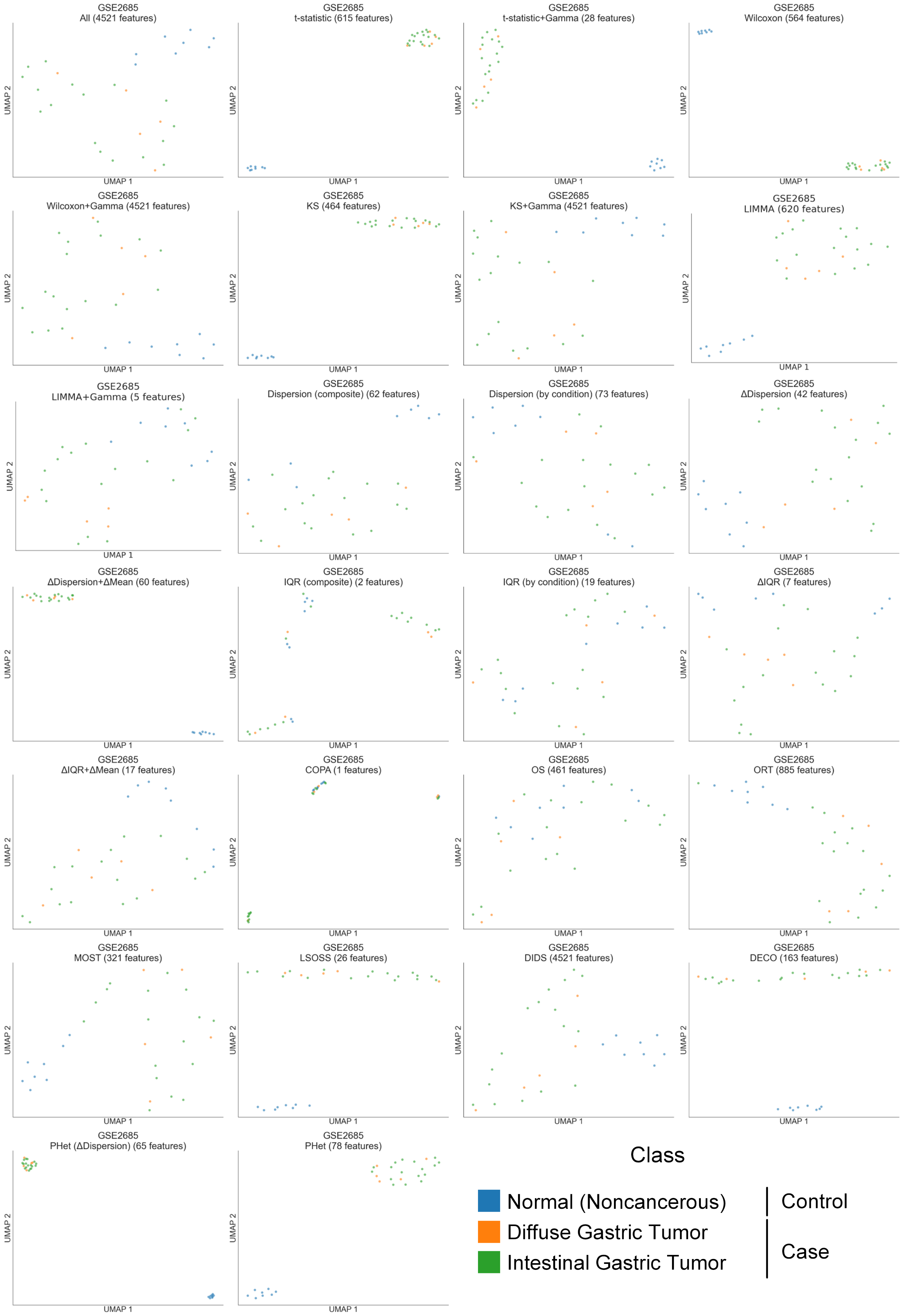


**Supplementary Figure 13:** UMAP visualizations of the gastric cancer microarray gene expression dataset (GSE2685) [17] using all and selected features by each method, colored by original patient types (Supplementary Table 4).

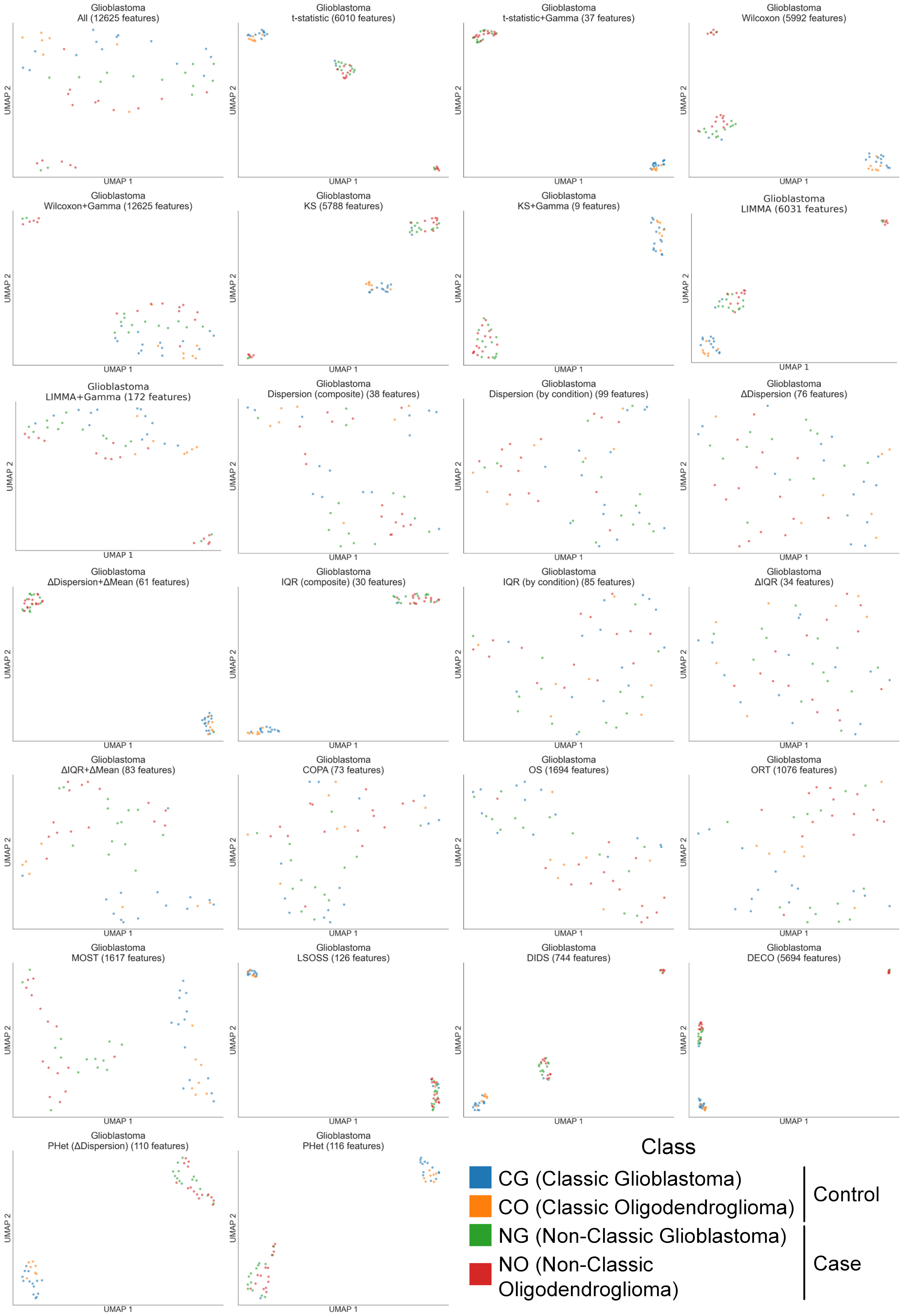


**Supplementary Figure 14:** UMAP visualizations of malignant glioma and oligodendroglioma patients from the glioblastoma microarray gene expression dataset (Glioblastoma) [21] using all and selected features by each method, colored by original patient types (Supplementary Table 4).

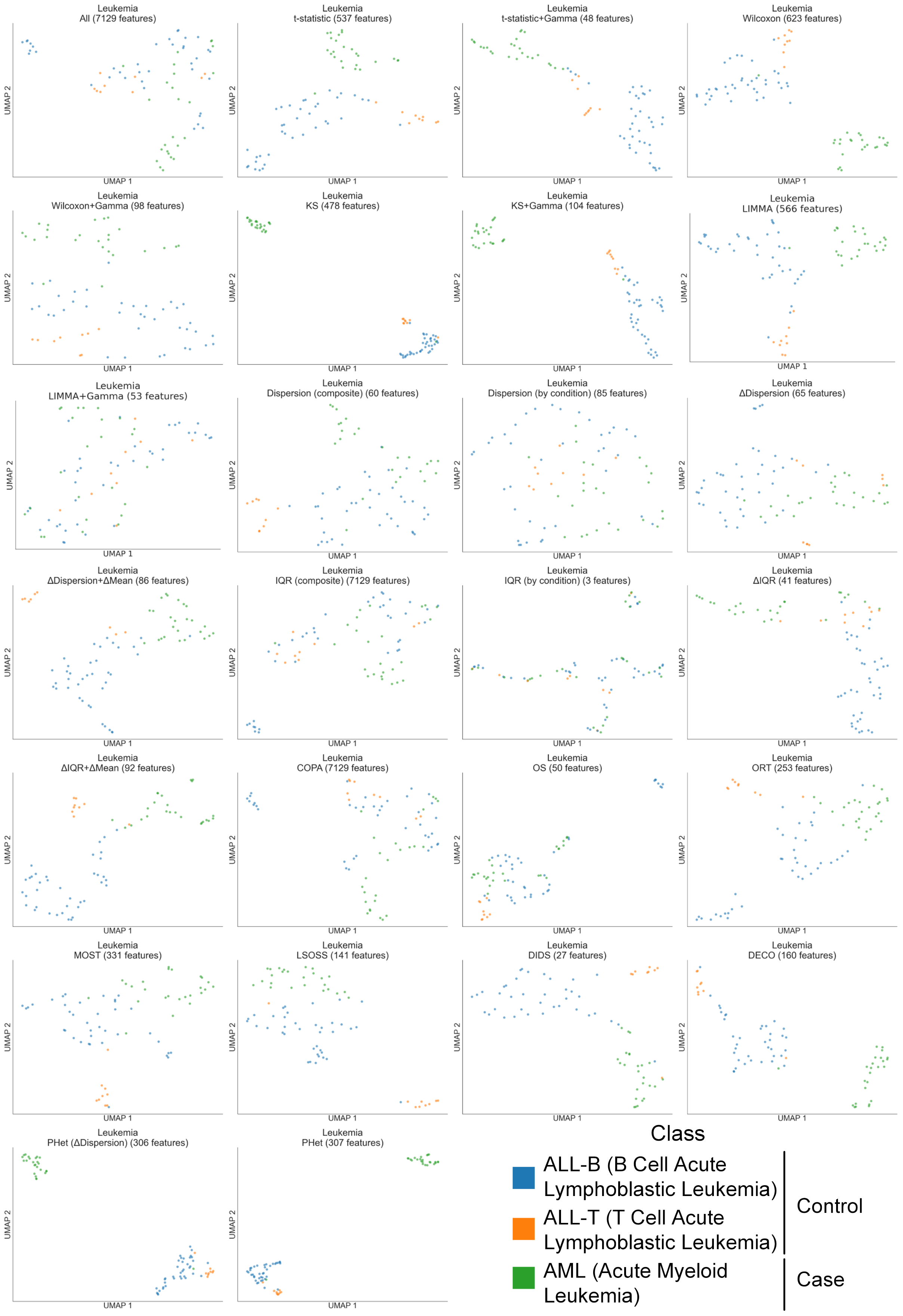


**Supplementary Figure 15:** UMAP visualizations of the leukemia microarray gene expression dataset (Leukemia) [14] using all and selected features by each method, colored by original patient types (Supplementary Table 4).

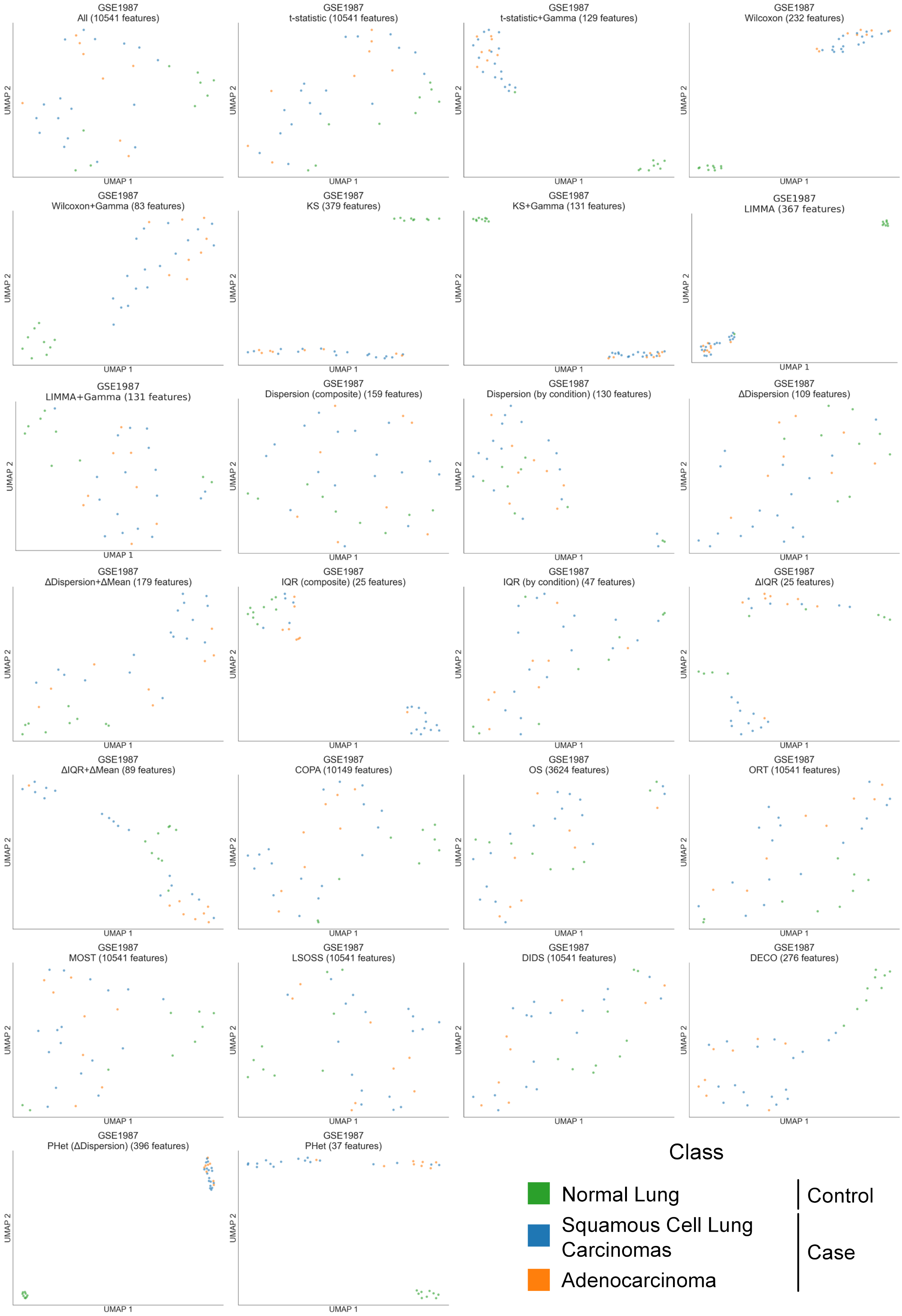


**Supplementary Figure 16:** UMAP visualizations of the lung microarray gene expression dataset (GSE1987) [11] with adenocarcinoma, squamous cell carcinoma and normal lung tissue samples using all and selected features by each method, colored by original patient types (Supplementary Table 4).

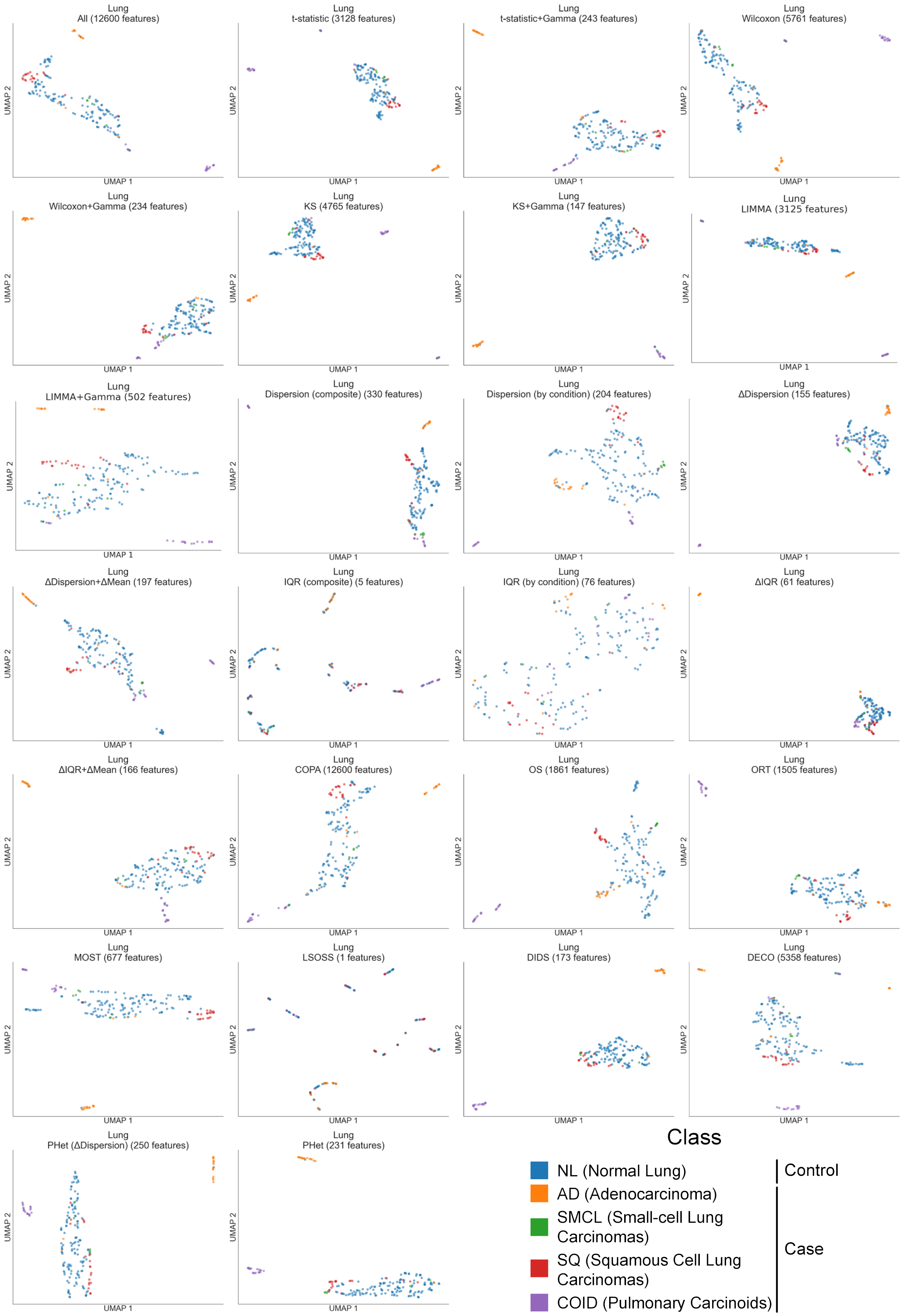


**Supplementary Figure 17:** UMAP visualizations of four lung cancer types (adenocarcinoma, small-cell lung carcinomas, squamous cell lung carcinomas, and pulmonary carcinoids) and normal lung tissue samples from the lung microarray gene expression dataset (Lung) [3] using all and selected features by each method, colored by original patient types (Supplementary Table 4).

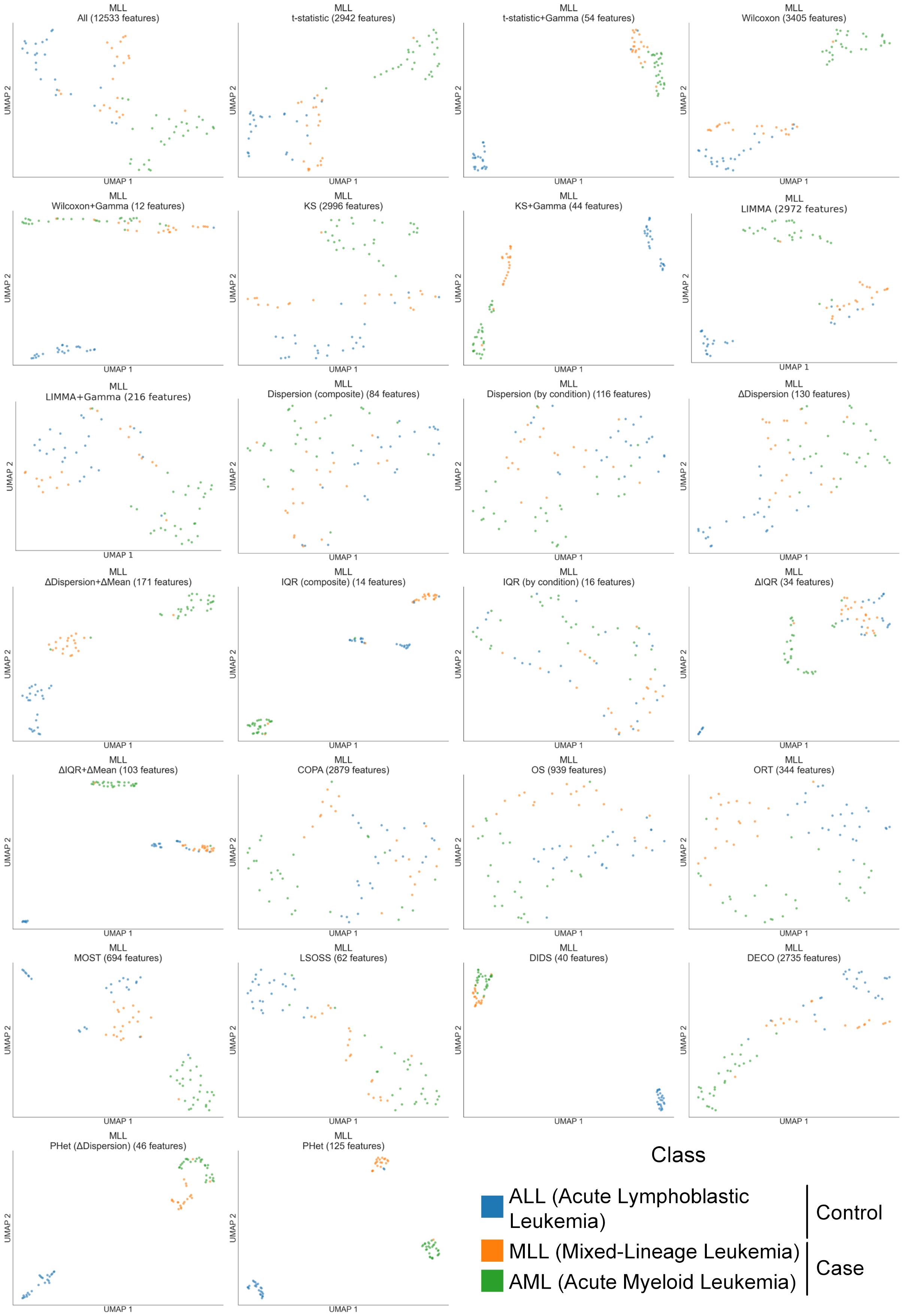


**Supplementary Figure 18:** UMAP visualizations of three subtypes of leukemia (acute lymphoblastic, mixed-lineage, and acute myeloid) from the mixed-lineage leukemia microarray gene expression dataset (MLL) [1] using all and selected features by each method, colored by original patient types (Supplementary Table 4).

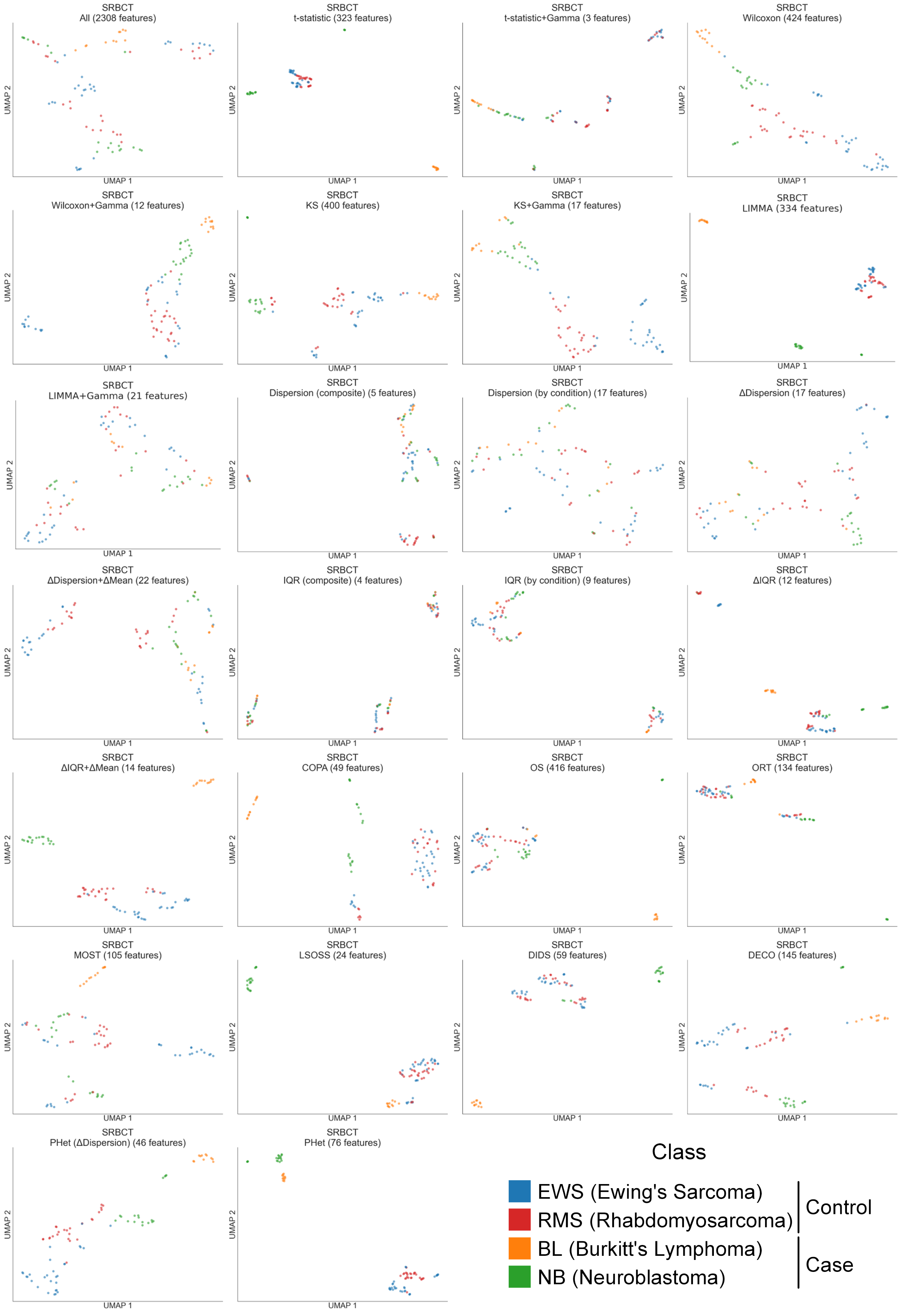


**Supplementary Figure 19:** UMAP visualizations of the small round blue cell tumor patients from SRBCT microarray gene expression dataset [18] using all and selected features by each method, colored by original patient types (Supplementary Table 4).

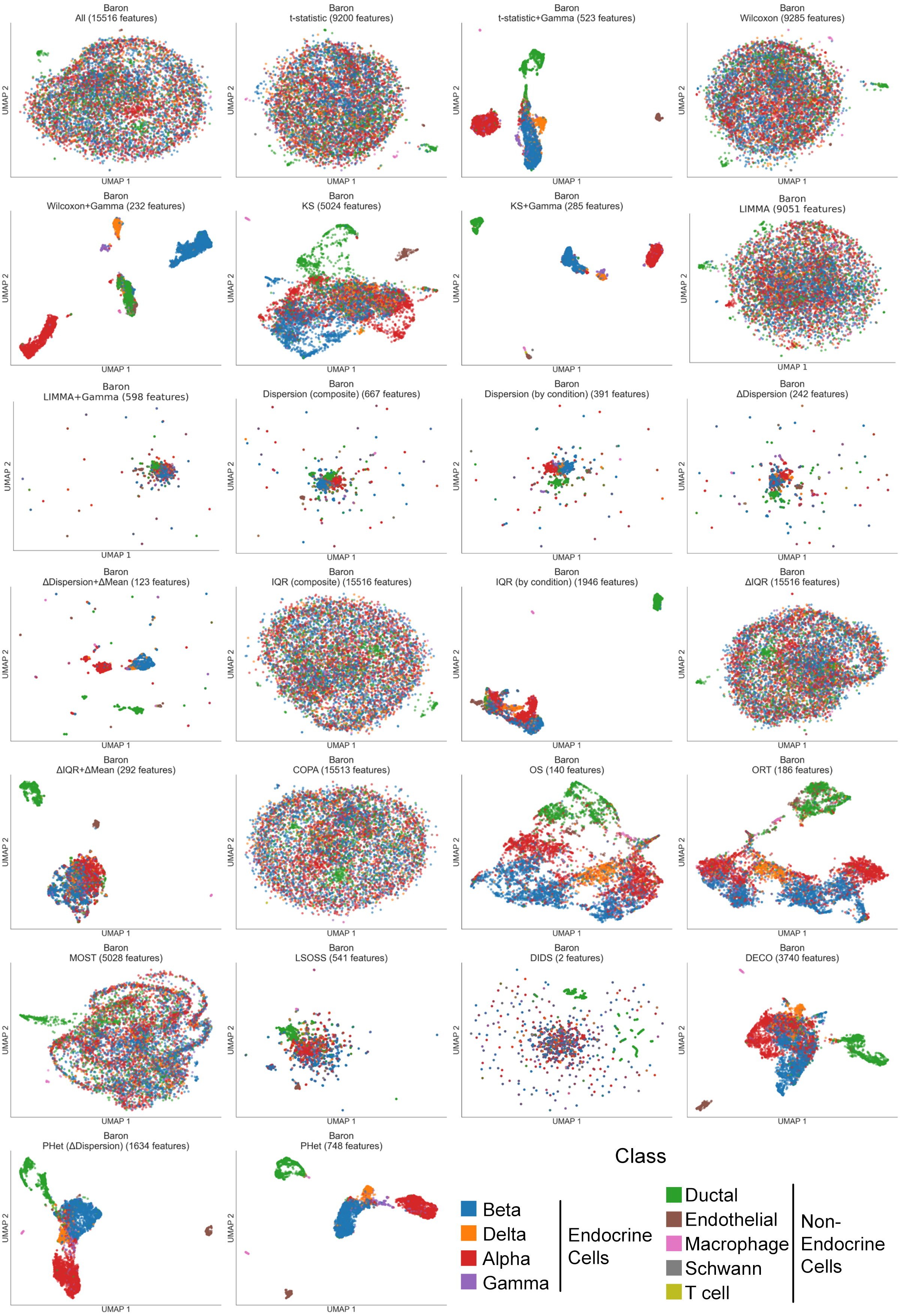


**Supplementary Figure 20:** UMAP visualizations of human pancreas cell types from the Baron single-cell transcriptomic dataset [2] using all and selected features by each method, colored by original cell types (Supplementary Table 5).

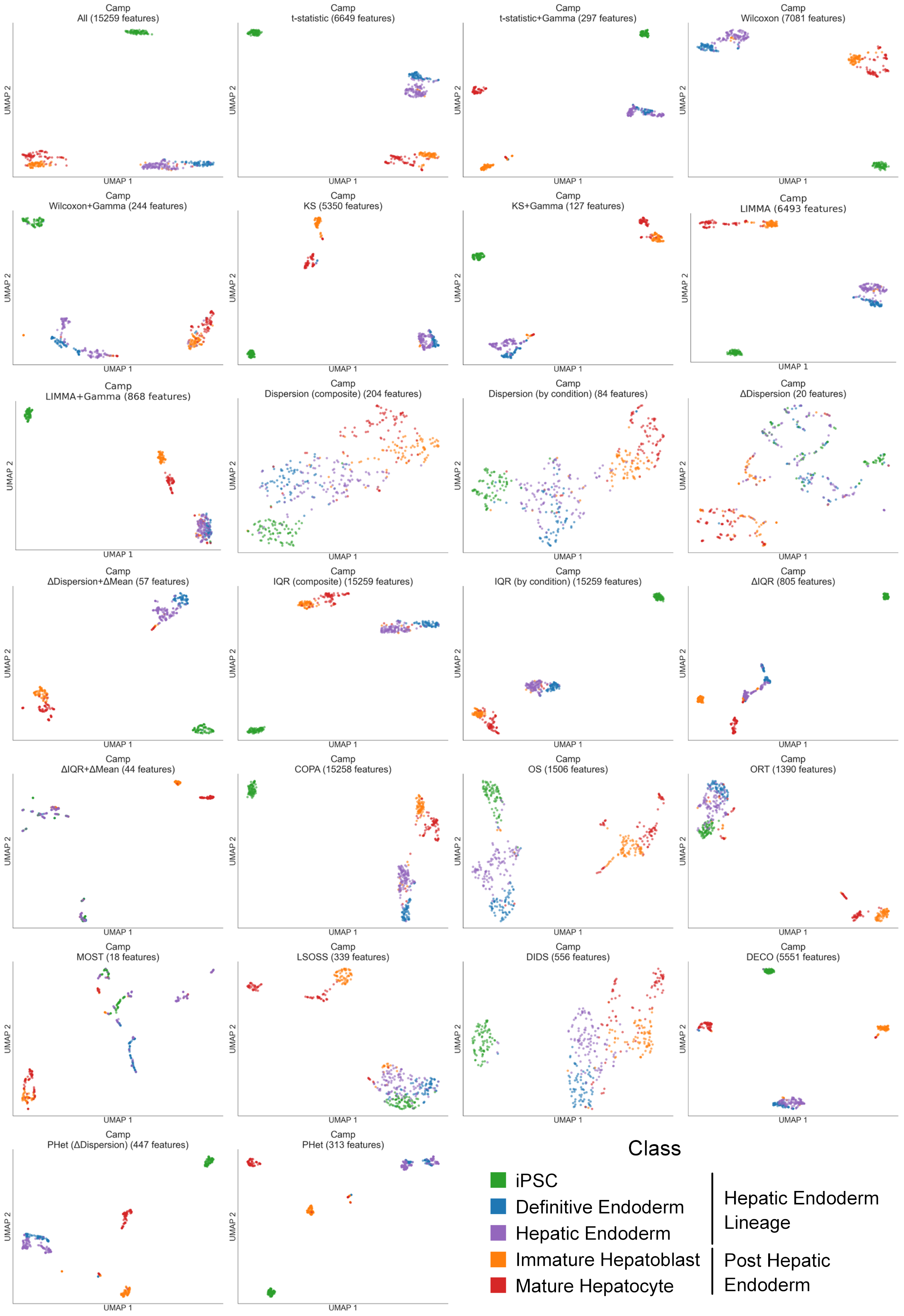


**Supplementary Figure 21:** UMAP visualizations of cells during the hepatocyte differentiation in fetal and adult liver from the Camp single-cell transcriptomic dataset [5] using all and selected features by each method, colored by original cell types (Supplementary Table 5).

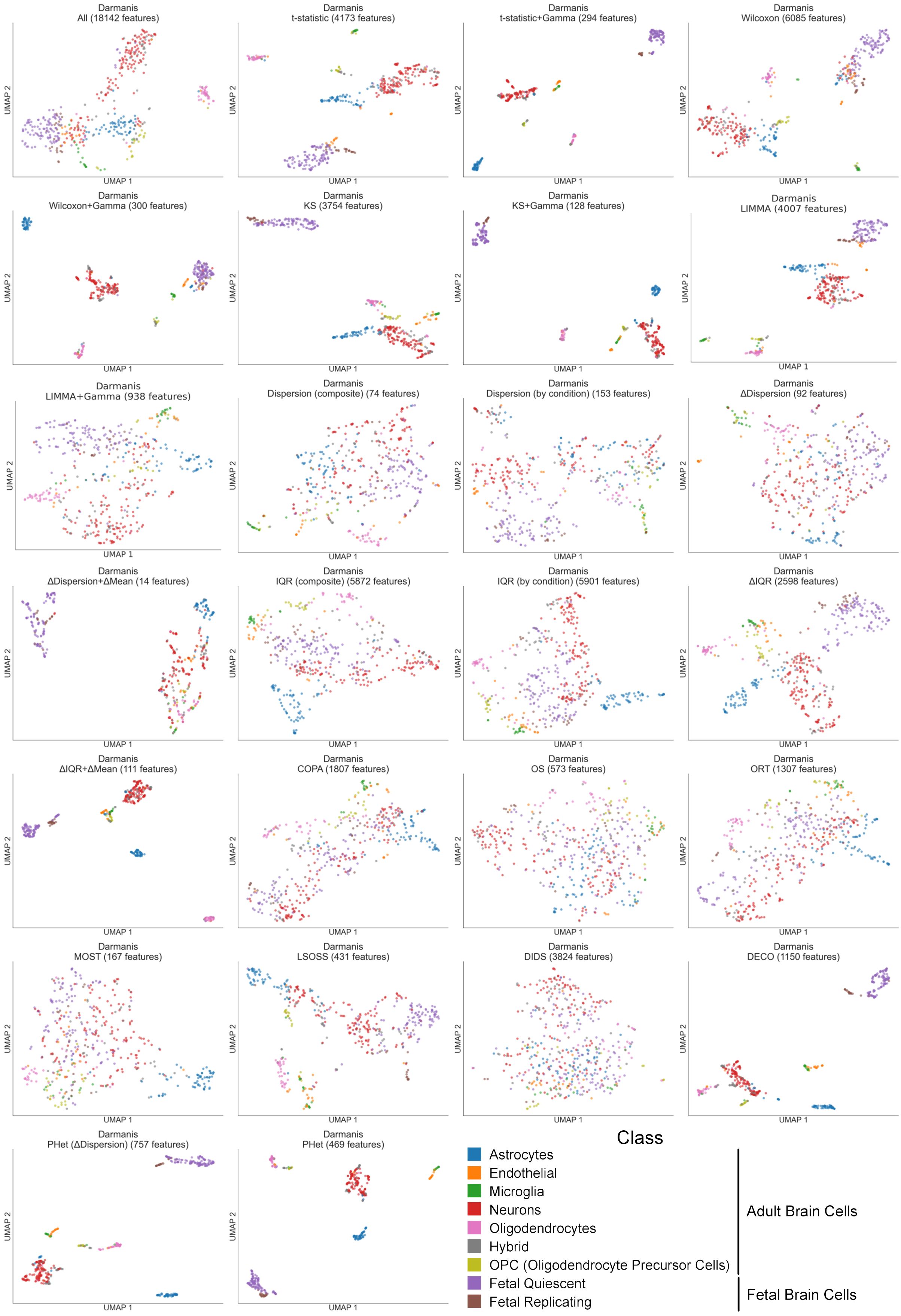


**Supplementary Figure 22:** UMAP visualizations of adult and fetal human brain cells from the Darmanis single-cell transcriptomic dataset [9] using all and selected features by each method, colored by original cell types (Supplementary Table 5).

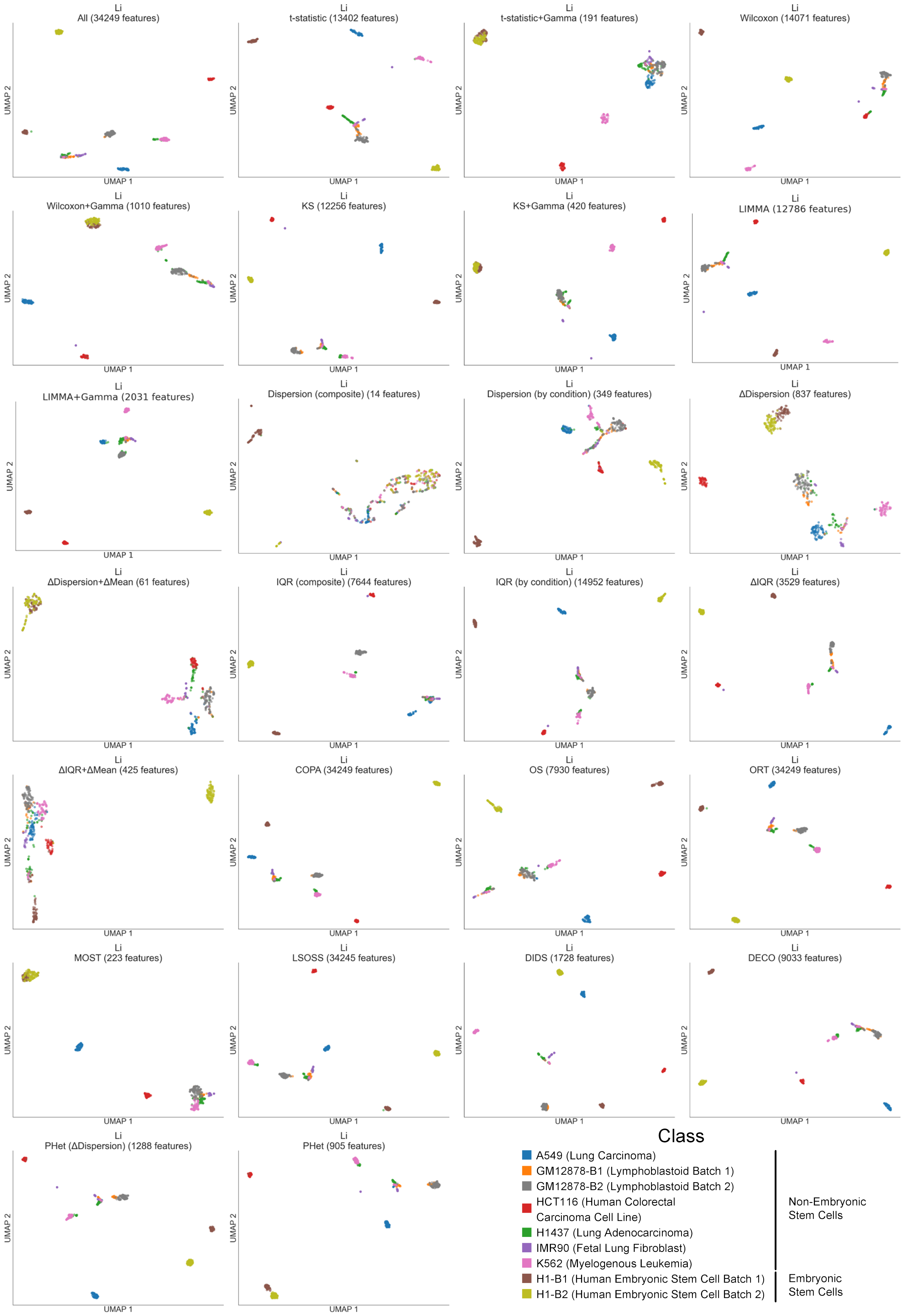


**Supplementary Figure 23:** UMAP visualizations of two batches of human embryonic stem & lymphoblastoid cells and several tumor tissues from patients with colorectal and lung cancers from the Li single-cell transcriptomic dataset [20] using all and selected features by each method, colored by original cell types (Supplementary Table 5).

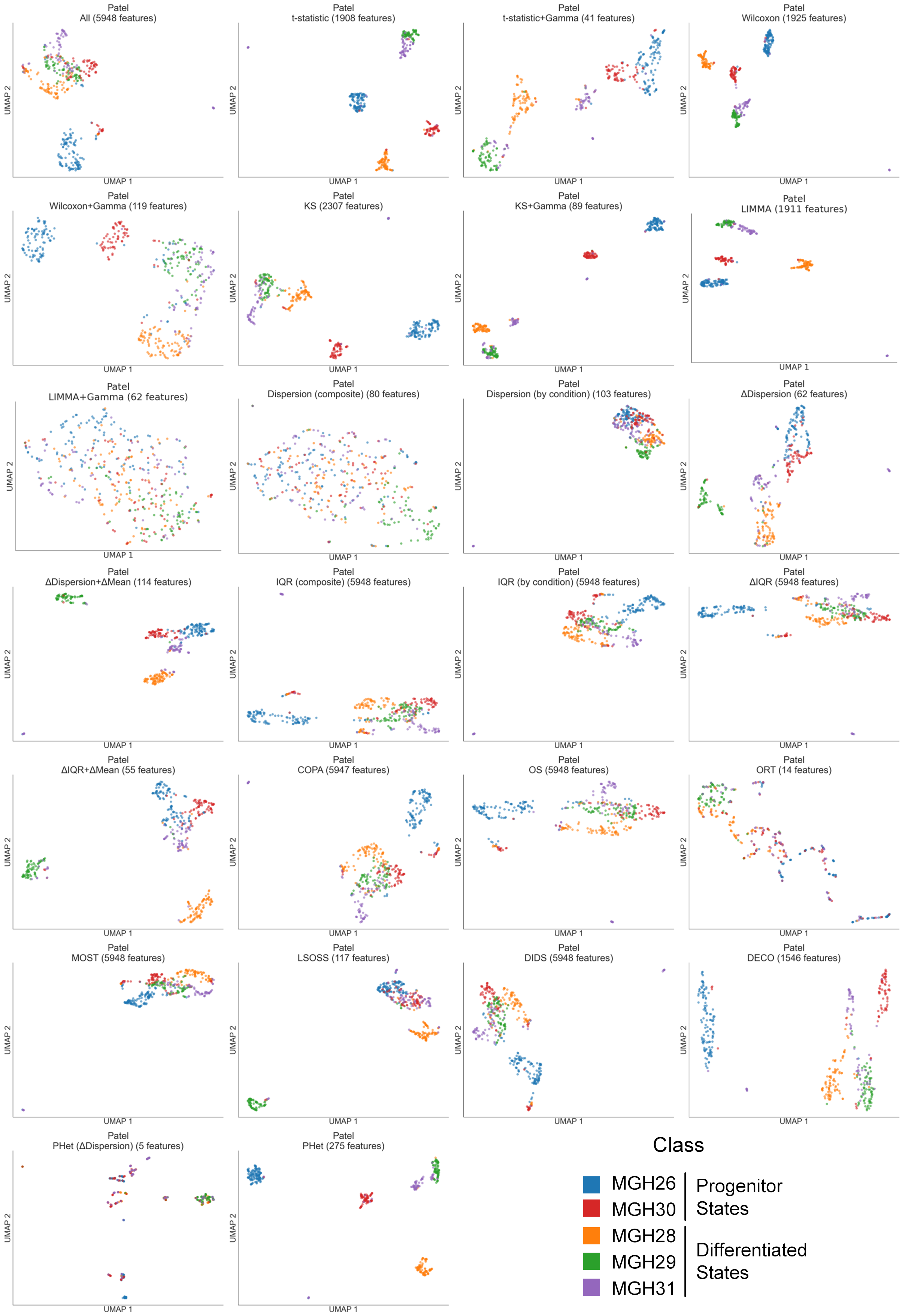


**Supplementary Figure 24:** UMAP visualizations of primary human glioblastomas from the Patel single-cell transcriptomic dataset [22] using all and selected features by each method, colored by original cell types (Supplementary Table 5).

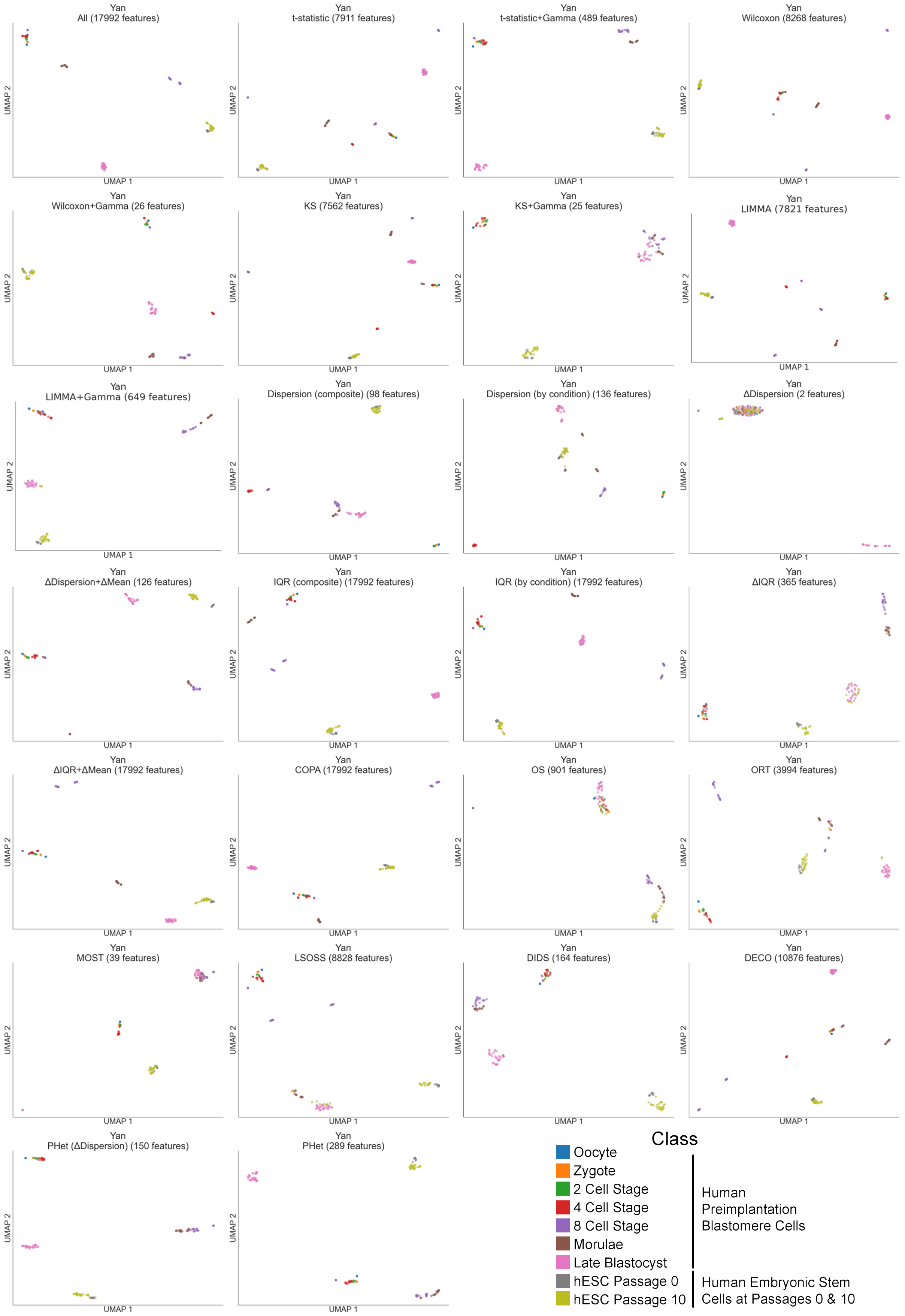


**Supplementary Figure 25:** UMAP visualizations of cells corresponding human preimplantation embryos and human embryonic stem cells at different stages from the Yan single-cell transcriptomic dataset [32] using all and selected features by each method, colored by original cell types (Supplementary Table 5).

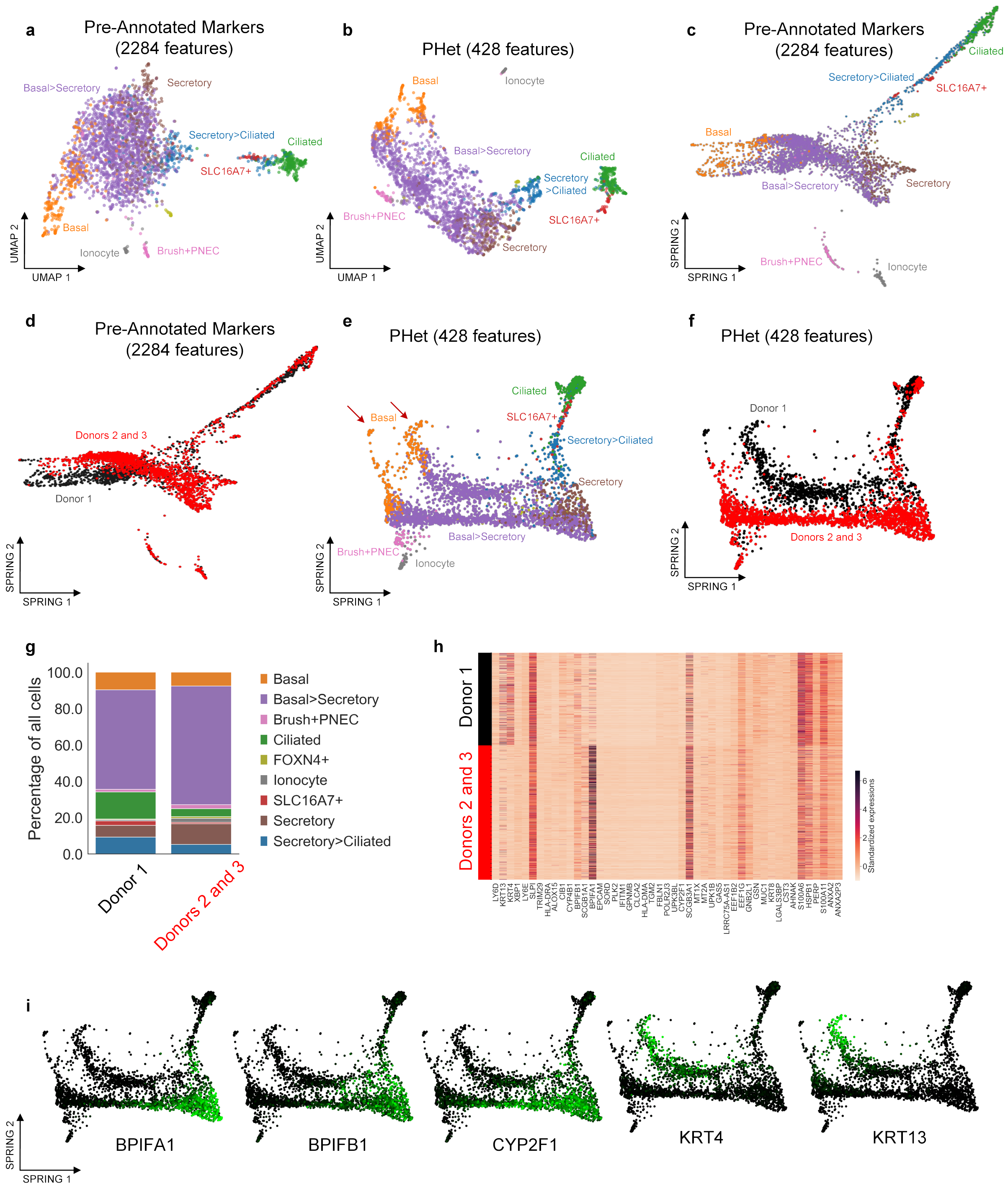


**Supplementary Figure 26:** HBECs single-cell transcriptomics data reveal distinct expression profiles. **a**,**b**,**c**,**e**, UMAP visualizations using the pre-annotated markers and PHet's selected features and their corresponding SPRING visualizations, respectively. Coloring represents the previously annotated cell types. The trajectories are visually represented by red colored arrows. **d**, A SPRING plot using pre-annotated markers displaying two types of donors: Donor 1 (black) and Donors 2 and 3 (red). However, the plot does not reveal clear and distinctive trajectories among the different donor types. **f**, A SPRING plots using PHet's features displaying two distinct trajectories: Donor 1 (black) and Donors 2 and 3 (red). **g**, A bar plot representing the relative abundance of cell types grouped by donors. **h**, A heatmap demonstrating the top differentially expressed features for the two identified trajectories. **i**, SPRING plots of the selected top features selected by PHet. The color gradient from black to green indicated cells enriched with the corresponding feature.

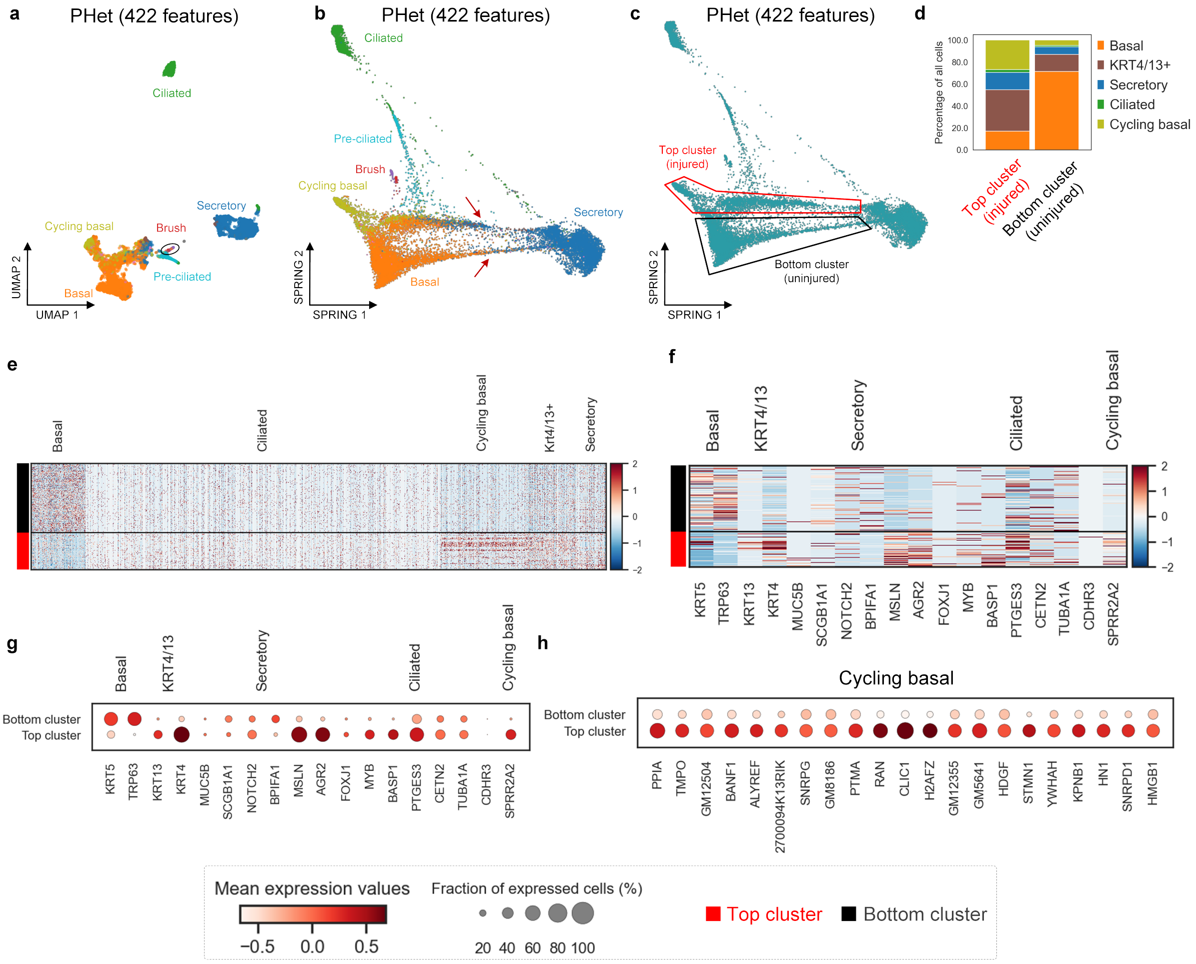


**Supplementary Figure 27:** MTECs single-cell transcriptomics data of uninjured and regenerating mice using PHet's selected features. **a**, The UMAP visualization of PHet's selected features (**a**). Coloring represents the previously annotated cell types. **b**, The SPRING visualizations of PHet's selected features to explore trajectories of cells. **c**, Two distinct clusters are displayed in the SPRING plot, with one cluster (injured) located at the top and another (uninjured) at the bottom. **d**, The relative abundance of cells between these clusters are shown as a bar plot. **e**,**f**,**g**,**h**, Heat maps and dot plots of lineage-specific markers between the top and bottom clusters. The size of the circles in dot plots represented the fractions of cells expressing a specific marker in each cluster.

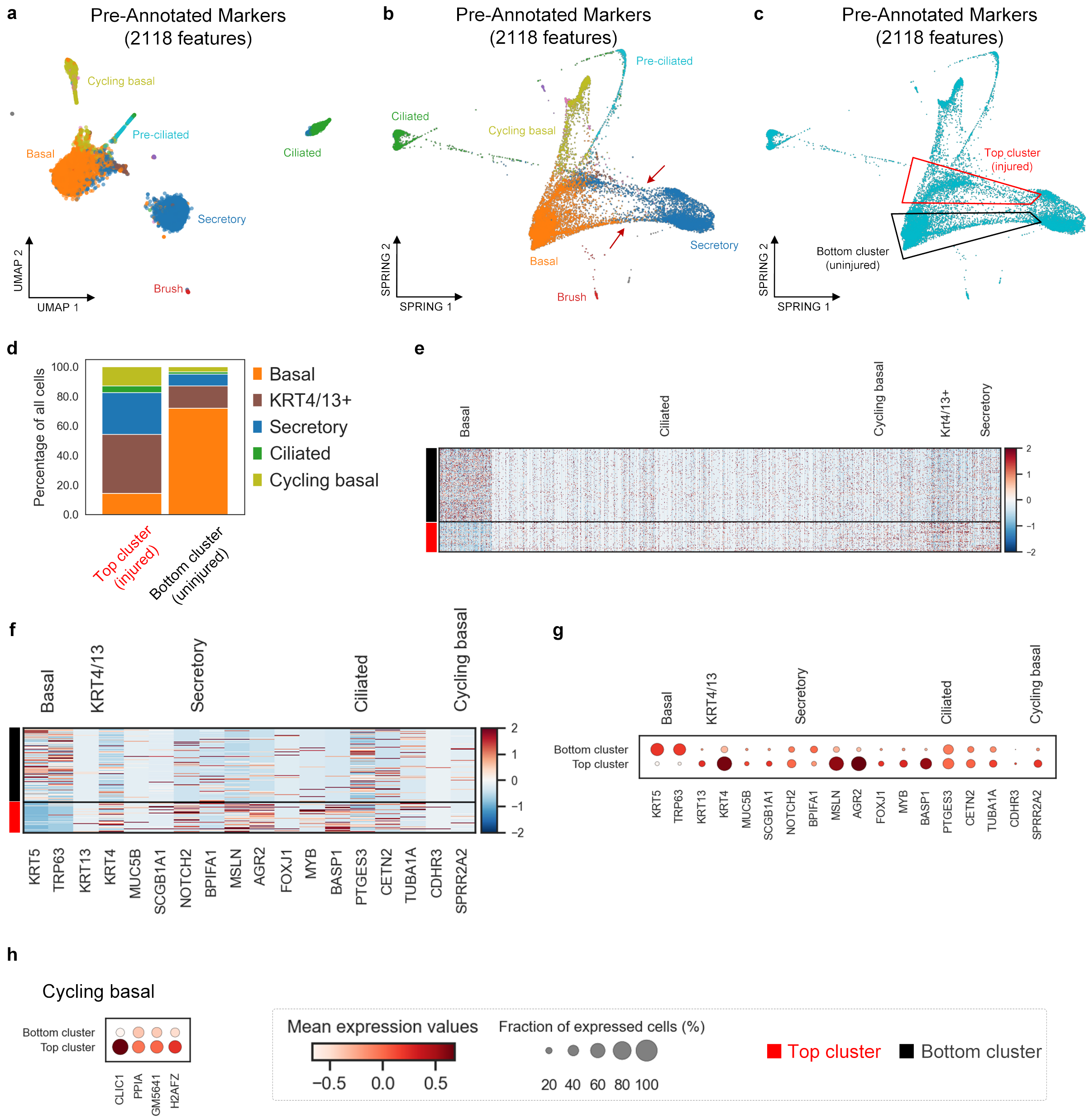


**Supplementary Figure 28:** MTECs single-cell transcriptomics data of uninjured and regenerating mice using pre-annotated markers. **a**, The UMAP visualization of pre-annotated markers (**a**). Coloring represents the previously annotated cell types. **b**, The SPRING visualizations of pre-annotated markers to explore trajectories of cells. **c**, Two clusters are displayed in the SPRING plot, with one cluster (injured) located at the top and another (uninjured) at the bottom. **d**, The relative abundance of cells between these clusters are shown as a bar plot. **e**,**f**,**g**,**h**, Heat maps and dot plots of lineage-specific markers between the top and bottom clusters. The size of the circles in dot plots represented the fractions of cells expressing a specific marker in each cluster.

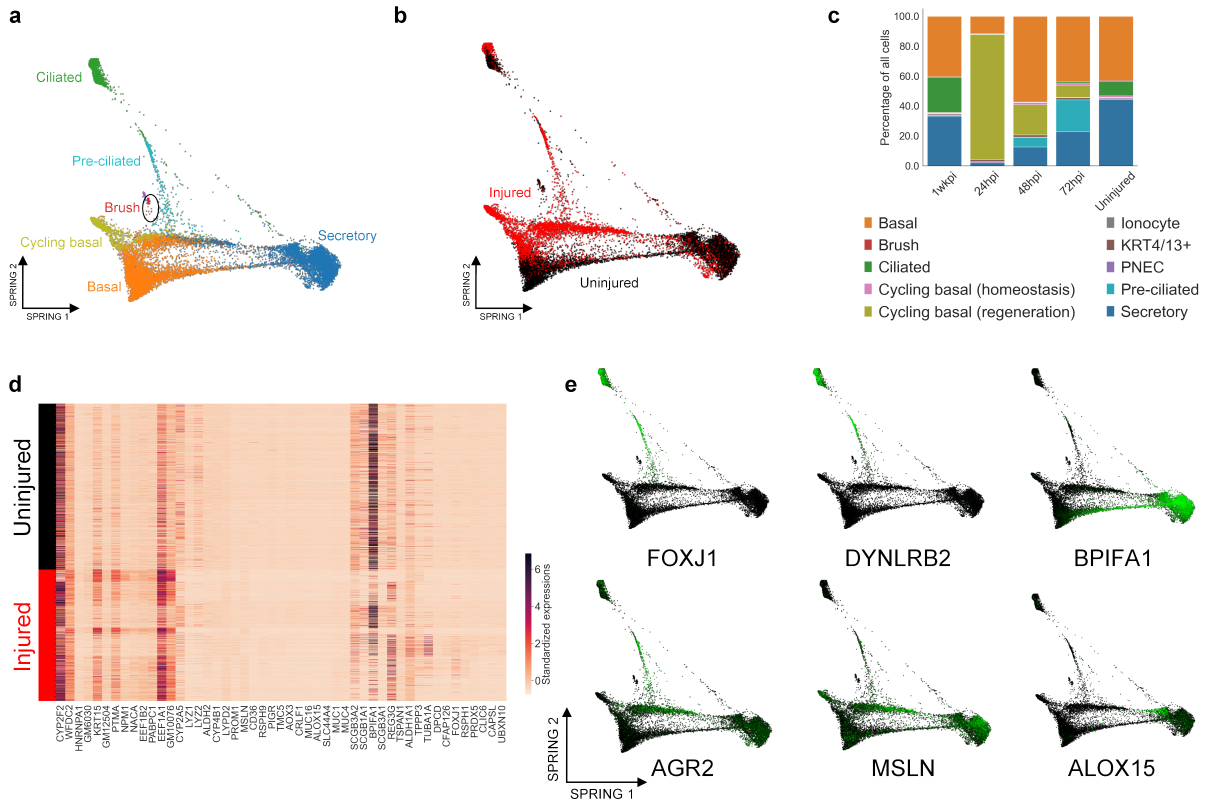


**Supplementary Figure 29:** Lineage tracing of MTECs single-cell transcriptomics data. **a**,**b**, SPRING plots of MTECs data showing cells from uninjured and regenerating mice. The first trajectory represents the healthy mice (black-colored cells) that are enriched in the BPIFA1 gene responsible for regulating secreted protein while the second upper red-colored trajectory represents regenerating mice at 1, 2, 3, and 7 days post-injury (dpi). **c**, The relative abundance of cell types at each time point (reproduced from Plasschaert and colleagues paper [24]). At 2 and 3 dpi, secretory cells reappeared, and by 7 dpi the relative abundance of cell populations is shown to exhibit similar to that seen in uninjured tracheae. **d**, Differential analysis between uninjured and regenerating mice. **e**, SPRING plots of the selected top features selected by PHet. The color gradient from black to green indicated cells enriched with the corresponding feature.

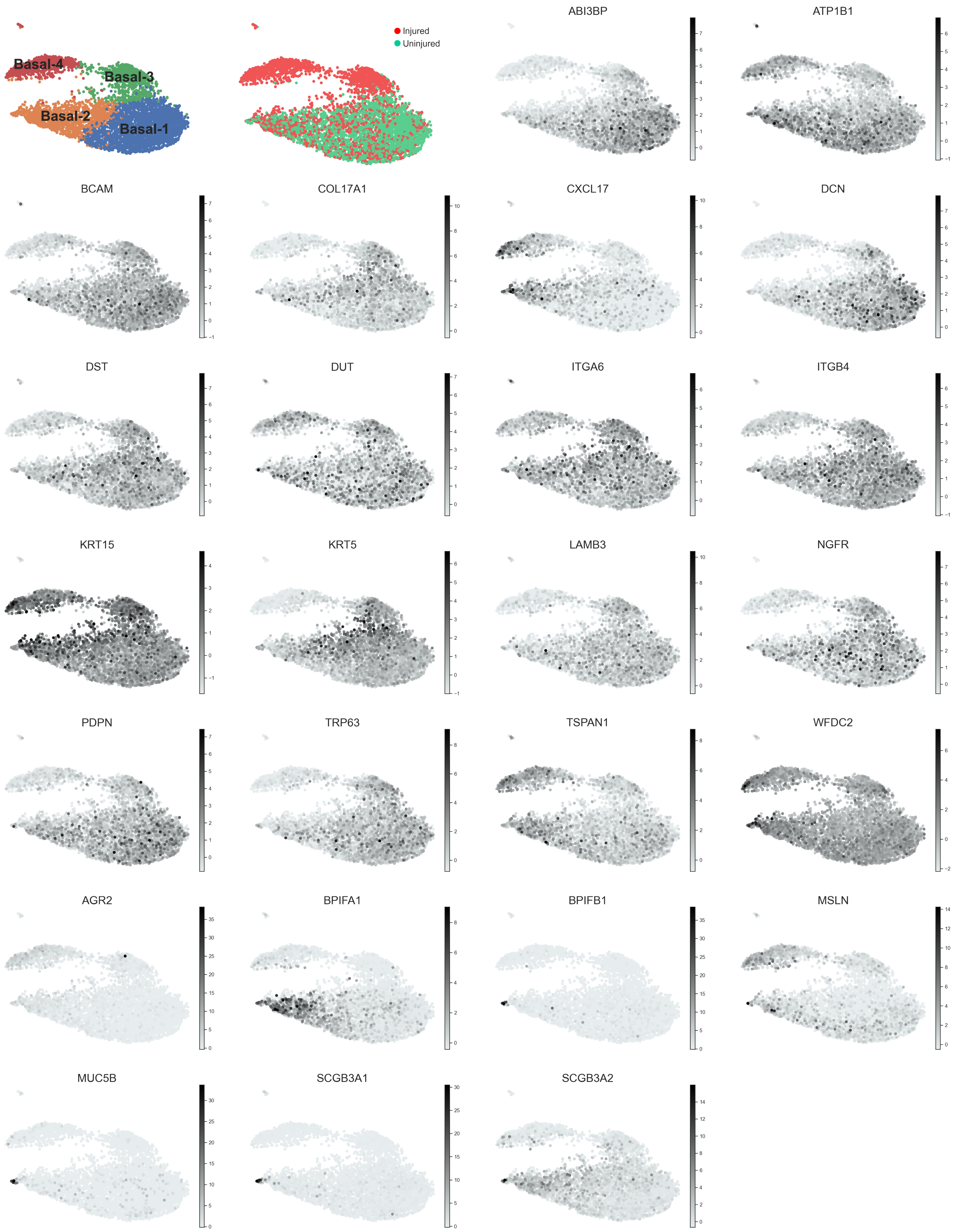


**Supplementary Figure 30:** UMAP visualizations of four subpopulations of basal cells within MTECs using PHet's selected features (422). Based on expression patterns of features selected by PHet, four distinct subsets of basal cells are identified: Basal-1, Basal-2, Basal-3, and Basal-4. Injured vs uninjured conditions using PHet's selected features is shown in the second figure of the first row. The first five rows corresponded to basal signatures, ranging from ABI3BP to WFDC2, while the last two rows represented the secretory gene signatures, from AGR2 to SCGB3A2.

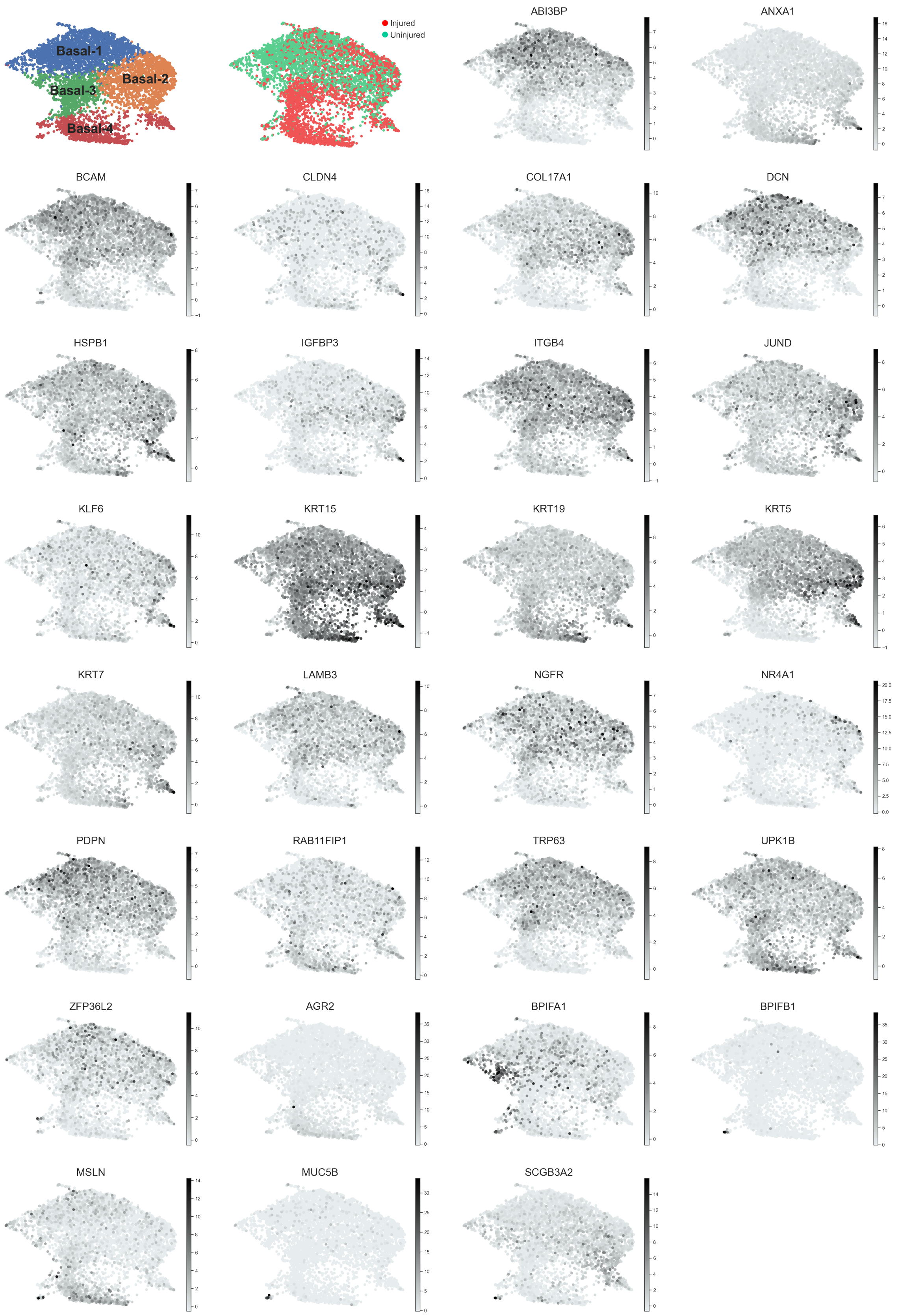


**Supplementary Figure 31:** UMAP visualizations of four subpopulations of basal cells within MTECs using pre-annotated markers (361). Based on expression patterns of markers, four subsets of basal cells are identified: Basal-1, Basal-2, Basal-3, and Basal-4. Injured vs uninjured conditions using pre-annotated markers is shown in the second figure of the first row. The first six rows corresponded to basal signatures, ranging from ABI3BP to ZFP36L2, while the last two rows represented the secretory gene signatures, from AGR2 to SCGB3A2.

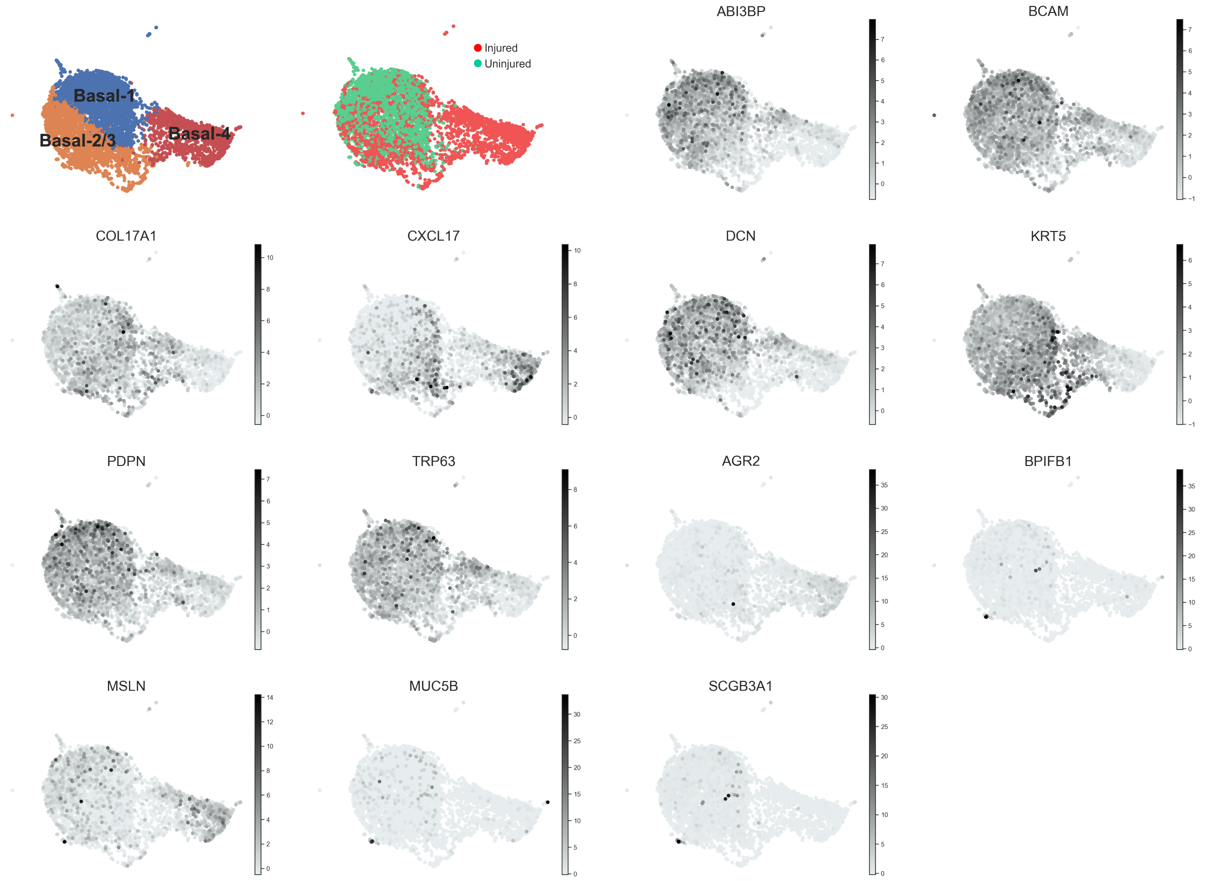


**Supplementary Figure 32:** UMAP visualizations of three subpopulations of basal cells within MTECs using HV features (5580). Based on expression patterns of HV features, three subsets of basal cells are identified: Basal-1, Basal-2/3, and Basal-4. Injured vs uninjured conditions using HV features is shown in the second figure of the first row. From the third column of the first row, the 8 figures corresponded to basal signatures, ranging from ABI3BP to TRP63, while the last two rows represented the secretory gene signatures, from AGR2 to SCGB3A1.

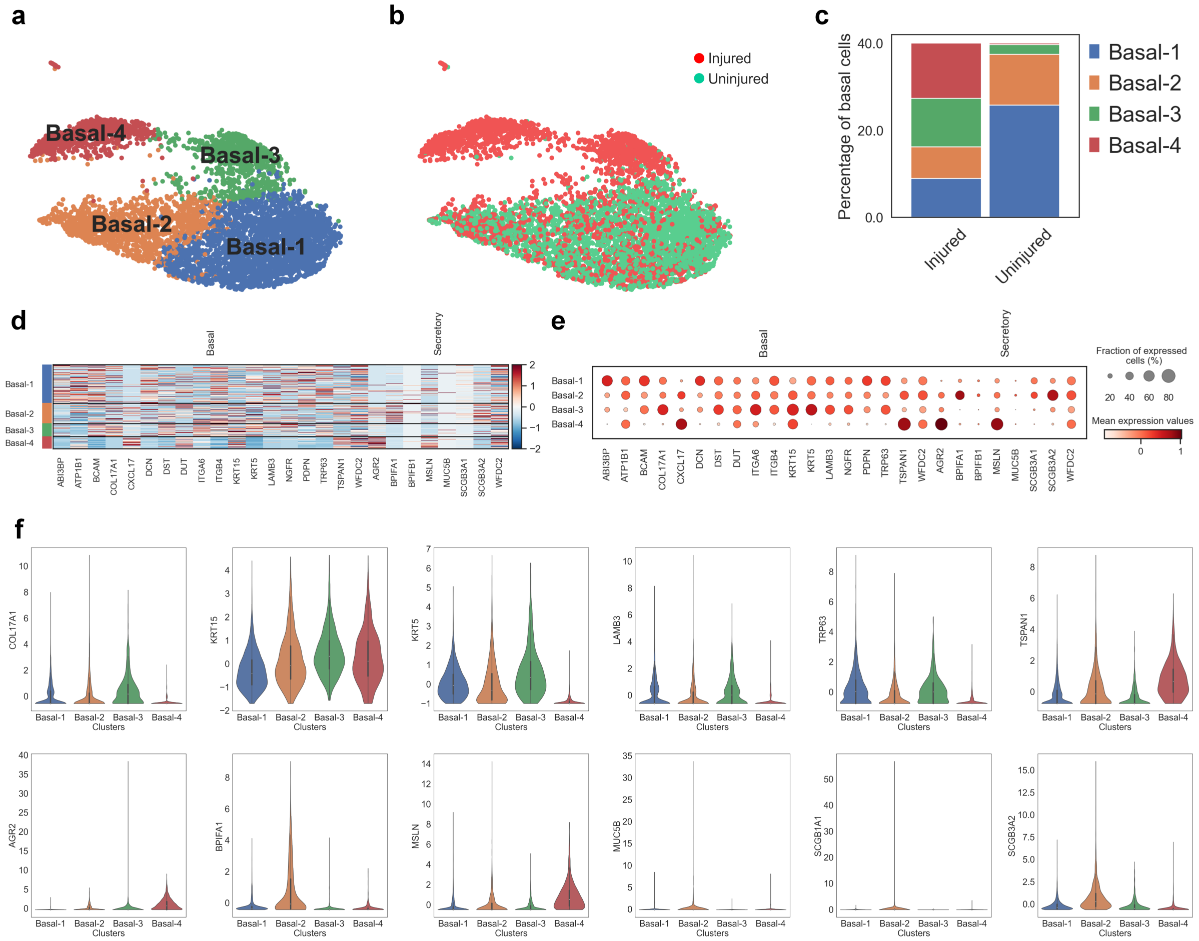


**Supplementary Figure 33:** Expression profiles of the four basal subsets using PHet's selected features (422). **a**,**b**, UMAP visualizations of the four basal subpopulations and injured vs uninjured conditions using PHet's selected features. **c**, The barplot shows the distribution of basal cells among injured and uninjured mice. **d**,**e**, A heat map and a dot plot analysis of the expression levels of basal and secretory cell markers among the four basal subsets. The size of the circles in the dot plot indicates the fractions of cells expressing a certain marker for each cell type. These plots reveal that Basal-1 and Basal-3 cell populations are enriched with canonical basal cell markers, such as KRT5 and TPR63. On the other hand, Basal-2 and Basal-4 cells show enrichment for SCGB3A2 and TSPAN1, respectively, suggesting that they represent separate basal cells transitioning to a luminal secretory phenotype. These results suggest that the four basal subsets have different functional roles and molecular characteristics in the airway epithelium. **f**, Violin plots of the distribution of features expressions of basal and secretory cells' markers among the four basal cell subpopulations. The top row of violin plots shows the distribution of expression of six basal cell markers: COL17A1, KRT15, KRT5, LAMB3, TRP63, and TSPAN1. The bottom row shows the distribution of expression of six secretory cell markers: AGR2, BPIFA1, MSLN, MUC5B, SCGB1A1, and SCGB3A2. The plots reveal that Basal-4 cells have higher expression in AGR2 and MSLN secretory cells gene signatures. AGR2 and MSLN encode proteins involved in cell adhesion and secretion, respectively. The higher expression of these genes in Basal-4 cells reassures that they are indeed transitioning to a secretory state.

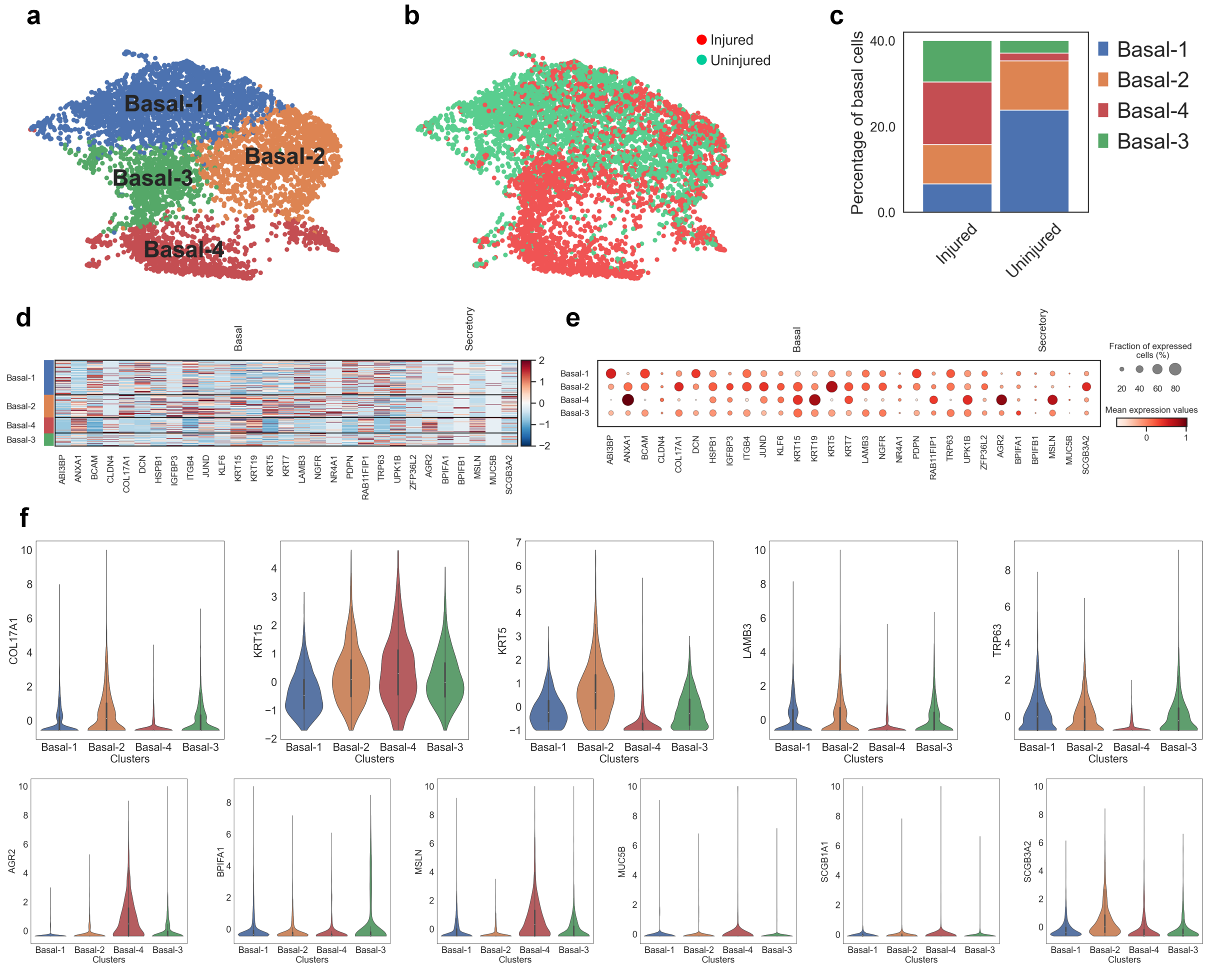


**Supplementary Figure 34:** Expression profiles of the four basal subsets using pre-annotated markers (361). **a**,**b**, UMAP visualizations of the four basal subpopulations and injured vs uninjured conditions using pre-annotated markers. **c**, The barplot shows the distribution of basal cells among injured and uninjured mice. **d**,**e**, A heat map and a dot plot analysis of the expression levels of basal and secretory cell markers among the four basal subsets. The size of the circles in the dot plot indicates the fractions of cells expressing a certain marker for each cell type. These plots revealed that Basal-1, Basal-2, and Basal-3 cell populations are enriched with canonical basal cell markers (e.g., KRT5 and TPR63). **f**, Violin plots of the distribution of features expressions of basal and secretory cells' markers among the four basal cell subpopulations. The top row of violin plots shows the distribution of expression of five basal cell markers: COL17A1, KRT15, KRT5, LAMB3, and TRP63. The bottom row shows the distribution of expression of six secretory cell markers: AGR2, BPIFA1, MSLN, MUC5B, SCGB1A1, and SCGB3A2. The plots reveal that Basal-4 cells have higher expression in AGR2 and MSLN secretory cells gene signatures. AGR2 and MSLN encode proteins involved in cell adhesion and secretion, respectively. The higher expression of these genes in Basal-4 cells reassures that they are indeed transitioning to a secretory state.

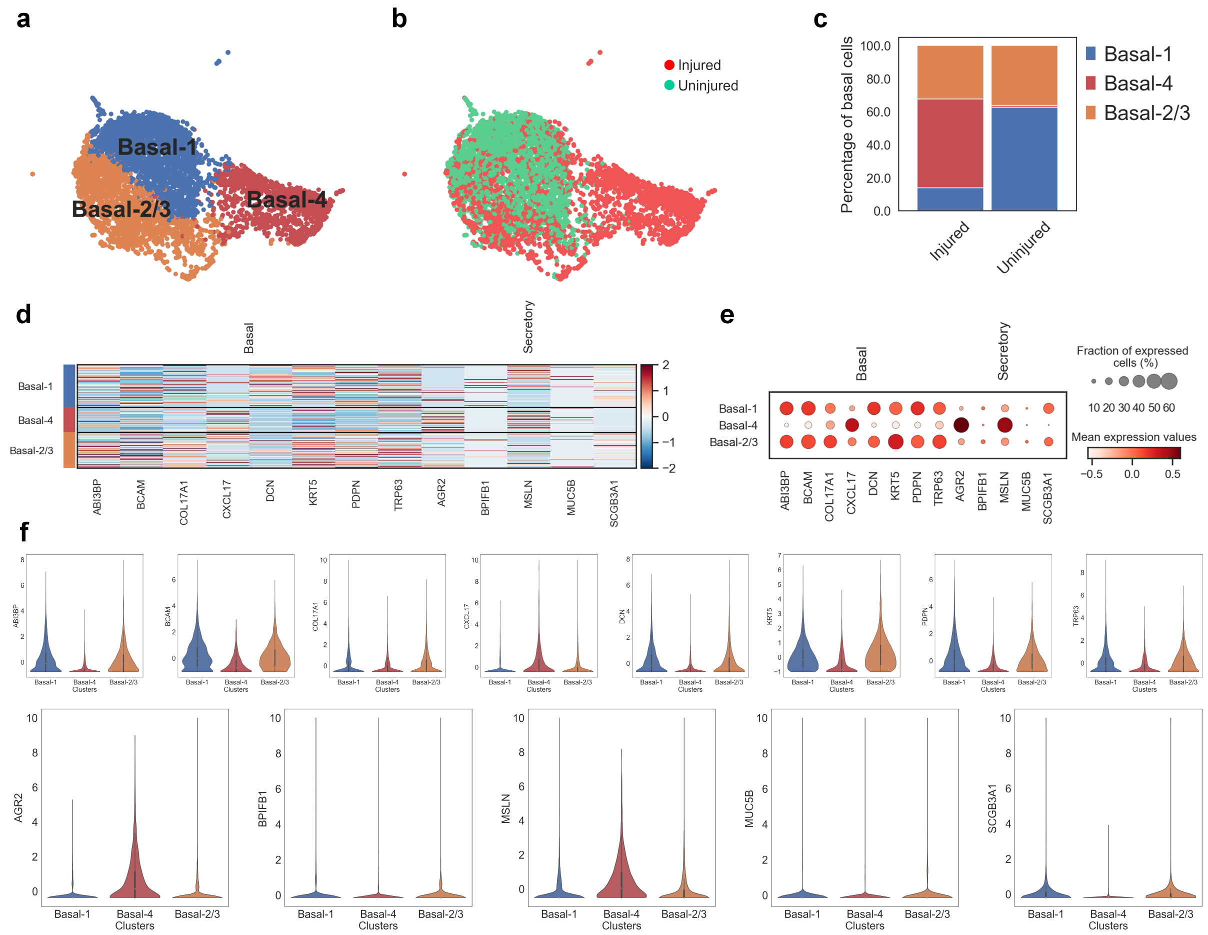


**Supplementary Figure 35:** Expression profiles of the three basal subsets using HV features (5580). **a**,**b**, UMAP visualizations of the three basal subpopulations and injured vs uninjured conditions using HV features identification. **c**, The barplot shows the distribution of basal cells among injured and uninjured mice. **d**,**e**, A heat map and a dot plot analysis of the expression levels of basal and secretory cell markers among the three basal subsets. The size of the circles in the dot plot indicates the fractions of cells expressing a certain marker for each cell type. These plots revealed that Basal-1 and Basal-2/3 cell populations are enriched with canonical basal cell markers (e.g., KRT5 and TPR63) while cells in Basal-4 showed enrichment for MSLN. **f**, Violin plots of the distribution of features expressions of basal and secretory cells' markers among the four basal cell subpopulations. The top row of violin plots shows the distribution of expression of 8 basal cell markers: ABI3BP, BCAM, COL17A1, CXCL17, DCN, KRT5, PDPN, and TRP63. The bottom row shows the distribution of expression of five secretory cell markers: AGR2, BPIFB1, MSLN, MUC5B, and SCGB1A1. The violin plots reveal that Basal-4 cells have higher expression in AGR2 and MSLN secretory cells gene signatures. AGR2 and MSLN encode proteins involved in cell adhesion and secretion, respectively. The higher expression of these genes in Basal-4 cells reassures that they are indeed transitioning to a secretory state.

### Supplementary Note 1: Hyperparameter selection for PHet

We evaluated the PHet performance under different hyperparameter settings. The main hyperparameters of PHet consist of the binning weights ($\mathbf{w}$) and the alpha cutoff threshold ($\alpha \in (0,1)$). To determine the optimal configurations for these hyperparameters, we conducted experiments using test datasets (Supplementary Tables 1 and 2) comprising six microarray (BCCA [16], DLBCL [27], GSE2191 [31], GSE2535 [8], MDS [23], and Prostate [28]) and four single-cell transcriptomics (scRNA-seq) (Knoblich [4], Lake [19], Segerstolpe [26], and Wang [30]) datasets. The assessment focused on evaluating the influence of different weight combinations and alpha values on the selection of features that contribute to robust clustering outcomes, as measured by the adjusted Rand index, adjusted mutual information, homogeneity, completeness, and V-measure. Spectral clustering was employed as the clustering method for these experiments. We systematically explored various weight combinations, including $\mathbf{w} \in \{(0.4, 0.3, 0.2, 0.1), (0.3, 0.4, 0.1, 0.2), (0.3, 0.3, 0.2, 0.2), (0.2, 0.1, 0.3, 0.4), (0.1, 0.2, 0.4, 0.3), (0.1, 0.2, 0.3, 0.4), (0.2, 0.2, 0.3, 0.3)\}$, in conjunction with alpha values at $\alpha \in \{0.01, 0.05, 0.1\}$. This exhaustive approach allowed for a comprehensive grid search and meticulous analysis of clustering results for each configuration. The optimal configuration was identified based on its ability to consistently yield a small set of features with high ARI scores across all datasets.

The tests showed that maintaining an alpha value of 0.01 across all datasets resulted in the lowest average number of predicted features ($\approx$ 507.45 features) (Fig. 36), and led to improved clustering outcomes with an average ARI score of 50.01%. Further investigation into the predicted features and ARI scores across different weights revealed that using a set of weights $\{(0.4, 0.3, 0.2, 0.1), (0.2, 0.1, 0.3, 0.4)\}$ at alpha 0.01 resulted the lowest number of features on average, with 395.1 and 492.5 features respectively, and achieved the best ARI scores of 61.02% and 52.30%, respectively (Fig. 36). Interestingly, when the alpha value was increased to 0.05 and 0.1, the weights $\mathbf{w} = (0.4, 0.3, 0.2, 0.1)$ continued to outperform other weights in terms of ARI scores, achieving 50.06% and 34.77%, respectively. It is worth noting that ARI scores were generally higher for transcriptomics data compared to microarray data, with the latter exhibiting ARI scores over 10% lower at alpha 0.01 (Fig. 36).

These findings suggest that users should consider adjusting the alpha value or explore the use of specific weights $\mathbf{w} \in \{(0.4, 0.3, 0.2, 0.1), (0.3, 0.4, 0.1, 0.2), (0.3, 0.3, 0.2, 0.2), (0.2, 0.1, 0.3, 0.4)\}$ or a combination of both, based on their data, to effectively capture features that contribute to the detection of subtypes. Therefore, for the case studies presented in this manuscript, we have chosen to adopt $\alpha = 0.01$ and $\mathbf{w} = (0.4, 0.3, 0.2, 0.1)$ as the default setting.

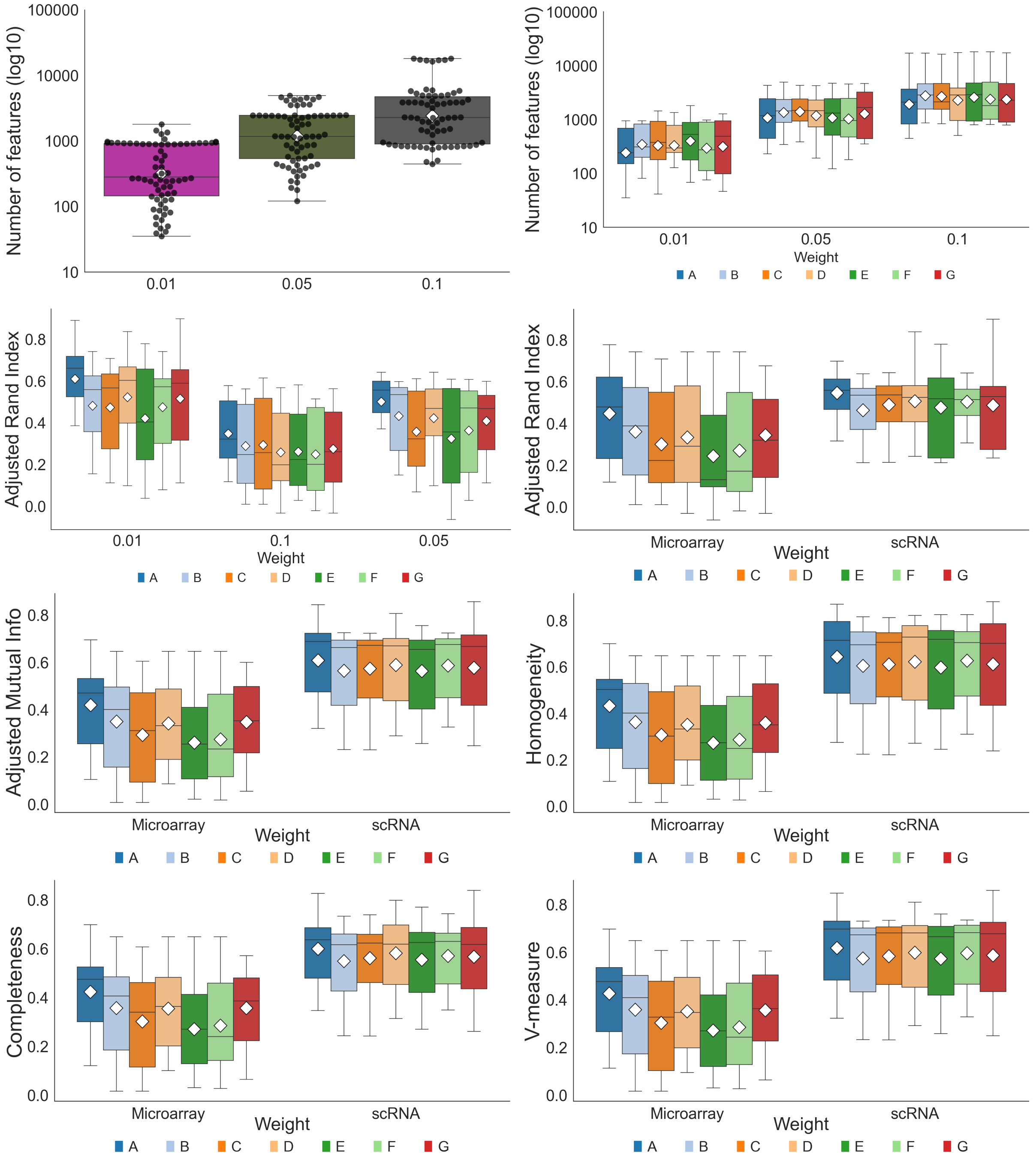


**Supplementary Figure 36:** PHet achieves the best overall performance at $\alpha = 0.01$ and $\mathbf{w} = (0.4, 0.3, 0.2, 0.1)$ across six microarray expression and four single-cell transcriptomic datasets (Supplementary Tables 1 and 2). Boxplots showing the performance of PHet and baseline methods evaluated using the number of selected features, F1 score, adjusted Rand index, adjusted mutual information, homogeneity, completeness, and V-measure. The box plots show the medians (centerlines), first and third quartiles (bounds of boxes), and 1.5× interquartile range (whiskers). A ♢ symbol represents a mean value. A: (0.4, 0.3, 0.2, 0.1), B: (0.3, 0.4, 0.1, 0.2), C: (0.3, 0.3, 0.2, 0.2), D: (0.2, 0.1, 0.3, 0.4), E: (0.1, 0.2, 0.4, 0.3), F: (0.1, 0.2, 0.3, 0.4), G: (0.2, 0.2, 0.3, 0.3).

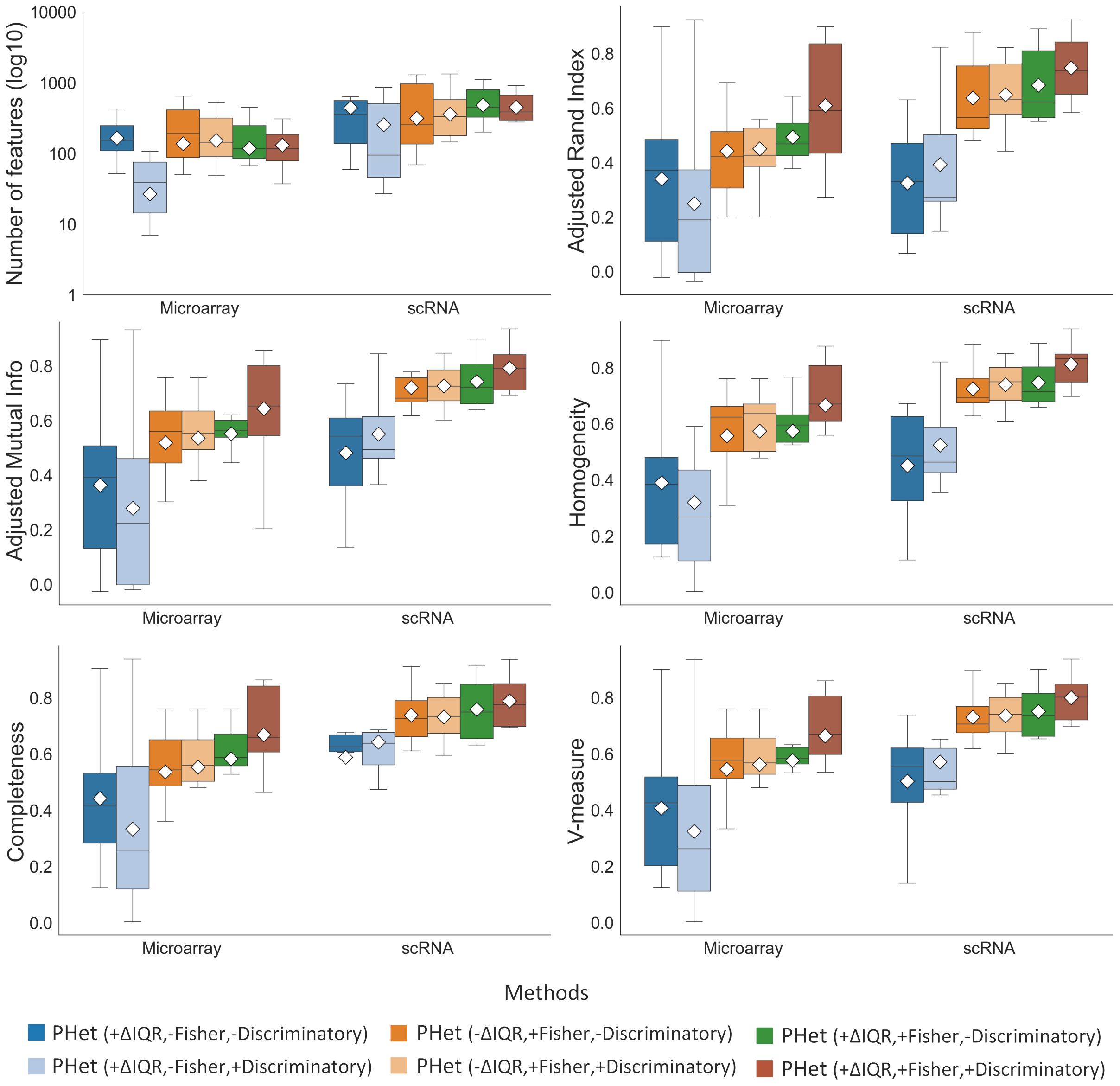


**Supplementary Figure 37:** PHet achieves the best overall performance when including the $\Delta$IQR, the Fisher's combined probability test, and the feature discriminatory power across six microarray expression and four single-cell transcriptomic datasets (Supplementary Tables 1 and 2). Boxplots showing the performance of PHet and baseline methods evaluated using the number of selected features, F1 score, adjusted Rand index, adjusted mutual information, homogeneity, completeness, and V-measure. The box plots show the medians (centerlines), first and third quartiles (bounds of boxes), and 1.5$\times$ interquartile range (whiskers). A $\Diamond$ symbol represents a mean value. A green dashed line highlights the best-performing result achieved by PHet on each dataset for each metric, while a red dashed line signifies the worst-performing result. The use of the (+) symbol indicates the addition of component(s) to PHet, while the (-) symbol signifies the removal of certain component(s) from PHet.

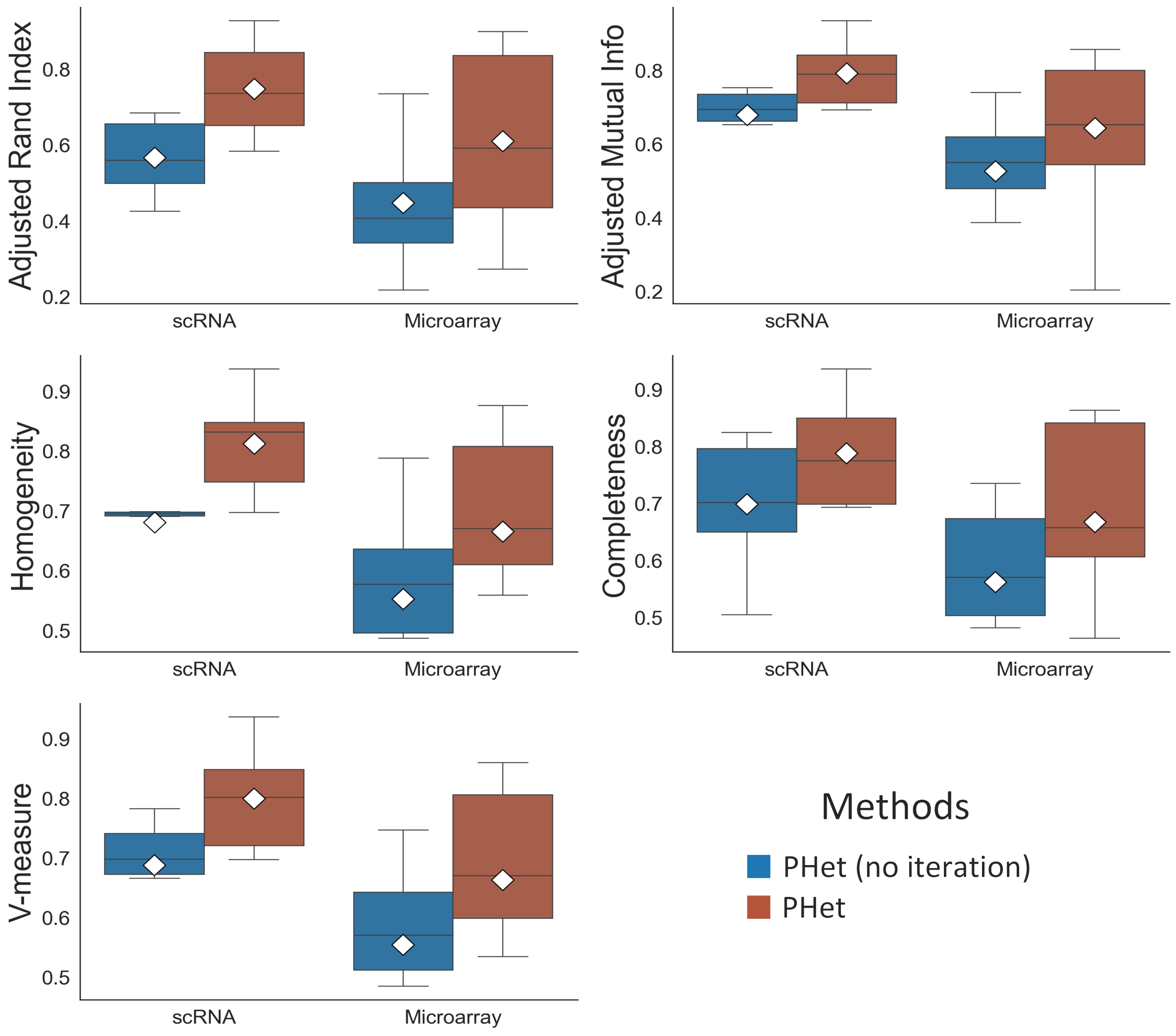


**Supplementary Figure 38:** PHet achieves the best overall performance with iterative subsampling approach across six microarray expression and four single-cell transcriptomic datasets (Supplementary Tables 1 and 2). Boxplots showing the performance of PHet and baseline methods evaluated using the number of selected features, F1 score, adjusted Rand index, adjusted mutual information, homogeneity, completeness, and V-measure. The box plots show the medians (centerlines), first and third quartiles (bounds of boxes), and 1.5× interquartile range (whiskers). A ◊ symbol represents a mean value.
